# Refined USP25/28 inhibitors with enhanced selectivity towards c-Myc driven squamous lung cancer cells

**DOI:** 10.64898/2026.03.17.712179

**Authors:** Adán Pinto-Fernández, Claire Heride, Andrew P Turnbull, Dora Pedroso, Victoria Smith, Lisa Mullee, Wojciech W Krajewski, Cameron Bell, Anthony Varca, Thomas Charlton, Dylan T Jones, Tom E McAllister, Roman Fischer, Elena Navarro-Guerrero, Daniel Ebner, Akane Kawamura, Sunkyu Kim, Dave Guerin, Tim R Hammonds, Jeffrey Kearns, Christopher Dinsmore, Neil Jones, Sara J Buhrlage, David Komander, Sylvie Urbé, Michael J Clague, Benedikt M Kessler

## Abstract

The ubiquitin specific protease 28 (USP28) is implicated in tumorigenesis by controlling the turnover of substrates including the oncogene c-MYC and the ubiquitin ligase FBW7. Here, we describe small molecule inhibitors of USP25 and USP28, leading to cancer cell cycle arrest and death. However, genetic deletion of USP25/28 does not replicate this effect. An integrated –omics approach revealed off-target effects for thienopyridine and thienopyrazine carboxamide compounds in protein translation. Chemoproteomics and biochemical analyses suggested binding of such compounds to a region near the ribosome complex polypeptide exit tunnel. Structural analysis of a USP28-inhibitor complex enabled the design of modified USP25/28 inhibitor molecules which minimized translation-related off-target effects. In distinction to earlier compounds, the optimized inhibitors were non-toxic to breast cancer cells yet retained potent anti-proliferative activity in squamous lung carcinoma cells, where USP28 is associated with disease progression. Together, our results demonstrate that refined USP25/28 inhibitors can selectively suppress tumor growth by targeting c-MYC driven pathways, offering a more precise therapeutic strategy for treating squamous lung cancers whilst minimizing undesired cytotoxicity.

## INTRODUCTION

Eliminating or inactivating oncogenes and their corresponding protein products remains a strong therapeutic strategy against cancer. KRAS mutant variants driving tumorigenesis have successfully been targeted with selective small molecules ^1,2^, but this has proven challenging for oncogenes that encode transcription factors, such as c-MYC, ETS, and ELK ^3^. Selective induction of their degradation and removal can be achieved by harnessing the ubiquitin system, either through small molecule “degraders”, heterobifunctional PROteolysis TArgeting Chimeras (PROTACs), molecular glues or by interfering with the removal of poly-ubiquitin chains from protein substrates by targeting deubiquitylating enzymes (DUBs) ^4^. Small molecule inhibitors against DUBs have recently attracted considerable attention, initially for USP7 ^5–9^, but also for other DUBs including USP25/28 ^10–13^. This interest in targeting USP28 was prompted by findings that c-MYC oncogene protein turnover is dependent on the ubiquitin E3 ligase FBW7 ^14^ and opposed by the ubiquitin specific proteases USP28 and USP36 ^15–18^. USP28 deletion and missense mutations have been proposed to affect 53BP1-dependent DNA repair pathways ^19,20^, p53 stabilisation^21^, centrosome loss and associated cell-cycle prolongation in cancer ^22–24^. Growth of several tumour types apparently depends on USP28, such as esophageal cancer ^25^, ovarian clear cell carcinoma (CCC) ^26^, gastric cancer ^27^, non-small-cell lung cancer (NSCLC) ^28^, lung squamous cell carcinoma (SCC) ^29^ and clear cell renal cell carcinoma (ccRCC) ^30^, many of which are c-Myc-dependent and p53 deficient. USP28 may also act as a tumour suppressor in melanoma ^31^, breast and liver carcinogenesis ^32^, possibly through the p53BP1-p53-p21-USP28 axis including caspase-8 ^19,23,33^, requiring wildtype p53 and preserving USP28-53BP1 interactions ^21^. Other, more tissue-specific factors, such as the squamous epithelial enriched transcription factor Δ-Np63, may be targeted by USP28 specifically in lung squamous cell cancer (LSCC), but not in the adenocarcinoma counterpart LADC ^29,34^. Several USP28 small molecule inhibitors have been described, such as AZ1 ^12^, Vismodegib (VSM) ^35^, bin-01-07-07 & derivatives ^36^, FT206 ^13^, Otilonium bromide ^11^ and CT1113 ^37^. Reduced squamous lung cancer growth *in vivo* was recapitulated with the small molecule inhibitors AZ1 ^29^, FT206 ^13^ and CT1113 ^38^, affecting c-MYC, LSD1 and Δ-Np63 levels. AZ1, VSM and FT206 all cross-react with USP25 due to its close homology with USP28 ^39^, sharing a common binding mode wedged between the finger- and α5-domains ^40–42^. This offers an additional opportunity for treatment of pancreatic cancer, in which USP25 has been implicated in progression ^43^. While the c-MYC–LSD1–ΔNp63 axis is most commonly implicated, several of these inhibitors also exhibit activity against targets beyond this pathway ^12,29,35^. Also, no marked small molecule-based selectivity towards either USP28 nor USP25 has been achieved, despite emerging molecular insights in the mechanism of USP25/28 inhibition ^40,41^ and more advanced inhibitor design ^44^.

In the present study, we compared the properties of USP25/28 small molecule inhibitors generated from different chemical scaffolds and their effects on cancer cells. The functional discrepancy between USP28 deletion and chemical inhibition, studied most prominently with thienopyridine carboxamide inhibitors, was determined to be due to off-target effects upon protein translation. Structure aided design yielded refined USP25/28 inhibitors with little or no effect on protein translation that retained their potency in selectively affecting growth of squamous lung cancer cells.

## RESULTS

### Profiling USP28 inhibitors for selectivity and cellular target engagement

The finding that USP28 plays a key role in LSCC tumour maintenance prompted us to identify small molecule inhibitors against this DUB. A small molecule discovery campaign based on the ubiquitin-rhodamine cleavable assay yielded a panel of compounds sharing a thienopyridine carboxamide chemical scaffold with inhibitory activity. These compounds were selective for USP28 and USP25 ^45,46^ when tested using the Ubiquitin-bromoethyl (Ub-C_2_Br) activity-based probe as a substrate (**Figure 1a**). The identified compounds FT206 ^13^, FT224 (**Figure 1b**), FT811 (**Figure S1a,b**, left panel) share thienopyridine / thienopyrazine carboxamide scaffolds similar to B55 and analogues ^47^. However, they represent a different chemical class from the previously described benzylic amino ethanol-based inhibitor AZ1 (**Figure S1a, b**, right panel) ^12^, Otilonium bromide ^11^ or Vismodegib ^35,41^. As shown previously for FT206, FT224 interferes with cellular USP25/28 activity-based probe HA-Ub-C_2_Br labelling (**Figure 1c**). FT224 is also selective for USP28 and USP25 at a cellular level as measured by an activity-based probe profiling mass spectrometry (ABPP-MS) assay, profiling 36 endogenous DUBs in MCF-7 breast cancer cells (**Figure 1d, e**).

**Figure 1:**
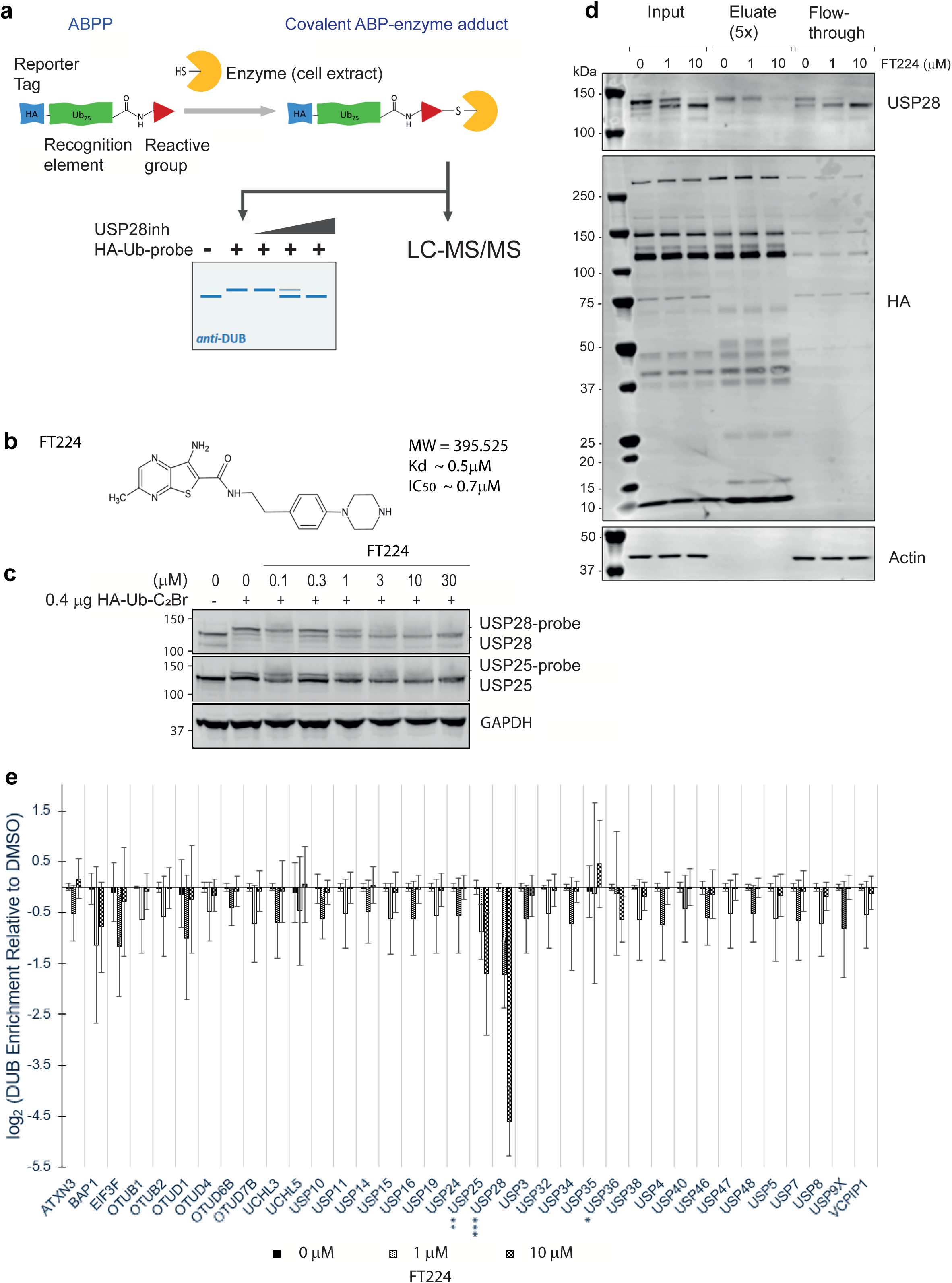
USP25/28 inhibitor FT224 cellular target engagement. **a** Schematic of activity-based protein profiling (ABPP) assay. HA (blue) -tagged ubiquitin (green) probes containing a C-terminal reactive group (red) covalently trap active cysteine protease deubiquitylating enzymes (DUBs, yellow) present in cellular extracts that can be visualised using western blotting or liquid chromatography tandem mass spectrometry (LC-MS/MS). **b** FT224 structure and chemical compound inhibitor properties. **c** FT224 potency towards USP28 and USP25 assessed by the ABPP assay. MCF-7 cell extracts (50μg) were exposed to FT224 inhibitor at the indicated concentrations for 30 min at 37°C, followed by addition of the HA-Ub-C_2_Br probe for 30 min, separation by SDS-PAGE and analysis by western blotting. **d** ABPP-MS assay controls. MCF-7 extract material (500μg) was incubated with FT224 at indicated concentrations, followed by HA-Ub-C_2_Br probe labelling as in (**c**), anti-HA immunoprecipitation. Input, eluate and flow-through material were separated by SDS-PAGE and analysed by immunoblotting. **e** ABPP-MS of the FT224 cellular DUB target engagement profile. MCF-7 cell extracts exposed to FT224 at the indicated concentrations as in (**d**) were analysed via quantitative proteomics. Shown is the differential enrichment of cellular DUBs in FT224-treated samples relative to DMSO (0 μM) control. Three technical replicates were included in the analysis. * p<0.05; ** p<0.01 and *** p<0.001, respectively.

**Table 1:**
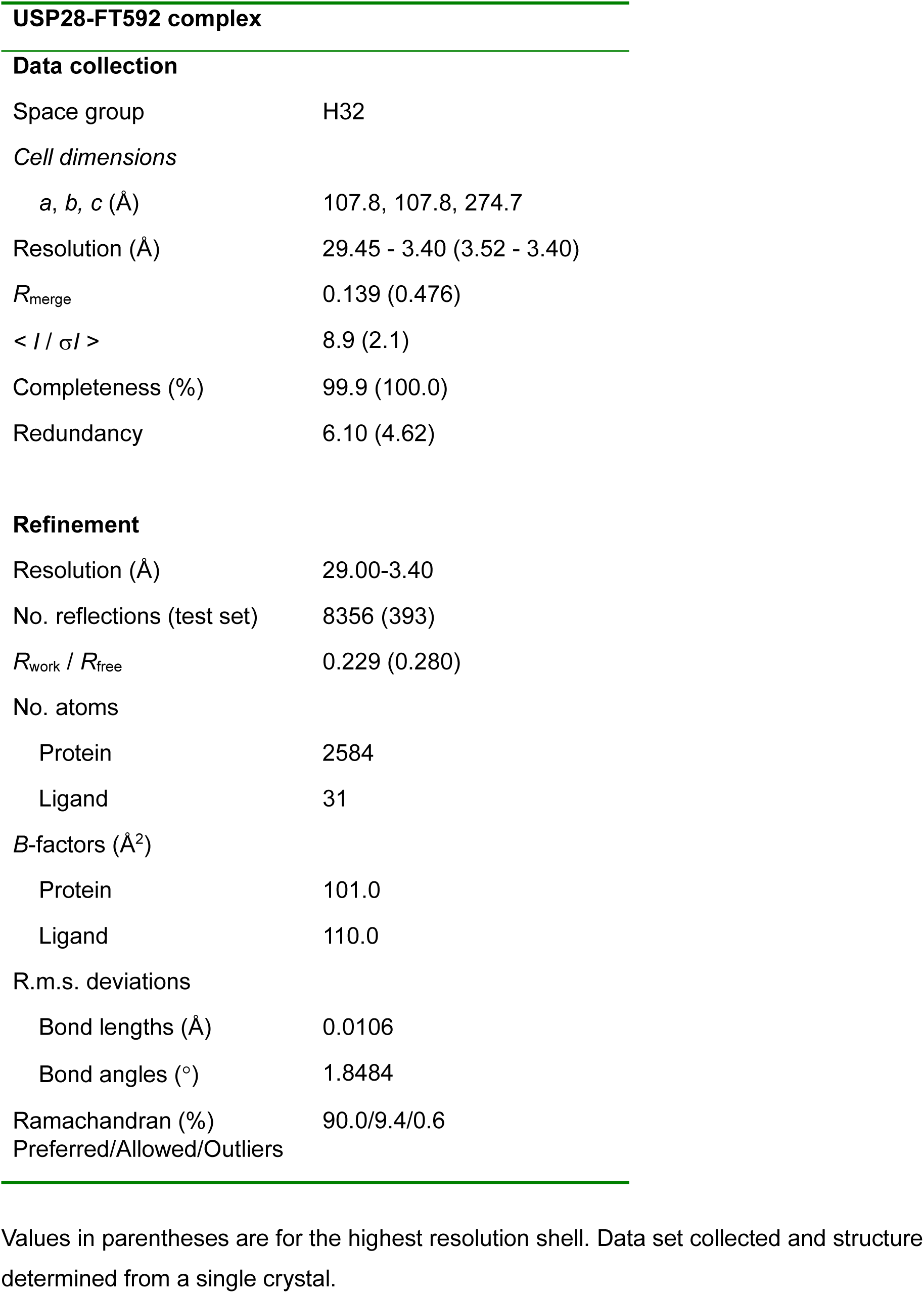
Data collection and refinement statistics. Values in parentheses are for the highest resolution shell. Data set collected and structure determined from a single crystal.

**Table 2:** Antibodies used in this study.

| Antibody name | Group | Raised in | Source | Catalog number | Concentration/time |
| --- | --- | --- | --- | --- | --- |
| Actin | Cytoskeleton | Rabbit | Sigma | A2266 | 1:2000 1hr. 1:10000 2.5 % BSA |
| Actin | Cytoskeleton | Mouse | Abcam | Ab6276 | 1:10,000, 50 mins |
| Aurora A (D3E4Q) | Cell cycle | Rabbit | Cell Signaling | (NEB) #14475 Y | 1:1000,o/n |
| Beta-actin | Cytoskeleton | Mouse | Cell Signaling | 3700 | 1:1000 o/n |
| c-Jun | Transcription Factor | Mouse | BD Transduction labor | 610326 | 1:1000 o/n |
| c-Myc | Transcription Factor | Rabbit | Cell Signaling | #5606 | 1: 1000 O/N |
| c-Myc | Transcription Factor | Rabbit | Cell Signaling | 13987 | 1:1000 o/n |
| Cyclin B1 | Cell cycle | Mouse | Abcam | 05-373 | 1:1000, 1hr or o/n |
| Cyclin D1 | Cell cycle | mouse | Santa Cruz | sc-8396 | 1:1000, O/N |
| Cyclin D1 | Cell cycle | Rabbit | Cell Signaling | 2922 | 1:1000 o/n |
| Cyclin E1 | Cell cycle | Mouse | Cell Signalling | HE12 | 1:1000, o/n |
| EEF2 | Translatoin | Rabbit | Cell Signaling | 2332 | 1:1000 o/n |
| Fbw7 | E3-Ligase etc | Rabbit | Bethyl | A301-720A | 1:1000, O/N |
| GAPDH | Metabolism | Mouse | Invitrogen | MA5-15738 | 1:1000 o/n |
| GST Tag Monoclonal Antibody (8-326) | GST-tag | Mouse | Invitrogen | MA4-004 | 1:1000 o/n |
| HA | Hemagglutinin tag | Mouse | Roche | 11666606001 | 1:2000 o/n |
| LSD1 | Chromatin | Mouse mAb | Cell Signaling | #4218 | 1:1000 o/n or 1hr |
| Notch1 (D1E11) XP | Transcription | Rabbit mAb | Cell Signaling | #3608 | 1:1000 |
| p21/Waf1/Cip | Signalling | Rabbit | Cell Signaling | 2947 | 1:1000 o/n |
| p4EBP1 (Thr 37/46) | Translatoin | Rabbit | Cell Signaling | 2855 | 1:1000 o/n |
| p53 | Transcription Factor | Mouse | Cell Signaling | 2524 | 1:1000 o/n |
| p70S6K (Thr389) | Signalling | Rabbit | Cell Signaling | 9234 | 1:1000 o/n |
| pAKT (Ser 473) | Signalling | Rabbit | Cell Signaling | 4060 | 1:1000 o/n |
| pCDK1 | Cell cycle | Rabbit | Cell Signalling | 9111 | 1:1000, o/n |
| pEEF2 (Tyr 56) | Translation | Rabbit | Cell Signaling | 2331 | 1:1000 o/n |
| PRAME | Translation | Rabbit | Cell Signaling | 45084 | 1:1000 o/n |
| PS6RP (Ser 235/236) | Signalling | Rabbit | Cell Signaling | 4858 | 1:1000 o/n |
| Puromycin | Translation | Mouse | Merck - Millipore | MABE343 | 1:1000 o/n |
| phospho-Rb | Cell cycle |  | Cell Signaling | (NEB) 9301 | 1:1000, o/n |
| RPS10 | Ribosome | Rabbit | Abcam | Ab151550 | 1:1000 o/n |
| RPS2 | Ribosome | Rabbit | Abcam | ab155961 | 1:1000 o/n |
| RPS27A | Ribosome | Rabbit | Invitrogen | PA5-141043 | 1:1000 o/n |
| RPS27L | Ribosome | Rabbit | Abcam | Ab221904 | 1:1000 o/n |
| RRBP1 | Ribosome | Mouse | Invitrogen | MA5-18289 | 1:1000 o/n |
| Streptavidin (IRDYE 800CW) | Biotin-specific | N/A | Licor Biosciences | 926-32230 | 1:1000 o/n |
| Ubiquitin | Protein turnover | Mouse | Merck - Millipore | MAB5486 | 1:1000 o/n |
| USP25 | DUB | Rabbit | Abcam | ab187156 | 1:1000 o/n |
| USP28 | DUB | Rabbit | Abcam | ab126604 | 1:1000 1h or o/n |
| USP28 | DUB | Goat | Abcam | ab130351 | 1:1000 1hr RT |
| 680RD Goat anti-Mouse IgG Secondary Antibody | Secondary Antibody | Goad | Licor Biosciences | 926-68070 | 1:20,000 |
| 800RD Goat anti-Rabbit IgG Secondary Antibody | Secondary Antibody | Goad | Licor Biosciences | 926-32211 | 1:20,000 |

### USP28 deletion and chemical inhibition alter cell cycle progression and the USP28-FBW7 network

We next explored whether USP25/28 inhibitors had any effects on cell cycle progression. This may occur via altering protein levels of shared USP28/FBW7 substrates, such as c-Myc, c-Jun, Aurora A (AURKA) and Cyclin E1. Indeed, in HCT116 colorectal cancer cells, USP25/28 chemical inhibition by FT224 for 4 and 24 hrs affected levels of c-Myc, Cyclin E1 and, to a lesser degree, Aurora A. Strikingly the strongest effect was seen on Cyclin D1 (**Figures 2a, c, S2, S3a**). C-Myc and Cyclin E1 protein levels were elevated in FBW7 knockout cells as reported previously ^48,49^, but remained sensitive to FT224, as did Cyclin D1 and Aurora A (**Figure S3a**). An overall decrease in S-phase progression was noted upon 24-hour exposure to FT224 (**Figure 2b**), associated with altered protein abundance of Cyclin D1, Cyclin E1 and a decrease in CDK1 phosphorylation (**Figure 2c**). Cyclin D1 mRNA levels were unaffected, suggesting that FT224’s mechanism of action is at the posttranscriptional level (**Figure S3b**). These effects were phenocopied by USP28 siRNA knockdown to some extent as demonstrated also by reduced levels of Rb S795 phosphorylation and Cyclin D1 protein levels (**Figure 2d**). We observed reduced c-Myc and Cyclin D1 protein levels in a time- and concentration-dependent manner, that could partially be rescued by combining USP25/28 inhibition with proteasomal blockade using epoxomicin or interfering with Cullin E3 ligase mediated ubiquitylation with the NEDDylation inhibitor MLN4924 in HCT116 colorectal (**Figure S3c**) and MCF-7 human breast cancer cells (**Figure S3d**), suggesting ubiquitin mediated alterations in the USP28-c-Myc-FBW7 network.

**Figure 2:**
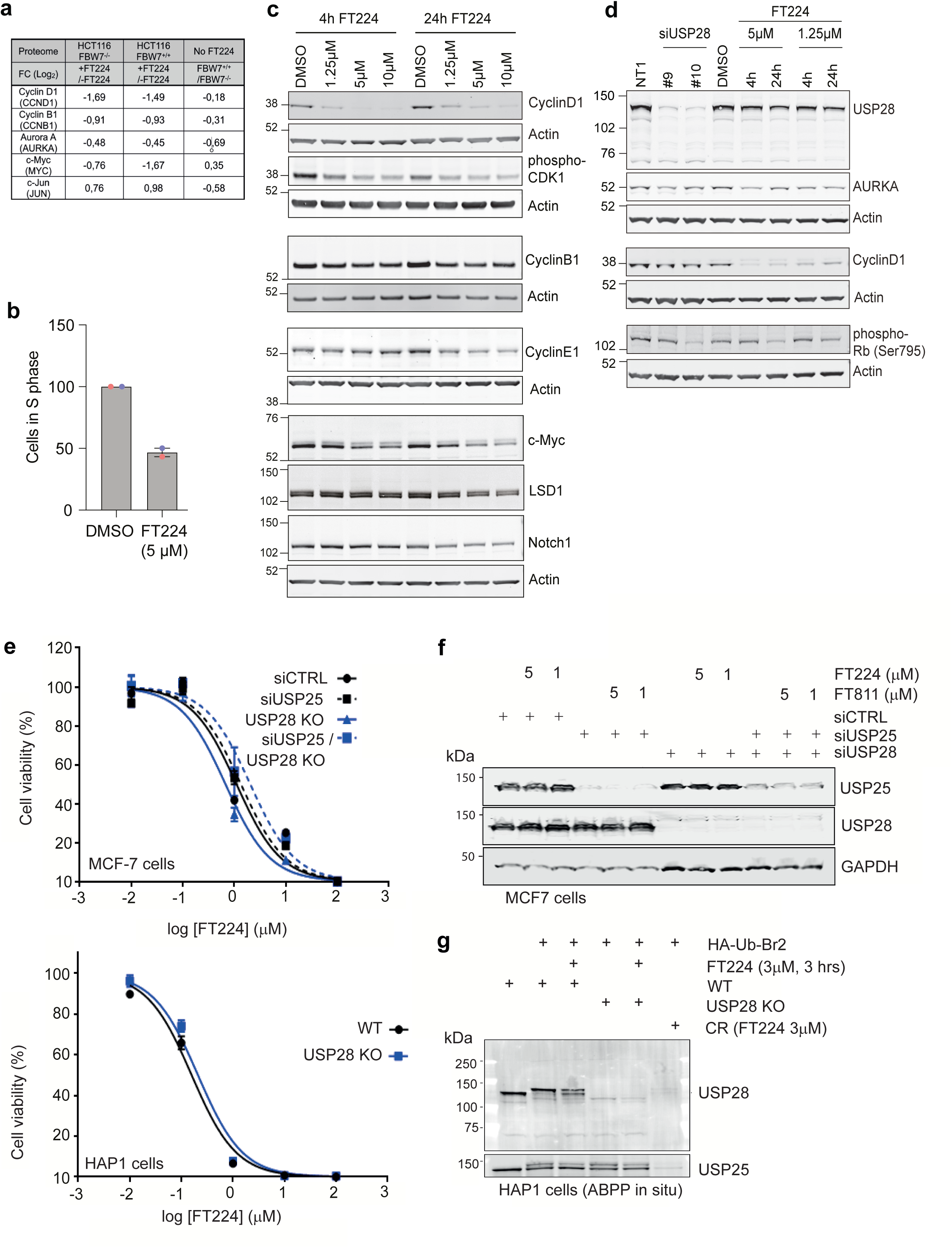
FT224 affects cell cycle progression independent of USP25/28. **a** Relative abundance data (Log_2_ Fold change) for select proteins extracted from a SILAC-based quantitative total proteome analysis of HCT116 FBW7 wild-type (+/+) or knockout (-/-) cells treated for 4 hrs with FT224 (5 µM) or vehicle (DMSO), prior to SDS-lysis. CCND1, CCNB1 (mean of n=2 independent experiments). See also Figure S2 for full dataset. **b** Treatment for 24 hrs with FT224 (5 µM) decreases the number of MCF-7 cells in S-phase. Cell cycle distribution was analysed by flow cytometry using propidium iodide staining. Shown are data from two independent experiments. Error bars show the range. **c** FT224 effects on known USP28 substrates (c-Myc, Cyclin E1, LSD1, Notch) and cell cycle regulators in MCF7 cells after 4 and 24 hrs. **d** USP28 siRNA knockdown (KD) using two independent siRNAs (#9, #10) or FT224 treatment for 4 and 24 hrs affect cell cycle regulators. Representative western blots are shown. **e** MCF-7 cell viability upon USP28 and/or USP25 knockdown (upper panel), and HAP1 cell viability upon USP28 KO (lower panel) exposed to FT224 for 72 hrs inhibitors at the indicated concentrations. **f** USP25/28 knockdown efficiencies in MCF-7 cells and effects of inhibitor exposure for 72 hrs. MCF-7 cell extracts were separated by SDS-PAGE and analysed by immunoblotting. **g** USP25/28 knockdown efficiencies in HAP1 cells exposed to the activity-based probe HA-Ub-Br_2_ and effects of FT224 inhibitor at 3μM concentration incubated for 3 or 24 hrs, respectively. MCF-7 cell extracts were separated by SDS-PAGE and analysed by immunoblotting. CR: compound resistant HAP1 cells.

### Thienopyrazine carboxamide inhibitor affects cell viability independently of USP25/28

To test the correlation between USP28 chemical inhibition and genetic deletion mediated effects on the cell cycle, we knocked out (KO) the *USP28* gene in a range of cell lines of different cancer indications such as breast cancer (MCF-7), chronic myeloid leukaemia (HAP1) and colorectal cancer (HCT116). This resulted in complete or partial reduction of USP28 protein expression (**Figure 2, S4, S5**). Remarkably, cell proliferation of MCF-7 and HAP1 WT and USP28 KO cells were indistinguishable, irrespective of USP25 depletion by siRNA, in contrast to chemical inhibition with FT224 (**Figure 2e-g**). These results indicated the existence of potential off-target effects of this thienopyridine carboxamide-based USP25/28 inhibitor. To confirm and explore this further, individual USP28 KO HCT116 colorectal cancer cell clones were generated (**Figure S4a**). Suppression levels of USP28 expression varied between USP28 KO clones (**Figure S4b**), but a strong positive correlation could be observed between the protein levels of USP28 and Cyclin D1 (**Figure S4c-e**). Analysis of USP28 KO clones 98-3 and 99-2 showed a reduction of Cyclin D1 levels as compared to the WT control clones 98-17 and 99-9, respectively, which was further decreased by adding inhibitors FT224 or FT811 (**Figure S5a, b**), confirming the polypharmacological nature of these inhibitors. Taken together, we noted clear discrepancies between USP25/28 chemical inhibition and deletion on cell viability and substrate abundances along the USP28-FBW7 axis.

### Mapping USP25/28 thienopyrazine carboxamide inhibitor off-target binding in the ribosome complex

To explore off-targets of these compounds, we generated a biotinylated USP25/28 inhibitor FT224 derivative, FT639 (**Figure 3a**) that retained USP28 inhibitory activity *in vitro* and enriched USP25/28 in pulldown assays (**Figure 3b**). Chemoproteomics experiments were optimised using bead types and various stringencies of buffer conditions (**Figure 3c**). Increasing amounts of the biotinylated FT639 inhibitor probe revealed a dose-dependent enrichment of components of the translation machinery including RPS27L, EEF2, DHX16 and RRBP1 (**Figures 3d** – left, middle and right panels). Most strikingly, RPS27A and RPS27L, enriched with FT639 as identified by quantitative MS, were confirmed by immunoblotting (**Figure 3e, f**). Some of these interactors are located near the ribosome mRNA channel, reminiscent to the binding region of cycloheximide (CHX) near the tRNA E-site of the ribosomal 60S subunit ^50^. Combined, our data suggest that FT224 does inhibit USP25/28 but also has an off-target in the ribosome complex, possibly leading to an interference with the formation of nascent polypeptide chains **(Figure 3g**).

**Figure 3:**
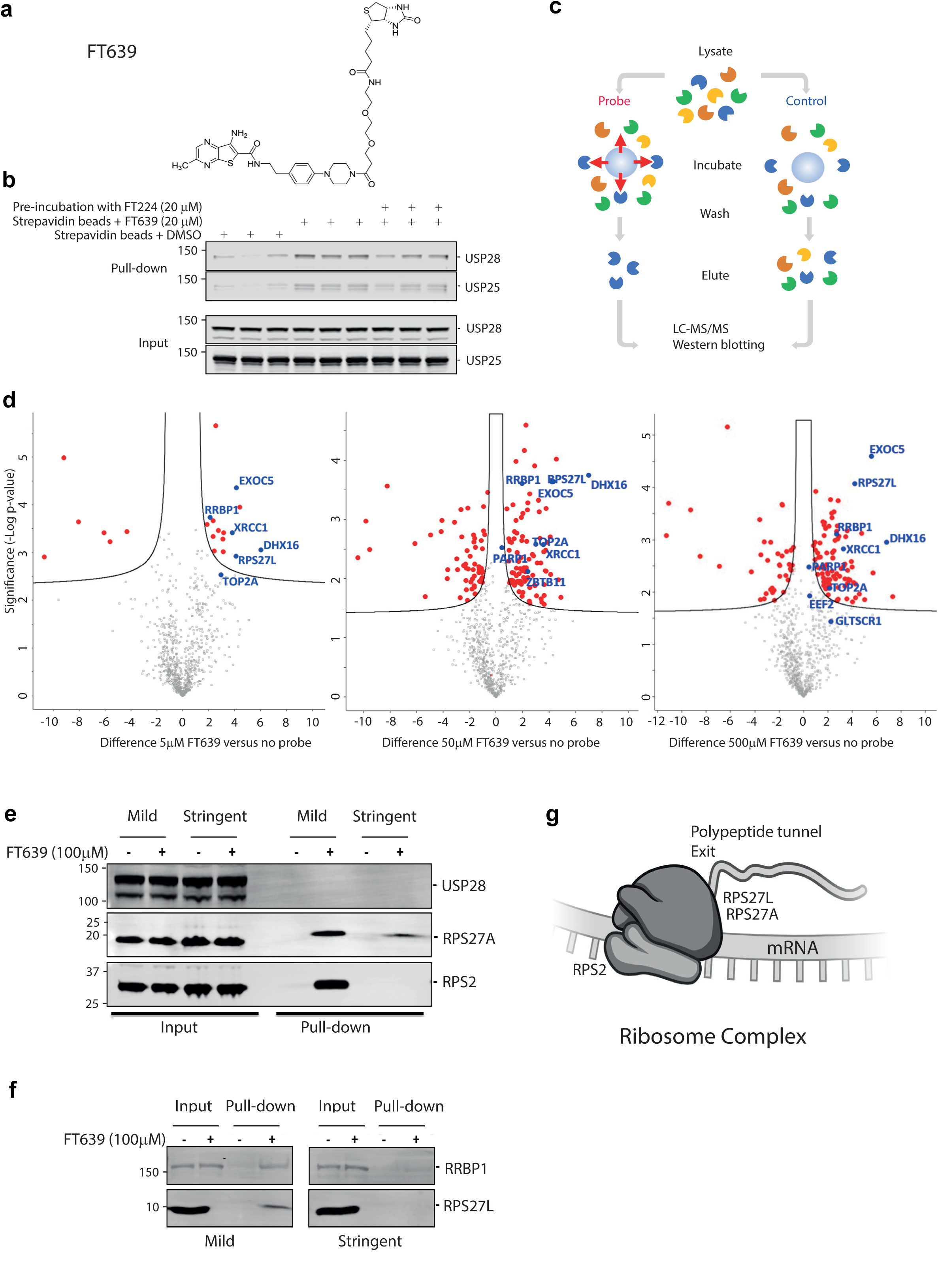
FT639 Chemoproteomics reveals the USP25/28 inhibitor interactome. **a** FT639 structure (biotinylated FT224 inhibitor). **b** USP25/28 pulldown with FT639. HAP1 cells, pre-treated with 20 μM FT224 or DMSO control for 30 min, were incubated with 20 μM FT639 or DMSO for 3 hrs and subsequently with Streptavidin beads for 2 hrs. Enriched material was separated by SDS-PAGE and analysed by immunoblotting. Three technical repeats are shown. **c** Scheme showing the chemoproteomics strategy for target discovery using biotinylated USP25/28 inhibitor FT639 (“probe”). **d** FT639 enriched cellular interactomes. Volcano plots are shown for FT639 pulldowns versus no probe control analysed by quantitative LC-MS/MS, displayed in log_2_ format (difference). Left panel: FT639 (5 μM) interactome; middle panel: FT639 (50 μM) interactome; right panel: FT639 (500 mM) interactome in HAP1 cells. Proteins that are enriched in either FT639 or control samples above the significance threshold (S0=0.05) are highlighted as dots in red or blue for labelled proteins. **e** RPS27A and to a lesser extent RPS2 co-precipitate with FT639 as analysed by FT639 enrichment, SDS-PAGE and immunoblotting. **f** RPS27L and to a lesser extent RRBP1 co-precipitate with FT639 as analysed by FT639 enrichment, SDS-PAGE and immunoblotting. **g** FT639/FT224 interaction with the ribosome complex. This may occur in a similar way than translation inhibitors that interfere with mRNA binding or translation elongation, blocking the release of nascent polypeptide chain at the exit site of the ribosome ^50^ Illustration created using www.bioart.niaid.nih.gov.

### USP25/28 thienopyrazine carboxamide inhibitors affect protein translation

To test this, we performed orthogonal assays to probe the inhibitor’s effects on protein synthesis (translation activity) in cellular and biochemical assays (**Figure 4**). First, a SUnSET assay ^51^, measuring the synthesis of nascent polypeptide chains by capping with puromycin, revealed a strong inhibitory effect of FT811 in a dose-dependent manner (**Figure 4a**), consistent with USP28 inhibitor’s effect on reducing levels of short-lived proteins, not rescuable by concomitant addition of proteasome inhibitor (**Figure 4b**). Second, we conducted the O-propargyl-puromycin (OPP) assay to visualise and quantitate nascent polypeptides in cells using fluorescence microscopy. We compared USP28 inhibitors FT206 ^13^ (thienopyridine), FT224 and FT811 (both thienopyrazines) to the USP7 inhibitor FT671 ^8^ as well as CHX and bortezomib (Bz) as positive and negative controls, respectively (**Figure 4c-e**). Strikingly, USP28 inhibitors affected protein translation in a similar manner to cycloheximide (CHX) (**Figure 4c-e**). Finally, we performed an *in vitro* assay using purified rabbit reticulocyte lysates containing the machinery needed to translate exogenous mRNA only. Translation of Luciferase mRNA was inhibited by FT224 in a dose dependent manner (**Figure S6a, b**). In contrast, FT671, a USP7 inhibitor, and AZ1 (FT214), another USP25/28 inhibitor based on a completely different chemical scaffold ^12^, did not show any effect in this assay (**Figure S6a, b**), further suggesting that thienopyridine-based USP25/28 inhibitors bind inside the translation apparatus. This is both scaffold selective and independent of USP28 as HAP1 wildtype and HAP1 USP28 KO cells revealed no difference in *de novo* protein synthesis as assayed by puromycin incorporation (**Figure S6c-e**). Consistent with a compound-dependent effect on protein translation, we observed activation of mTOR downstream signalling factors regulating protein translation such as p70S6K and S6RP (**Figure 4f**). A consequence of blocked protein translation is intracellular depletion of ubiquitin, which has been associated with cellular toxicity and aberrant translation ^51^. In line with this, we observed a dose-dependent depletion of cellular poly-ubiquitin chain material by treatment with FT224 (**Figure S6f**). Taken together, we conclude that both thienopyrazide and thienopyridine carboxamide based inhibitors may have an off-target effect that decreases protein translation.

**Figure 4:**
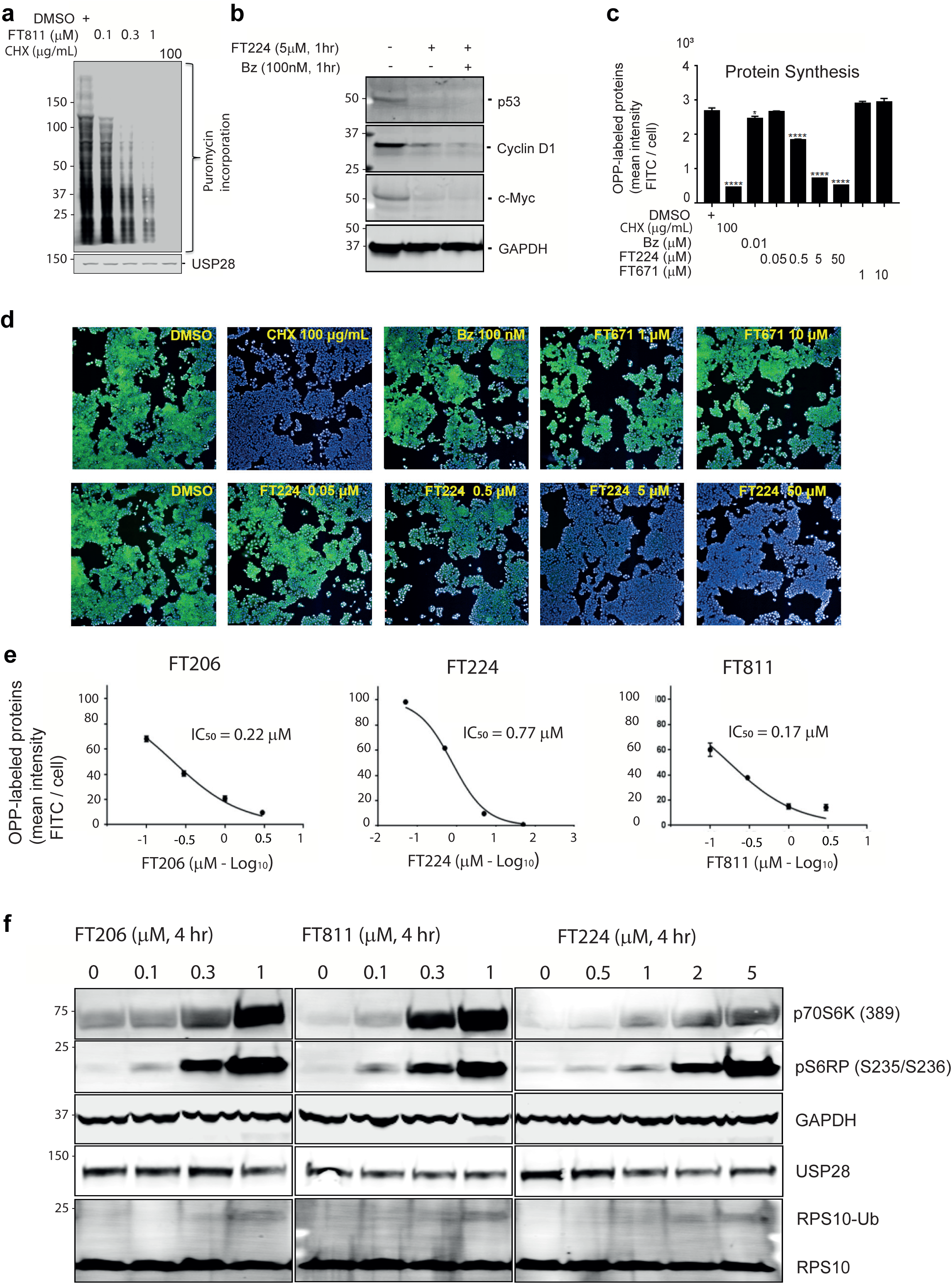
USP28 Inhibitor off-target effects on protein translation. **a** Nascent protein synthesis in HAP1 cells affected by 4 hrs exposure to FT811 inhibitor or CHX at the indicated concentrations using the SunSet assay. **b** Short-lived proteins affected by treatment of HAP1 cells with FT224 for 1 hr with or without proteasome inhibitor Bortezomib (Bz). **c** FT224 compound exposure for 4 hrs affects protein synthesis in the OPP assay in HAP1 cells. Quantitation of data shown in panel **d** from three biological replicates. Error bars show mean ± SEM. P-value style is GP: 0.1234 (ns), 0.0332 (*), 0.0021 (**), 0.0002 (***) and 0.0001 (****), respectively. **d** Thienopyridine carboxamide inhibitors FT224, affect nascent protein synthesis in HAP1 cells as measured by the OPP assay and assessed by microscopy. **e** IC_50_ values of FT224, FT206 and FT811 in the OPP assay measured in HAP1 cells exposed for 4 hrs. **f** Proteins associated with translation and ribosomal stalling affected by FT224, FT206 and FT811 in a dose dependent manner as assayed in HAP1 cells.

### USP28 enzyme-inhibitor structures reveal conserved features for on-target activity

To understand the molecular basis for the poly-pharmacology of these compounds, a crystal structure of the insert-deleted apo form of the human USP28 catalytic domain was obtained and soaked with a thienopyrimidine-based inhibitor, FT592 (**Figure 5, S7a**), an earlier chemical precursor of FT224/FT811 (**Figure 1b**). The USP28-FT592 complex highlights the molecular basis of compound selectivity. FT592 resides in a cleft formed by α1, α2, α5 and α6 of the thumb subdomain, and β5, β8, β12, β13 and β14 of the palm subdomain (**Figure 5a**, **5b**). The USP28-FT592 complex superimposes on apo form insert-deleted USP28 (PDB code 6HEH) with an r.m.s.d. of 1.02 Å (Cα atoms; 314 residues aligned; **Figure S7b**). A rotation of ∼3.8° coupled with a translation of ∼2 Å (pivoted about residue L273), displaces α5 in the USP28- FT592 complex compared with the apo form structure, expanding the pocket in which FT592 resides (**Figure S7b, c**). The thienopyridine moiety sits in a pocket flanked by S179, L180, H261, L264, F292, Q315, T364, and M646, and directly hydrogen bonds to the side chain amide of Q315 (**Figure 5b**). In addition, the central amide moiety of FT592 hydrogen bonds to the side chain carboxyl of E366. The imidazole substituent is positioned in the channel that usually accommodates the ubiquitin substrate C-terminal tail. Compared with the ubiquitin-propargylamide (UbPA)-bound complex of insert-deleted USP28 (PDB code 6HEI), there is a steric clash between the imidazole substituent and the side chain of L73 of the UbPA substrate, preventing the substrate from binding and being cleaved (**Figure 5c**, **5d**). The binding mode of FT592 is analogous to previously reported inhibitors including FT206, AZ1, and Vismodegib, which wedge between α5 and the palm subdomain preventing transition to a substrate-binding competent state ^11,40,42^ (**Figure S7d, e, f-g**).

**Figure 5:**
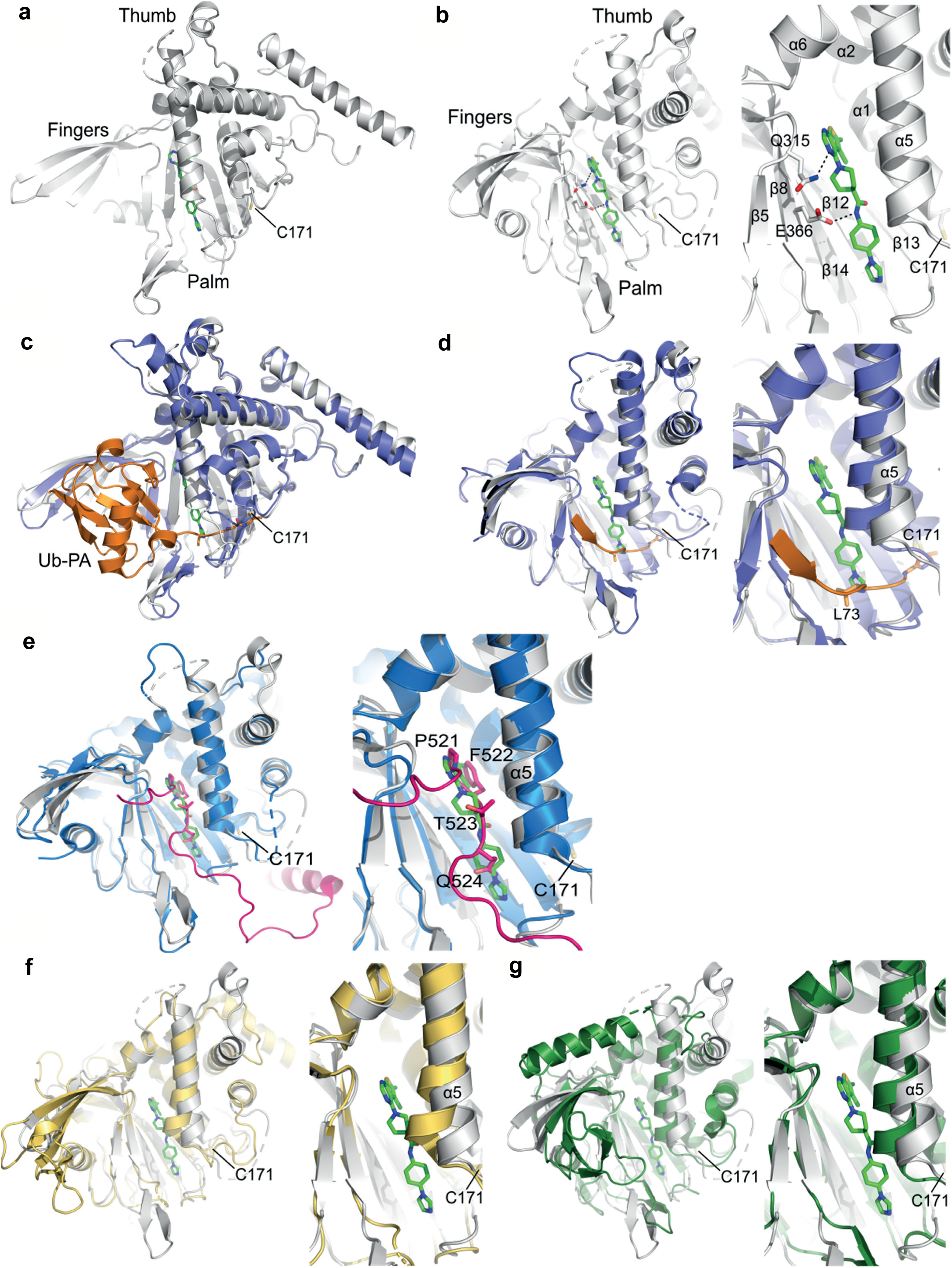
USP28-FT592 complex structure and molecular basis for selective inhibition. **a** Structure of human USP28 in complex with FT592. FT592 is shown as a stick representation with carbon atoms colored green. The thumb, palm and fingers subdomains of the catalytic domain and catalytic cysteine, C171, are highlighted. **b** Left: view in panel **a** rotated on the y-axis by 50°; right: close-up view of the FT592 binding site highlighting secondary structural elements forming the binding pocket, key residues, and hydrogen-bonding interactions represented as dotted lines. **c** Superposition of human USP28 in complex with Ub-PA (USP28 colored violet, Ub-PA colored orange; PDB code 6HEI) on the USP28-FT592 complex structure colored grey. FT592 is shown as a stick representation with carbon atoms colored green. The catalytic cysteine, C171, is highlighted. **d** Left: view in panel **c** rotated on the y-axis by 50°; right: close-up view of the FT592 binding site. The C-terminal tail of Ub-PA is shown in orange. The imidazole moiety of FT592 sterically clashes with the position of L73 in the C-terminal tail of Ub-PA, thereby preventing Ub binding and isopeptide bond cleavage. **e** Left: superposition of human USP25 colored blue and the autoinhibitory motif (AIM) in the tetramer colored magenta (PDB code 6HEL) on the structure of the human USP28-FT592 complex colored grey with FT592 shown as a stick representation and carbon atoms colored green; right: close-up view of the FT592 binding site. Residues in AIM including the Pro-Phe motif (P521-F522) are shown as stick representations and labelled. FT592 sterically clashes with AIM in the autoinhibited USP25 tetramer. **f** Left: superposition of apo form human USP7 colored gold (PDB code 1NB8) on the structure of the USP28-FT592 complex colored grey with FT592 shown as a stick representation and carbon atoms colored green; right: close-up view of the FT592 binding site highlighting the conformation of α5 in USP28 compared with α5 in USP7, which is kinked and reduces the volume of the pocket in USP7. **g** Left: superposition of human apo form USP8 colored dark green (PDB code: 2GFO) on the structure of human USP28 in complex with FT206 colored grey with FT592 shown as a stick representation with carbon atoms colored green; right: close-up view of the FT592 binding pocket highlighting the conformation of α5 in USP28 compared with USP8 in which the helix is pivoted inwards reducing the volume of the pocket. Figure items were prepared using PyMOL (The PyMOL Molecular Graphics System, Version 2.5.8, Schrödinger, LLC.).

A hallmark unique to USP25 and USP28 is that they both form homodimers, with USP25 additionally forming homotetramers via an autoinhibitory element from a neighbouring molecule ^39,52,53^. The USP28-FT592 complex, and the catalytic domain of USP25 (PDB code 6HEL) superimpose with an r.m.s.d. of 1.27 Å (Cα atoms; 299 residues aligned; **Figure 5e**). This places FT592 in the autoinhibitory motif (AIM) binding pocket of the catalytic domain of USP25, which is responsible for autoinhibition by tetramerization of USP25. There are 21 residues residing within 5 Å of FT592 in the USP28-FT592 complex (V176, S179, L180, V256, S257, T260, H261, L264, F292, G314, Q315, T364, E366, F370, L590, H592, A596, Y601, Y643, C644, and M646). These residues are fully conserved in human USP25, explaining why FT592 also inhibits USP25 catalytic activity. The 14-mer AIM is highly conserved in USP25 amongst mammalian species and centres on a Pro-Phe motif (P521-F522), whereas the equivalent sequence in USP28 lacks conservation ^39^. Superposition of USP28-FT592 on the autoinhibited USP25 tetramer reveals that the Pro-Phe motif of the adjacent dimer overlays with the position of the thienopyridine moiety in the USP28-FT592 complex (**Figure 5e**). In contrast, the USP7 inhibitors, GNE6640 (PDB code 5UQV) and GNE6776 (PDB code 5UQX), are largely solvent exposed and lie between α5 and α6 at the interface between the palm, fingers, and thumb subdomains in the catalytic domain of USP7, in a pocket that partially overlaps with the FT592 pocket in USP28 (**Figure S7h-i**). The phenol substituent of GNE6640 and GNE6776 overlays with P521 of the Pro-Phe motif in USP25. The region corresponding to the AIM pocket in USP25/28 appears larger in size as compared with USP7 (**Figure 5f**) and USP8 (**Figure 5g**), as the residues surrounding it have in many cases sterically less demanding side chains (e.g., T364 in USP25/USP28 vs. H403 in USP7 or C644 in USP25/USP28 vs. M515 in USP7 and in many other DUBs), which explains the unique selectivity of these inhibitors for USP25 and USP28.

### Optimising different USP25/28 inhibitor chemical scaffolds to remove off-target effect on translation

Structural information allowed the exploration of a systematic structure activity relationship (SAR) to preserve critical elements for selective USP25/28 inhibition and vary other parts of small molecules that may underlay the observed effects on protein translation. To this end, a panel of different molecule derivatives encompassing various chemical scaffolds including thienopyridine carboxamides and pyridothenopyridines, (pyrido[3’,2’:4,5]thieno[3,2-d]pyrimidines), but also a variety of different chemical moieties, were generated and tested for USP28 enzyme inhibition (**Figure S8-10**), demonstrating potency and USP25/28 selectivity in an ABPP assay ^36,47,54,55^. To test their potential off-target activity on protein translation, the OPP assay was used in MCF-7 cells. We found that a subset of compounds with structural differences to the FT compounds, including bin-07-07 and AV-24-34, showed much reduced or no inhibition of protein synthesis as compared to FT224 or FT811 (**Figure 6a-c**), correlating with structural features that preserved USP25/28 inhibition capacity (**Figure S11**). Confirming our previous observations, cell viability measurements of MCF-7 breast cancer cells correlated with the compound’s potency to inhibit protein translation (**Figure 6d, S8**). This confirms evidence for ribosome off-targets that were revealed also in complementary screening assays (**Figures S12, 13**), mainly affecting short-lived proteins (**Figure S14**), cell growth dynamics (**Figure S15**) and mTOR activation due to altered protein translation (**Figure S16**).

**Figure 6:**
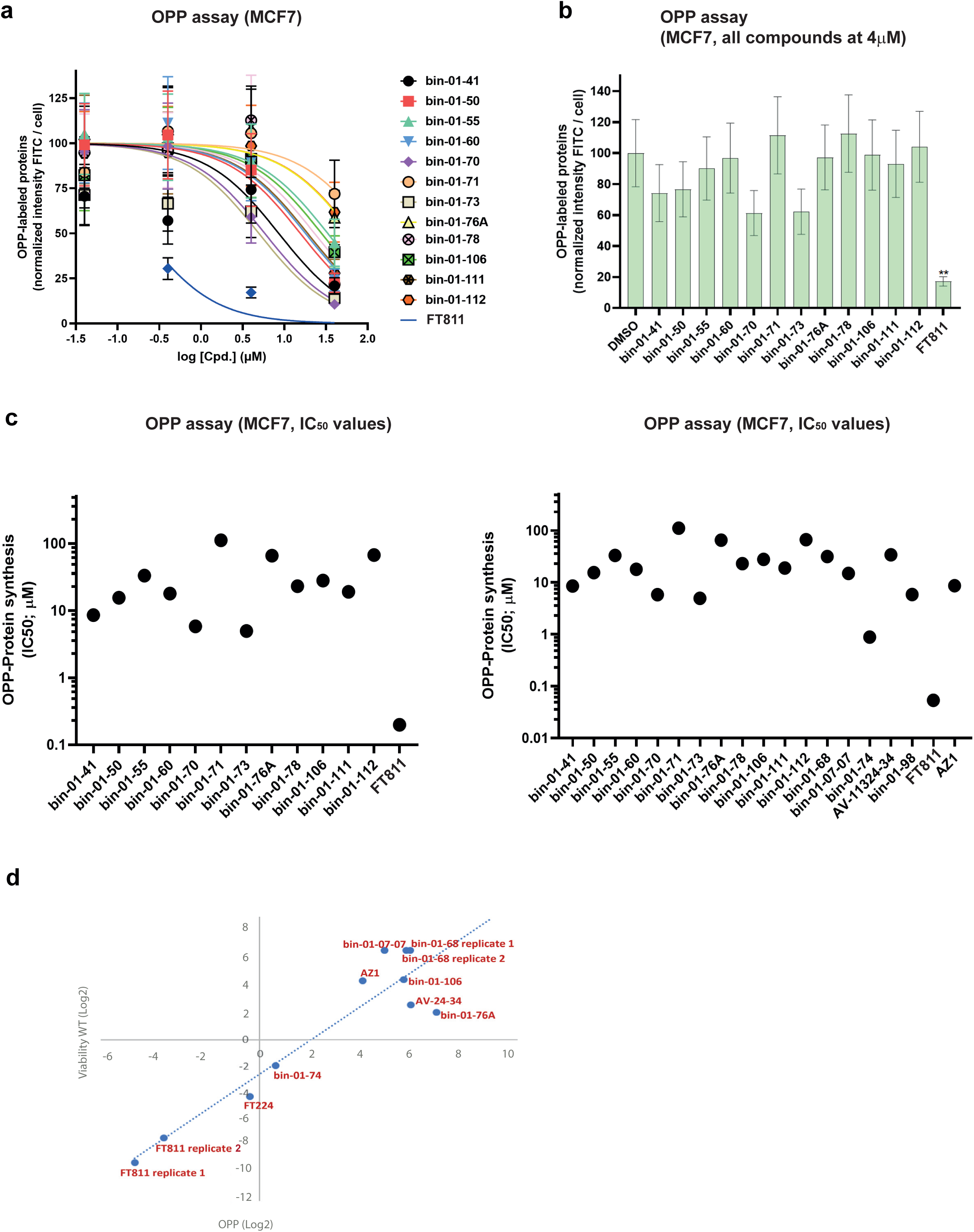
USP25/28 inhibitors with and without off-target interference with protein synthesis. **a** O-propargyl-puromycin (OPP) assay showing dose-dependent effects on protein synthesis for a panel of USP28 inhibitor compounds added to MCF-7 cells for 30 min at the indicated concentration prior to adding OPP for 1 hr. N=3 experiments, error bars show mean ± SEM. **b** OPP assay showing effects on protein synthesis in MCF-7 cells exposed for 30 min to a panel of USP28 inhibitors at 4 μM prior to adding OPP for 1 hr. N=3 experiments, error bars show mean ± SEM. **c** OPP assay compound IC_50_ values (based on panel **a**). **d** Correlation of compound OPP activity and effect on cell viability for the indicated compounds in MCF-7 cells exposed to inhibitors for 30 min followed by the addition of OPP.

### Improved selectivity towards squamous lung cancer cell susceptibility due to reduced off-target inhibition

USP28 dependence is linked to cancer types driven by c-Myc and p63, such as squamous lung cancer ^13,29,56^. To test this, we exposed wildtype MCF-7 breast cancer cells as well as H-520 squamous lung cancer cells to refined USP25/28 inhibitors with a reduced or lack of potency in protein translation inhibition, namely bin-07-07 and AV-24-34 (**Figure 7**). As controls, we used FT224 and bin-01-74, two inhibitors with similar structures and clearly established translation inhibition profiles (**Figure S11**). FT224 and bin-01-74 were, as expected, the most potent cell viability inhibitors in both cell lines (**Figure 7a, b**). In contrast, bin-07-07 and AV-24-34, relatively ineffective when exposed to MCF-7 breast cancer cells (**Figure 7c, e**), showed an enhanced cell viability profile for H-520 squamous lung cancer cells (**Figure 7d, f**). Half-Maximal Inhibitory Concentration values were calculated based on independent measurements, revealing the selectivity of the refined inhibitors bin-07-07 and AV-24-34 to affect squamous lung cancer H520 (IC_50_ 4.1μM / 50.7μM) over breast cancer MCF-7 cells (IC_50_ > 500μM for both compounds), a trait not observed with the original compound FT224 and bin-01-74 (**Figure 7g**). We conclude that this apparent selectivity may be due to a reduction or loss of off-target activity in translation (**Figure 7h**).

**Figure 7:**
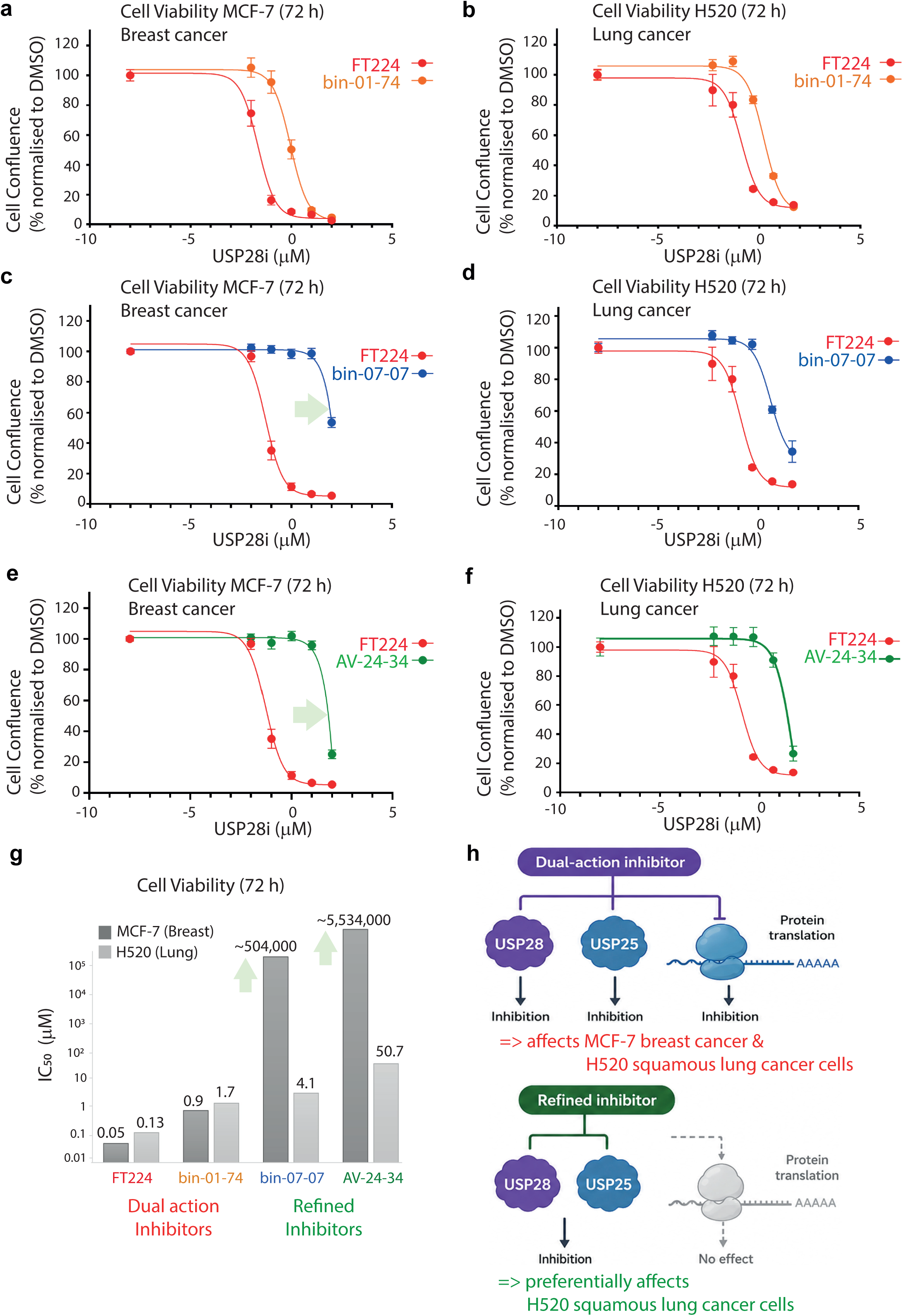
Refined inhibitors preferentially affect squamous lung cancer cell susceptibility. **a** MCF-7 breast cancer cell viability assessed after 72 hrs exposure to inhibitors FT224 or bin-01-74 at different concentrations. Two independent experiments with quadruplicate measurements are shown. Data are presented as mean ± SEM. **b** H520 squamous lung cancer cell viability assessed after 72 hrs exposure of inhibitors FT224 or bin-01-74 at different concentrations. Two independent experiments with quadruplicate measurements are shown. Data are presented as mean ± SEM. **c** MCF-7 breast cancer cell viability assessed after 72 hrs exposure of inhibitors FT224 or bin-07-07 at different concentrations. Two independent experiments with quadruplicate measurements are shown. Data are presented as mean ± SEM. **d** H520 squamous lung cancer cell viability assessed after 72 hrs exposure of inhibitors FT224 or bin-07-07 at different concentrations. Two independent experiments with quadruplicate measurements are shown. Data are presented as mean ± SEM. **e** MCF-7 breast cancer cell viability assessed after 72 hrs exposure of inhibitors FT224 or AV-24-34 at different concentrations. Two independent experiments with quadruplicate measurements are shown. Data are presented as mean ± SEM. **f** H520 squamous lung cancer cell viability assessed after 72 hrs exposure of inhibitors FT224 or AV-24-34 at different concentrations. Two independent experiments with quadruplicate measurements are shown. Data are presented as mean ± SEM. **g** IC_50_ values shown for **a-f.** Data were obtained from two biological replicates per condition/concentration, with four IncuCyte images analysed per biological replicate at each time point as technical quadruplicates. Data are presented as mean ± SEM, and IC₅₀ values were calculated using a three-parameter nonlinear regression fit using Prism software (Supplementary Information). **h** Dual action inhibitors FT224 (red) and bin-01-74 (orange) versus refined inhibitors bin-07-07 (blue) and AV-24-34 (green) mechanism of action. Illustration created with ChatGPT Edu University of Oxford (GPT-5.5).

## DISCUSSION

Lung squamous cell carcinoma (LSCC) development is characterised by i) mutational loss of p53 and RB1 tumour suppressors, ii) lineage-specific transcription factors, also referred to as lineage oncogenes ASCL1, NEUROD1, POU2F3, inducing a neuro-endocrine expression programme, iii) inhibition of NOTCH signalling ^57^ and iv) overexpression of members of the myc family, such as c-MYC, MYCL or MYCN ^58^. USP28 small molecule inhibition/deletion affects NOTCH-1, c-MYC, LSD1 and Δ-Np63 levels ^29,59^, reflecting common targets for squamous cell carcinomas ^34^. Thus, targeting USP28 represents an attractive therapeutic strategy against LSCC and potentially other c-MYC-dependent p53 deficient cancer types.

Multiple chemical scaffolds have led to the discovery of USP25/28 inhibitors. This includes thienopyridine/thienopyrazine carboxamides ^45^, benzyl ethanolamines ^12^ and hybrid scaffolds ^54^, with inhibition potencies between high nM to the mid μM range. CT1113 ^60,61^, which shows a clear similarity to the previously reported thienopyridine carboxamide scaffold-based FT206 ^13^, was selected for clinical trials against myeloid leukemia, hemic and lymphatic diseases ^37,62^. For thienopyridine and thienopyrazine carboxamide-based USP25/28 inhibitors, we observed cell viability effects irrespective of the presence or absence of USP25/28, thereby suggesting off-target effects. This is reminiscent to other USP25/28 inhibitors that are known to target other proteins, such as *Otilonium bromide*, originally described as an approved drug for treating irritable bowel syndrome ^11^ and *Vismodegib*, previously described as a hedgehog signalling inhibitor ^35,63^. Together, these observations indicate multi-target recognition for such chemical core structures. Similarly, thienopyridine carboxamide scaffold containing medicinal drugs have been previously described, for example for inhibiting HIV Rev-response elements protein complexes ^64^. No noticeable off-target effects in translation/protein synthesis have been reported for these, suggesting that further derivatisation of thienopyridine / thienopyrazide carboxamide moieties to optimise targeting USP25/28 may have inadvertently increased affinity to off-target(s) in the translation apparatus.

Uncovering off-targets is challenging. Several approaches were used including chemoproteomics (**Figure 3**), genetic and proteomics screens and cellular adaptation experiments (see Supplementary Information for an extended discussion and **Figures S12-S17** for additional details). Combined, these extended analyses confirmed target-independent effects, all pointing towards an alternative mechanism of action for thienopyridine / thienopyrazide carboxamides within the translation machinery.

Other hallmarks were the observed decrease in global levels of conjugated ubiquitin upon FT224 treatment (**Figure S6f**) and effects on short-lived proteins (**Figure S14**), reminiscent of CHX mediated depletion of ubiquitin observed in yeast and due to preventing *de-novo* protein synthesis ^51^. The potential structural overlap between binding modes within USP25/28 and the tRNA E-site of the ribosomal 60S subunit is not obvious. For instance, we did not observe compound interference with tRNA charging (**Figure S17**). Generally, CHX and other compounds, such as *lactimidomycin*, *phyllanthoside*, *T-2 toxin*, *deoxynivalenol*, *verrucarin A*, *narciclasine*, *lycorine*, *nagilactone C*, *anisomycin*, *homoharringtonine* and *cryptopleurine* all bind to the same region in the 60S. This may reflect the presence of a small molecule “hot spot” ^65^ that might exert affinity for some thienopyridine/thienopyrazine carboxamides including FT224, FT811 and derivatives.

The USP28 – inhibitor structures provided an opportunity to generate refined inhibitors with reduced off-target effects, by identifying other chemical scaffolds away from the thienopyridine carboxamide moiety that correlated with a loss of translation inhibition (OPP activity) (**Figure 6, S10**). The chemical rationale for removing effects on translation may benefit from structural insights into the exact binding mode to the 60S ribosome “hot spot” region. Another approach taken here, was to determine minimal structural features for inhibitors to retain selectivity for USP25/28 inhibition. The thienopyrazine carboxamide scaffold seemed key, and we observed that inhibitors including molecular extensions of a bicyclic nature correlated with retained OPP activity (**Figure S11**). This observation may provide useful information regarding which parts of the inhibitor molecule need to be preserved to retain USP25/28 inhibition while dialling out off-target effects on translation. Refined USP25/28 inhibitors with minimal or no off-target on translation turned out to be more selective for squamous lung carcinoma in cell-based viability experiments (**Figure 7**). The potency is shifted towards the μM range, possibly reflecting their unoptimized state. AZ1 (FT214), another inhibitor based on a distinct chemical scaffold, not having any noticeable effect on translation, also displays a clear reduction in the growth of squamous lung carcinoma *in vivo* ^29^. *In vivo* potency of small molecule inhibitors may benefit from drug multi-specificity ^66^, which may be the case for the USP25/28 inhibitor CT1113 ^62^ as it shares the thienopyrazine (methylated amine) carboxamide scaffold with the compounds described in this study. Protein translation inhibitors are in various states of clinical evaluation ^67^.

In summary, the discovery of an off-target effect on protein translation related to the thienopyridine carboxamide moiety reveals insights into multidrug specificity. The development of refined USP25/28 inhibitor scaffolds provides the framework for more potent and potentially safer therapeutics targeting USP28 for the treatment of squamous lung cancer in particular, and perhaps other c-MYC-dependent cancer types in general.

## MATERIALS AND METHODS

A detailed description of the materials and methods used in this study can be found in the **Supplementary Information** section of the online article (URL Address).

## ACKNOWLEDGEMENTS

We are grateful for critical inputs from members of the Pinto-Fernandez, Kessler and Urbé & Clague laboratories and of the DUB Alliance (CRUK-Forma Therapeutics). We also thank the Discovery Proteomics Facility, led by Dr Iolanda Vendrell, and Dr Simon Davis and Marie-Laëtitia Thézénas for expert help with proteomics and mass spectrometry analysis. We thank Val Miller and Jennifer Ward for their expert help with microscopy and chemoproteomics experiments, respectively. We thank Dr Sebastian Nijman for the kind gift of the HAP1 USP28 knockout (KO) cell line. In addition, we thank Dr Ross Chapman (Oxford, UK) for generously providing the MCF7 USP28 knockout (KO) cell line. We also thank Dr Feng Zhang for providing the Human CRISPR Activation Library (SAM v1). This study was funded by Forma Therapeutics (Watertown, Boston, MA, USA). Work in the Kessler laboratory was supported by a John Fell Fund 133/075, the Wellcome Trust (097813/Z/11/Z), the Engineering and Physical Sciences Research Council (EP/N034295/1) and the Chinese Academy of Medical Sciences (CAMS) Innovation Fund for Medical Science (CIFMS), China (grant number: 2024-I2M-2-001-1).

## AUTHORSHIP CONTRIBUTIONS

B.M.K., S.U. and M.J.C. conceptualised the study. S.K., D.G., T.H., J.K., N.J., S.J.B., S.U., D.K., M.J.C. and B.M.K. directed projects involved in this study. A.P.F. and C.H. performed and directed biochemical and cell biological experiments. A.P.T. and W.W.K. performed structural studies. C.B., D.P., V.S. and L.M. assisted with the cell biology experiments. A.V., D.G. and S.J.B. designed and synthesised small molecule inhibitors. T.C., T. M.A., A.K. and A.P.F. performed chemical biology experiments. D.J., E.N.G. and D.E. performed genetic screening experiments. A.P.F., C.H., R.F., S.U. and B.M.K. performed data analysis. S.K., T.H., J.K., C.D., N.J. and D.K. have contributed to funding via the DUB Alliance supported by Forma Therapeutics. A.P.F., C.H, S.U., M.J.C. and B.M.K. wrote the manuscript. All co-authors have approved the finalised version.

## USE OF AI FOR MANUSCRIPT PREPARATION

We have used ChatGPT Edu University of Oxford (GPT-5.5) for minor text editing and for creating the illustration shown in **Figure 7h**.

## DATA AVAILABILITY

The crystal structure of human USP28 catalytic domain in complex with FT592 was deposited (D_1292155111) and is assigned to the accession code(s) PDB ID 29GK, Extended PDB ID pdb_000029GK.

Mass spectrometry (MS) data are available via ProteomeXchange with identifier PXD075950.

CRISPR sequencing data has been submitted to the GEO-NCBI-NIH repository under the ticket #25764846 (GEO Submission elenang@orcid).

## DECLARATION OF COMPETING INTERESTS

C.H., A.P.T., W.W.K., C.B., N.J. are employees of CRUK Cancer Research Horizon. T.R.H. is employed by TRH Consulting. S.K., D.G., J.K. and C.D. were employees of FORMA Therapeutics. This work was performed by the DUB Alliance and funded by FORMA Therapeutics.

## Abbreviations

c-MYC: oncogene transcription factor
CCND1: cyclin D1
USP28: ubiquitin specific protease 28
proteomics, CRISPR, MS/MS: tandem mass spectrometry, DUB inhibitor

## Notes

### Competing Interest Statement

C.H., A.P.T., W.W.K., C.B., N.J. are employees of CRUK Cancer Research Horizon. T.R.H. is employed by TRH Consulting. S.K., D.G., J.K. were employees of FORMA Therapeutics. This work was performed by the DUB Alliance and funded by FORMA Therapeutics.

### Summary of Updates

This version has been revised regarding minor edits on the text, modifications on Figure 7 and adding additional information to the figure legends.

## REFERENCES

1. Wu, X. et al. Small molecular inhibitors for KRAS-mutant cancers. Front. Immunol. 14, (2023).

2. Popow, J. et al. Targeting cancer with small-molecule pan-KRAS degraders. Science (1979). 385, 1338–1347 (2024).

3. Lewin, B. Oncogenic conversion by regulatory changes in transcription factors. Cell Preprint at 10.1016/0092-8674(91)90640-K (1991).

4. Harrigan, J. A., Jacq, X., Martin, N. M. & Jackson, S. P. Deubiquitylating enzymes and drug discovery: Emerging opportunities. Nature Reviews Drug Discovery Preprint at 10.1038/nrd.2017.152 (2018).

5. Altun, M. et al. Activity-based chemical proteomics accelerates inhibitor development for deubiquitylating enzymes. Chem. Biol. 18, (2011).

6. Reverdy, C. et al. Discovery of specific inhibitors of human USP7/HAUSP deubiquitinating enzyme. Chem. Biol. 19, 467–477 (2012).

7. Kategaya, L. et al. USP7 small-molecule inhibitors interfere with ubiquitin binding. Nature https://doi.org/10.1038/nature24006 (2017) doi:10.1038/nature24006.

8. Turnbull, A. P. et al. Molecular basis of USP7 inhibition by selective small-molecule inhibitors. Nature 550, (2017).

9. Lamberto, I. et al. Structure-Guided Development of a Potent and Selective Non-covalent Active-Site Inhibitor of USP7. Cell Chem. Biol. 24, 1490–1500.e11 (2017).

10. Liu, Z. et al. Discovery of [1,2,3]triazolo[4,5-d]pyrimidine derivatives as highly potent, selective, and cellularly active USP28 inhibitors. Acta Pharm. Sin. B https://doi.org/10.1016/j.apsb.2019.12.008 (2019) doi:10.1016/j.apsb.2019.12.008.

11. Xu, Z. et al. Otilonium Bromide acts as a selective USP28 inhibitor and exhibits cytotoxic activity against multiple human cancer cell lines. Biochem. Pharmacol. 215, 115746 (2023).

12. Wrigley, J. D. et al. Identification and Characterization of Dual Inhibitors of the USP25/28 Deubiquitinating Enzyme Subfamily. ACS Chem. Biol. 12, 3113–3125 (2017).

13. Josue Ruiz, E., et al. Usp28 deletion and small-molecule inhibition destabilizes c-myc and elicits regression of squamous cell lung carcinoma. Elife 10, (2021).

14. Davis, R. J., Welcker, M. & Clurman, B. E. Tumor suppression by the Fbw7Ubiquitin ligase: Mechanisms and opportunities. Cancer Cell Preprint at 10.1016/j.ccell.2014.09.013 (2014).

15. Popov, N. et al. The ubiquitin-specific protease USP28 is required for MYC stability. Nat. Cell Biol. https://doi.org/10.1038/ncb1601 (2007) doi:10.1038/ncb1601.

16. Popov, N., Herold, S., Llamazares, M., Schülein, C. & Eilers, M. Fbw7 and Usp28 regulate Myc protein stability in response to DNA damage. Cell Cycle https://doi.org/10.4161/cc.6.19.4804 (2007) doi:10.4161/cc.6.19.4804.

17. Wang, X. et al. Targeting deubiquitinase USP28 for cancer therapy. Cell Death and Disease Preprint at 10.1038/s41419-017-0208-z (2018).

18. Sun, X. X. et al. The nucleolar ubiquitin-specific protease USP36 deubiquitinates and stabilizes c-Myc. Proc. Natl. Acad. Sci. U. S. A. https://doi.org/10.1073/pnas.1411713112 (2015) doi:10.1073/pnas.1411713112.

19. Cuella-Martin, R. et al. 53BP1 Integrates DNA Repair and p53-Dependent Cell Fate Decisions via Distinct Mechanisms. Mol. Cell https://doi.org/10.1016/j.molcel.2016.08.002 (2016) doi:10.1016/j.molcel.2016.08.002.

20. Fong, C. S. et al. 53BP1 and USP28 mediate p53- dependent cell cycle arrest in response to centrosome loss and prolonged mitosis. Elife https://doi.org/10.7554/eLife.16270 (2016) doi:10.7554/eLife.16270.

21. Belal, H., Ng, E. F. Y., Ohta, M. & Meitinger, F. Cancer-associated USP28 missense mutations disrupt 53BP1 interaction and p53 stabilization. Nat. Commun. 16, 10310 (2025).

22. Knobel, P. A. et al. USP28 Is Recruited to Sites of DNA Damage by the Tandem BRCT Domains of 53BP1 but Plays a Minor Role in Double-Strand Break Metabolism. Mol. Cell. Biol. https://doi.org/10.1128/mcb.00197-14 (2014) doi:10.1128/mcb.00197-14.

23. Lambrus, B. G. et al. A USP28-53BP1-p53-p21 signaling axis arrests growth after centrosome loss or prolonged mitosis. Journal of Cell Biology https://doi.org/10.1083/jcb.201604054 (2016) doi:10.1083/jcb.201604054.

24. Meitinger, F. et al. 53BP1 and USP28 mediate p53 activation and G1 arrest after centrosome loss or extended mitotic duration. Journal of Cell Biology https://doi.org/10.1083/jcb.201604081 (2016) doi:10.1083/jcb.201604081.

25. Weili, Z., Zhikun, L., Jianmin, W. & Qingbao, T. Knockdown of USP28 enhances the radiosensitivity of esophageal cancer cells via the c-Myc/hypoxia-inducible factor-1 alpha pathway. J Cell Biochem 120, 201–212 (2019).

26. Ito, F., Yoshimoto, C., Yamada, Y., Sudo, T. & Kobayashi, H. The HNF-1β-USP28-Claspin pathway upregulates DNA damage-induced Chk1 activation in ovarian clear cell carcinoma. Oncotarget https://doi.org/10.18632/oncotarget.24776 (2018) doi:10.18632/oncotarget.24776.

27. Zhao, L. J. et al. USP28 contributes to the proliferation and metastasis of gastric cancer. J. Cell. Biochem. https://doi.org/10.1002/jcb.28040 (2019) doi:10.1002/jcb.28040.

28. Li, P., Huang, Z., Wang, J., Chen, W. & Huang, J. Ubiquitin-specific peptidase 28 enhances STAT3 signaling and promotes cell growth in non-small-cell lung cancer. Onco. Targets. Ther. https://doi.org/10.2147/OTT.S194917 (2019) doi:10.2147/OTT.S194917.

29. Prieto-Garcia, C., et al. Maintaining protein stability of ΔNp63 via USP28 is required by squamous cancer cells. EMBO Mol. Med. e11101, (2020).

30. Ren, Y. et al. Targeting USP28 inhibits clear cell renal cell carcinoma growth. Cell. Signal. 139, 112344 (2026).

31. Saei, A. et al. Loss of USP28-mediated BRAF degradation drives resistance to RAF cancer therapies. Journal of Experimental Medicine https://doi.org/10.1084/jem.20171960 (2018) doi:10.1084/jem.20171960.

32. Richter, K. et al. USP28 deficiency promotes breast and liver carcinogenesis as well as tumor angiogenesis in a HIF-independent manner. Molecular Cancer Research https://doi.org/10.1158/1541-7786.MCR-17-0452 (2018) doi:10.1158/1541-7786.MCR-17-0452.

33. Müller, I. et al. Cancer Cells Employ Nuclear Caspase-8 to Overcome the p53-Dependent G2/M Checkpoint through Cleavage of USP28. Mol. Cell https://doi.org/10.1016/j.molcel.2019.12.023 (2020) doi:10.1016/j.molcel.2019.12.023.

34. Prieto-Garcia, C., Tomašković, I., Shah, V. J., Dikic, I. & Diefenbacher, M. USP28: Oncogene or Tumor Suppressor? A Unifying Paradigm for Squamous Cell Carcinoma. Cells 10, (2021).

35. Wang, H. et al. USP28 and USP25 are downregulated by Vismodegib in vitro and in colorectal cancer cell lines. FEBS J. 288, 1325–1342 (2021).

36. Varca, A. C. et al. Identification and validation of selective deubiquitinase inhibitors. Cell Chem. Biol. 28, 1758–1771.e13 (2021).

37. Xu, J. et al. Preclinical testing of CT1113, a novel USP28 inhibitor, for the treatment of T-cell acute lymphoblastic leukaemia. Br. J. Haematol. 204, 2301–2318 (2024).

38. Cai, C. et al. The deubiquitinase USP28 promotes esophageal squamous cell carcinoma proliferation by stabilizing ΔNp63 protein. Cell. Signal. 143, 112489 (2026).

39. Gersch, M. et al. Distinct USP25 and USP28 Oligomerization States Regulate Deubiquitinating Activity. Mol. Cell https://doi.org/10.1016/j.molcel.2019.02.030 (2019) doi:10.1016/j.molcel.2019.02.030.

40. Patzke, J. V. et al. Structural basis for the bi-specificity of USP25 and USP28 inhibitors. EMBO Rep. 25, 2950–2973 (2024).

41. Zhou, D. et al. Structure-based discovery of potent USP28 inhibitors derived from Vismodegib. Eur. J. Med. Chem. 254, 115369 (2023).

42. Wang, F., et al. Molecular Basis of USP28 Allosteric Inhibition by Small-Molecule Inhibitor AZ1. https://papers.ssrn.com/sol3/papers.cfm?abstract_id=4676302 (2024).

43. Nelson, J. K. et al. USP25 promotes pathological HIF-1-driven metabolic reprogramming and is a potential therapeutic target in pancreatic cancer. Nat. Commun. 13, 2070 (2022).

44. Hernandez-Olmos, V. et al. Structure Merging Approach Leads to New Dual Potent and Selective USP25/USP28 Inhibitors. J. Med. Chem. 69, 10140–10168 (2026).

45. Zablocki, M.-M. et al. CARBOXAMIDES AS UBIQUITIN-SPECIFIC PROTEASE INHIBITORS. Preprint at https://patentscope.wipo.int/search/en/detail.jsf?docId=WO2019032863 (2019).

46. Guerin, D. et al. THIENOPYRIDINE CARBOXAMIDES AS UBIQUITIN-SPECIFIC PROTEASE INHIBITORS. 1–361 Preprint at https://patentscope.wipo.int/search/en/detail.jsf?docId=WO2017139778 (2017).

47. Bratt, A. S. et al. Pharmacologic interrogation of USP28 cellular function in p53 signaling. Cell Chem. Biol. 32, 1166–1182.e27 (2025).

48. Sancho, R. et al. F-box and WD Repeat Domain-Containing 7 Regulates Intestinal Cell Lineage Commitment and Is a Haploinsufficient Tumor Suppressor. Gastroenterology 139, 929–941 (2010).

49. Schülein-Völk, C. et al. Dual Regulation of Fbw7 Function and Oncogenic Transformation by Usp28. Cell Rep. 9, 1099–1109 (2014).

50. Garreau de Loubresse, N., et al. Structural basis for the inhibition of the eukaryotic ribosome. Nature 513, 517–522 (2014).

51. John, H., S., L. D. & Daniel, F. Ubiquitin Depletion as a Key Mediator of Toxicity by Translational Inhibitors. Mol. Cell. Biol. 23, 9251–9261 (2003).

52. Liu, B., Sureda-Gómez, M., Zhen, Y., Amador, V. & Reverter, D. A quaternary tetramer assembly inhibits the deubiquitinating activity of USP25. Nat. Commun. https://doi.org/10.1038/s41467-018-07510-5 (2018) doi:10.1038/s41467-018-07510-5.

53. Sauer, F. et al. Differential Oligomerization of the Deubiquitinases USP25 and USP28 Regulates Their Activities. Mol. Cell https://doi.org/10.1016/j.molcel.2019.02.029 (2019) doi:10.1016/j.molcel.2019.02.029.

54. Varca, A. C., Casalena, D., Auld, D. & Buhrlage, S. J. Identification of deubiquitinase inhibitors via high-throughput screening using a fluorogenic ubiquitin-rhodamine assay. STAR Protoc. 2, 100896 (2021).

55. Chan, W. C. et al. Accelerating inhibitor discovery for deubiquitinating enzymes. Nat. Commun. 14, 686 (2023).

56. Prieto-Garcia, C. et al. USP28 enables oncogenic transformation of respiratory cells, and its inhibition potentiates molecular therapy targeting mutant EGFR, BRAF and PI3K. Mol. Oncol. 16, 3082–3106 (2022).

57. Shih, I. M. & Wang, T. L. Notch signaling, γ-secretase inhibitors, and cancer therapy. Cancer Research Preprint at 10.1158/0008-5472.CAN-06-3958 (2007).

58. Gazdar, A. F. & Minna, J. D. Small cell lung cancers made from scratch. Journal of Experimental Medicine https://doi.org/10.1084/jem.20182216 (2019) doi:10.1084/jem.20182216.

59. Wu, Y. et al. The Deubiquitinase USP28 Stabilizes LSD1 and Confers Stem-Cell-like Traits to Breast Cancer Cells. Cell Rep. https://doi.org/10.1016/j.celrep.2013.08.030 (2013) doi:10.1016/j.celrep.2013.08.030.

60. Peng, J. et al. Identification of a class of potent USP25/28 inhibitors with broad-spectrum anti-cancer activity. Signal Transduct. Target. Ther. 7, 393 (2022).

61. Li, J. et al. The deubiquitinase USP28 maintains the expression of the transcription factor MYCN and is essential in neuroblastoma cells. Journal of Biological Chemistry 299, 104856 (2023).

62. Zhou, S., et al. Abstract LB367: CT1113, a dual USP25/28 inhibitor, promotes antitumor immunity by preventing YTHDF2-mediated complement activation and potentiates anti-PD-1 therapy. Cancer Res. 85, LB367–LB367 (2025).

63. Aditya, S. & Rattan, A. Vismodegib: A smoothened inhibitor for the treatment of advanced basal cell carcinoma. Indian Dermatol. Online J. 4, 365–8 (2013).

64. Nakamura, R. et al. Identification and Optimization of Thienopyridine Carboxamides as Inhibitors of HIV Regulatory Complexes. Antimicrob. Agents Chemother. 61, e02366–16 (2021).

65. Lin, A. et al. Off-target toxicity is a common mechanism of action of cancer drugs undergoing clinical trials. Sci. Transl. Med. https://doi.org/10.1126/scitranslmed.aaw8412 (2019) doi:10.1126/scitranslmed.aaw8412.

66. Deshaies, R. J. Multispecific drugs herald a new era of biopharmaceutical innovation. Nature 580, 329–338 (2020).

67. Jia, X. et al. Protein translation: biological processes and therapeutic strategies for human diseases. Signal Transduct. Target. Ther. 9, 44 (2024).

