## SUPPLEMENTAL FIGURES 1-32 for "Refined USP25/28 inhibitors with enhanced selectivity towards c-Myc driven squamous lung cancer cells"

### Figure S1

**a**

FT811 | FT214 / AZ1

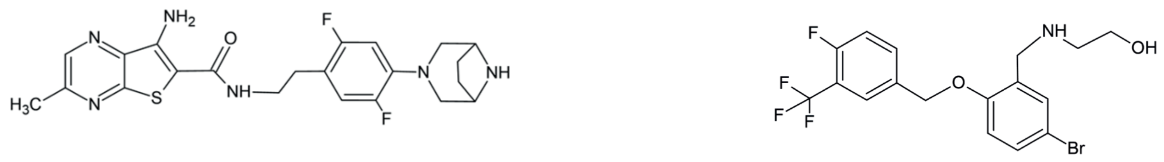

Thienopyrazine carboxamide scaffold

Benzyl ethanolamine scaffold

**b**

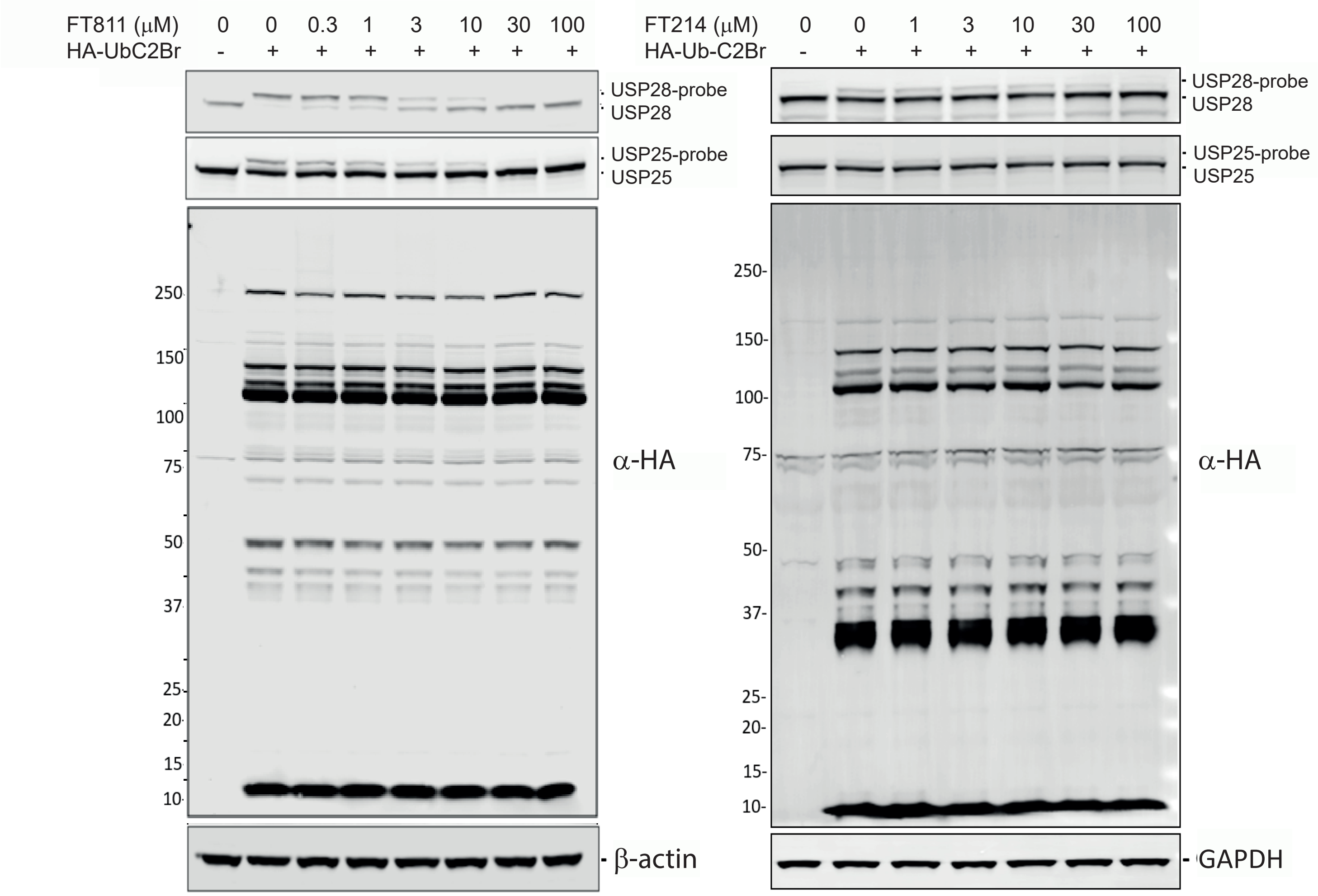

**Figure S2**

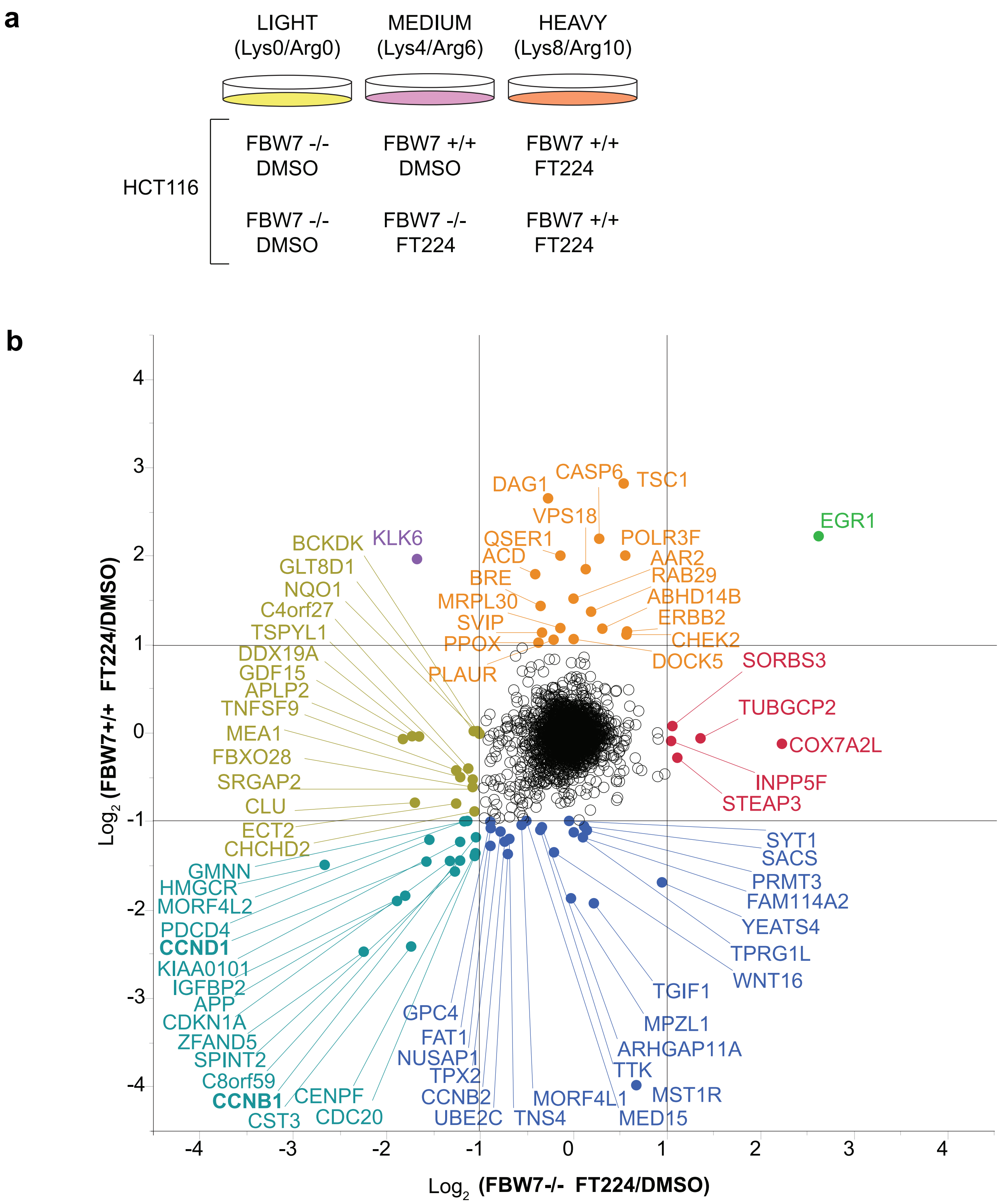
a
LIGHT (Lys0/Arg0)
MEDIUM (Lys4/Arg6)
HEAVY (Lys8/Arg10)
HCT116
FBW7 -/- DMSO
FBW7 +/+ DMSO
FBW7 +/+ FT224
FBW7 -/- DMSO
FBW7 -/- FT224
FBW7 +/+ FT224
b
Log2 (FBW7+/+ FT224/DMSO)
Log2 (FBW7-/- FT224/DMSO)
4
3
2
1
0
-1
-2
-3
-4
EGR1
KLK6
DAG1
CASP6
TSC1
VPS18
QSER1
POLR3F
ACD
AAR2
BRE
RAB29
MRPL30
ABHD14B
SVIP
ERBB2
PPOX
CHEK2
PLAUR
DOCK5
SORBS3
TUBGCP2
COX7A2L
INPP5F
STEAP3
BCKDK
GLT8D1
NQO1
C4orf27
TSPYL1
DDX19A
GDF15
APLP2
TNFSF9
MEA1
FBXO28
SRGAP2
CLU
ECT2
CHCHD2
GMNN
HMGCR
MORF4L2
PDCD4
CCND1
KIAA0101
IGFBP2
APP
CDKN1A
ZFAND5
SPINT2
C8orf59
CCNB1
CST3
CENPF
CDC20
GPC4
FAT1
NUSAP1
TPX2
CCNB2
UBE2C
TNS4
MORF4L1
MED15
MST1R
TTK
ARHGAP11A
MPZL1
TGIF1
WNT16
TPRG1L
YEATS4
FAM114A2
PRMT3
SACS
SYT1

#### Figure S3

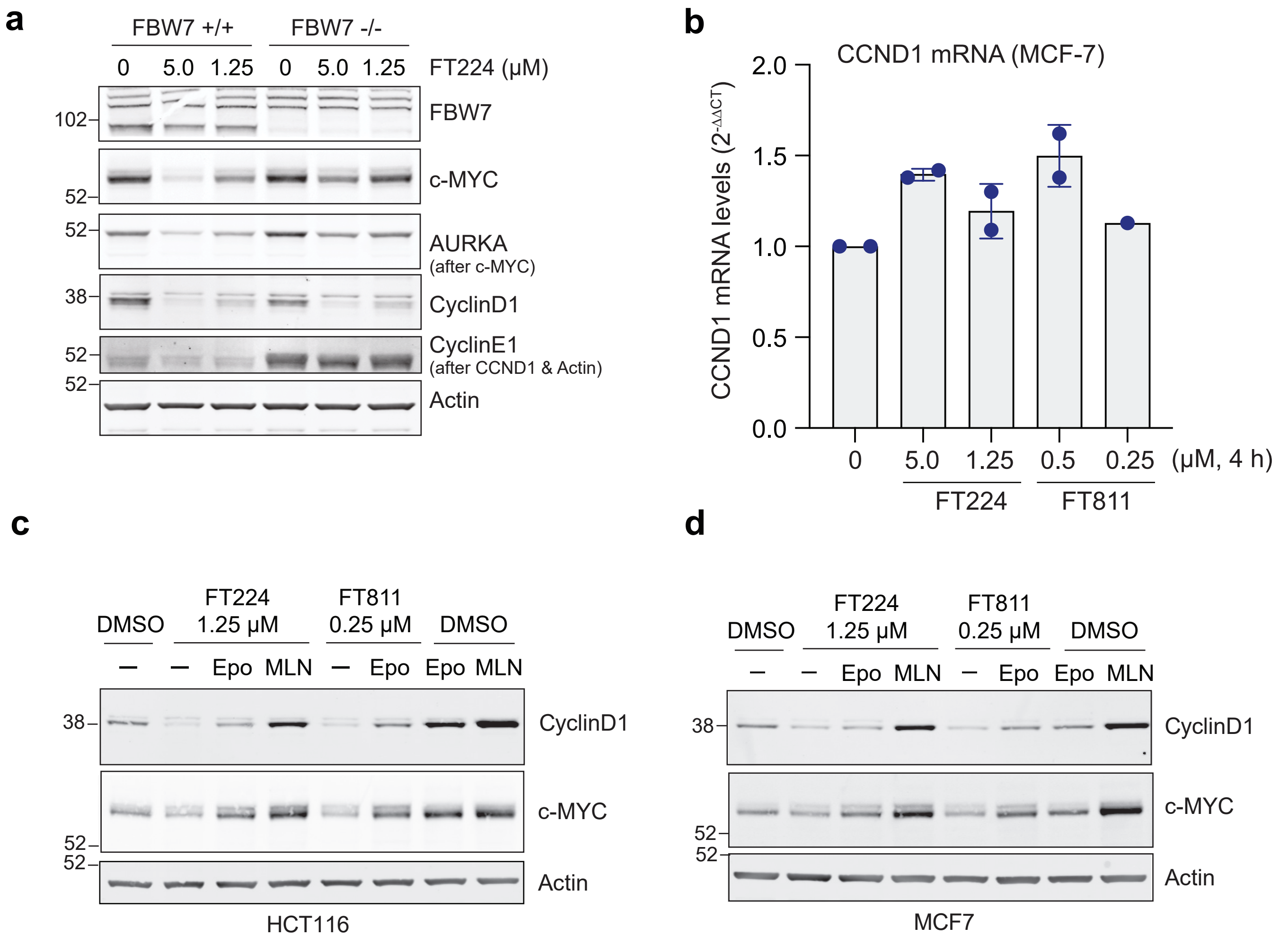
a
FBW7 +/+
FBW7 -/-
0 5.0 1.25 0 5.0 1.25 FT224 (µM)
102
FBW7
c-MYC
52
52
AURKA
(after c-MYC)
38
CyclinD1
52
CyclinE1
(after CCND1 & Actin)
52
Actin
b
CCND1 mRNA (MCF-7)
CCND1 mRNA levels (2-ΔΔCT)
2.0
1.5
1.0
0.5
0.0
0 5.0 1.25 0.5 0.25 (µM, 4 h)
FT224
FT811
c
DMSO
FT224 1.25 µM
FT811 0.25 µM
DMSO
– – Epo MLN – Epo Epo MLN
38
CyclinD1
c-MYC
52
52
Actin
HCT116
d
DMSO
FT224 1.25 µM
FT811 0.25 µM
DMSO
– – Epo MLN – Epo Epo MLN
38
CyclinD1
c-MYC
52
52
Actin
MCF7

### Figure S4

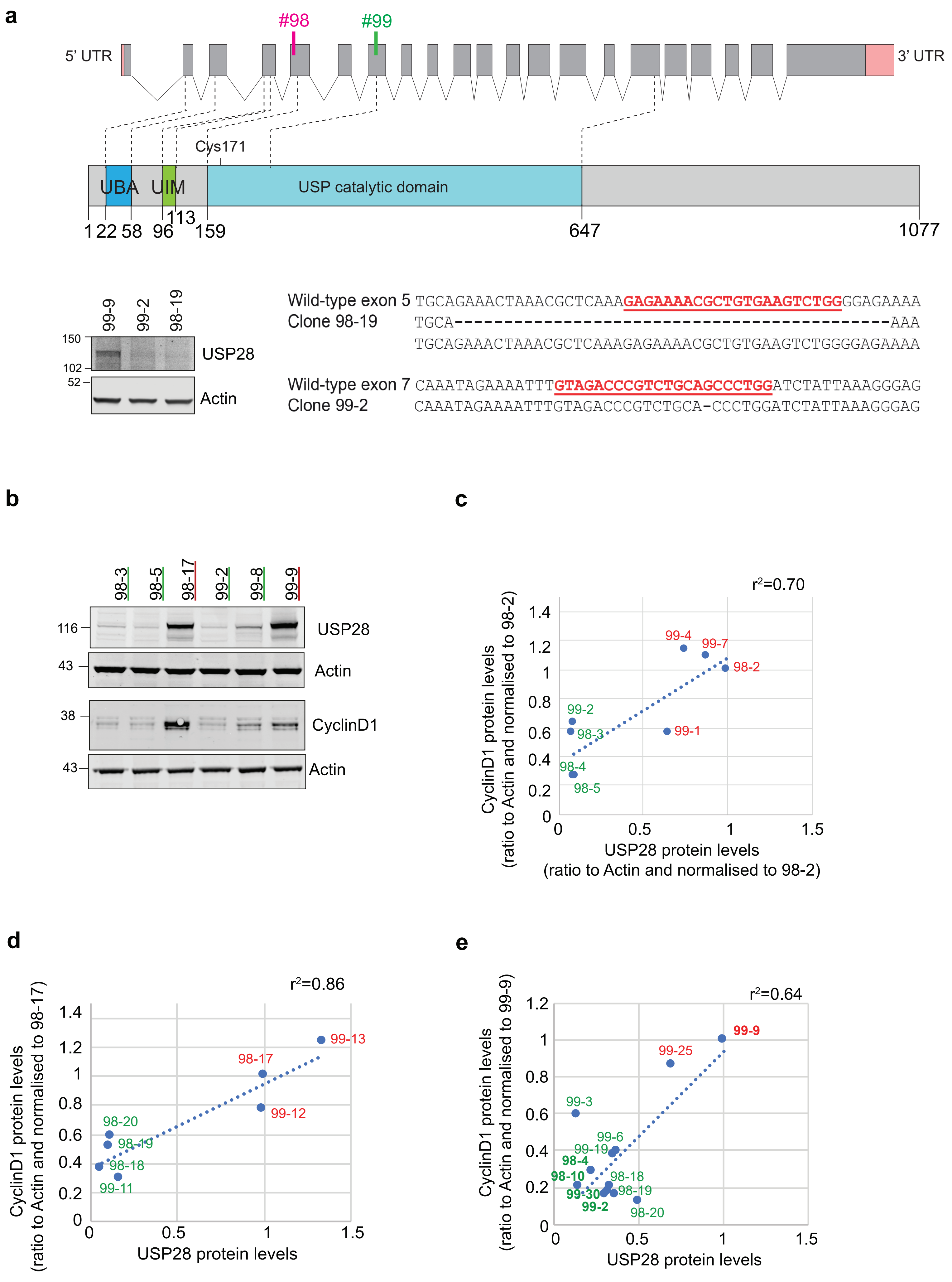
a
#98
#99
5' UTR
3' UTR
Cys171
UBA
UIM
USP catalytic domain
1 22 58 96 113 159 647 1077
99-9 99-2 98-19
150
102
52
USP28
Actin
Wild-type exon 5 TGCAGAAACTAAACGCTCAAAGAGAAAACGCTGTGAAGTCTGGGGAGAAAA
Clone 98-19 TGCA----------------------------------------------AAA
TGCAGAAACTAAACGCTCAAAGAGAAAACGCTGTGAAGTCTGGGGAGAAAA
Wild-type exon 7 CAAATAGAAAATTTGTAGACCCGTCTGCAGCCCTGGATCTATTAAAGGGAG
Clone 99-2 CAAATAGAAAATTTGTAGACCCGTCTGCA-CCCTGGATCTATTAAAGGGAG
b
98-3 98-5 98-17 99-2 99-8 99-9
116
43
38
43
USP28
Actin
CyclinD1
Actin
c
r²=0.70
99-4
99-7
98-2
99-2
98-3
99-1
98-4
98-5
CyclinD1 protein levels (ratio to Actin and normalised to 98-2)
USP28 protein levels (ratio to Actin and normalised to 98-2)
d
r²=0.86
99-13
98-17
99-12
98-20
98-19
98-18
99-11
CyclinD1 protein levels (ratio to Actin and normalised to 98-17)
USP28 protein levels
e
r²=0.64
99-9
99-25
99-3
99-6
99-19
98-4
98-10
98-18
99-30
98-19
99-2
98-20
CyclinD1 protein levels (ratio to Actin and normalised to 99-9)
USP28 protein levels

#### Figure S5

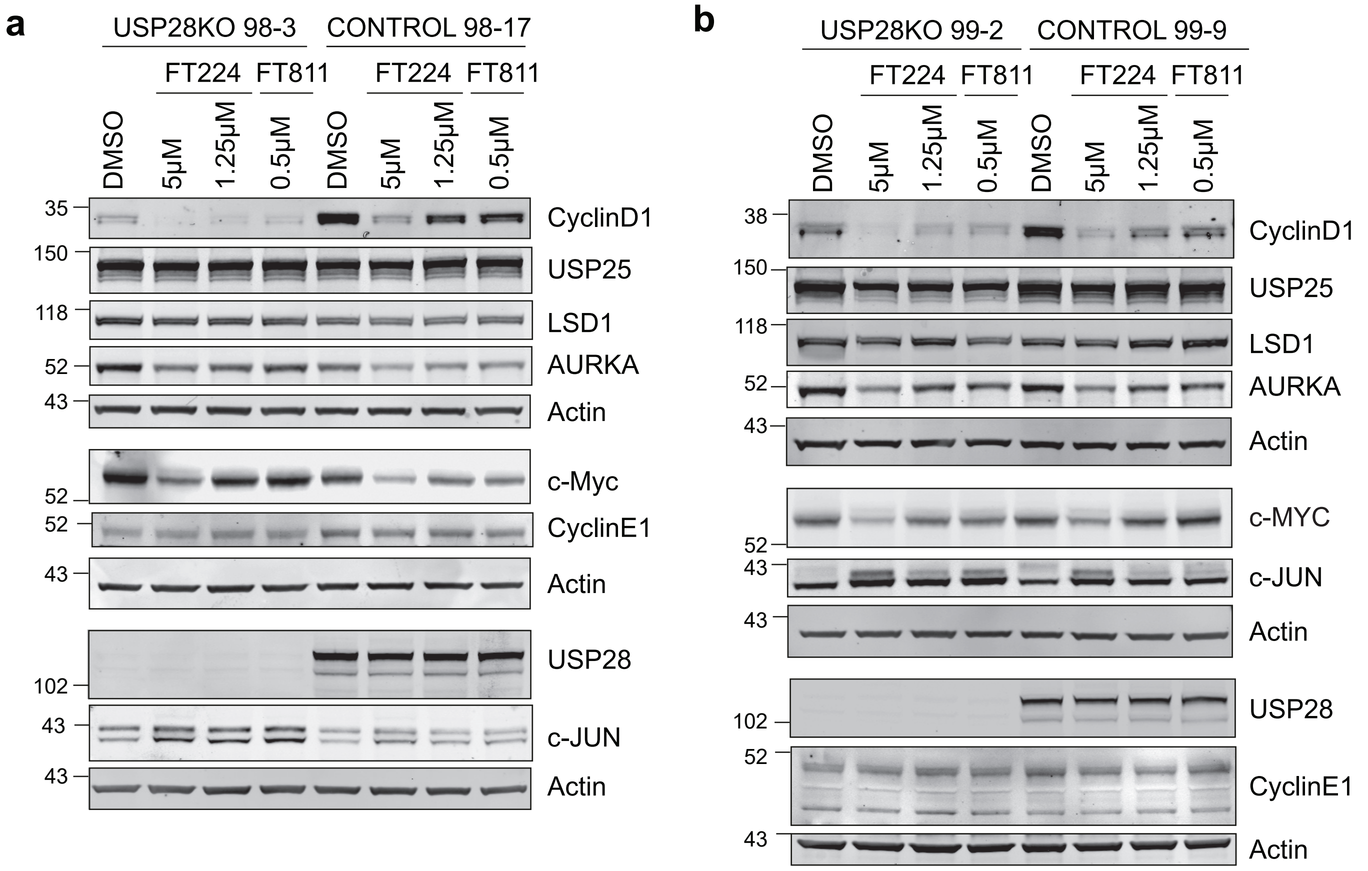

a
USP28KO 98-3
CONTROL 98-17
FT224
FT811
DMSO
5μM
1.25μM
0.5μM
CyclinD1
USP25
LSD1
AURKA
Actin
c-Myc
CyclinE1
USP28
c-JUN
35
150
118
52
43
102
b
USP28KO 99-2
CONTROL 99-9
c-MYC
38

#### Figure S6

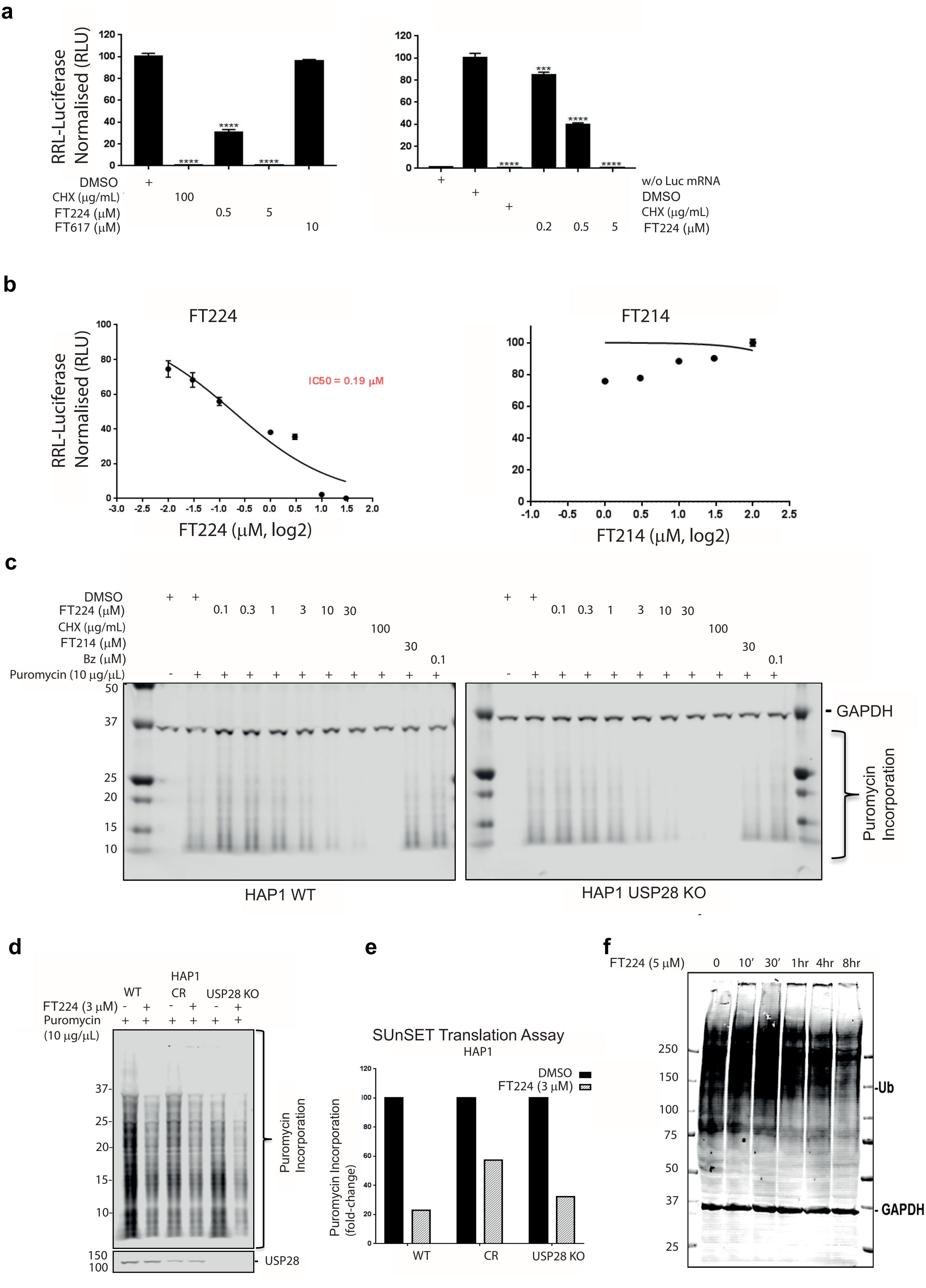

a
RRL-Luciferase Normalised (RLU)
DMSO
CHX (μg/mL)
FT224 (μM)
FT617 (μM)
100
0.5
5
10
w/o Luc mRNA
DMSO
CHX (μg/mL)
FT224 (μM)
0.2
0.5
5
b
FT224
FT214
IC50 = 0.19 μM
FT224 (μM, log2)
FT214 (μM, log2)
c
DMSO
FT224 (μM)
CHX (μg/mL)
FT214 (μM)
Bz (μM)
Puromycin (10 μg/μL)
0.1 0.3 1 3 10 30
100
30
0.1
GAPDH
Puromycin Incorporation
HAP1 WT
HAP1 USP28 KO
d
HAP1
WT
CR
USP28 KO
FT224 (3 μM)
Puromycin (10 μg/μL)
Puromycin Incorporation
USP28
e
SUnSET Translation Assay
HAP1
DMSO
FT224 (3 μM)
Puromycin Incorporation (fold-change)
WT
CR
USP28 KO
f
FT224 (5 μM)
0 10′ 30′ 1hr 4hr 8hr
Ub
GAPDH

Figure S7

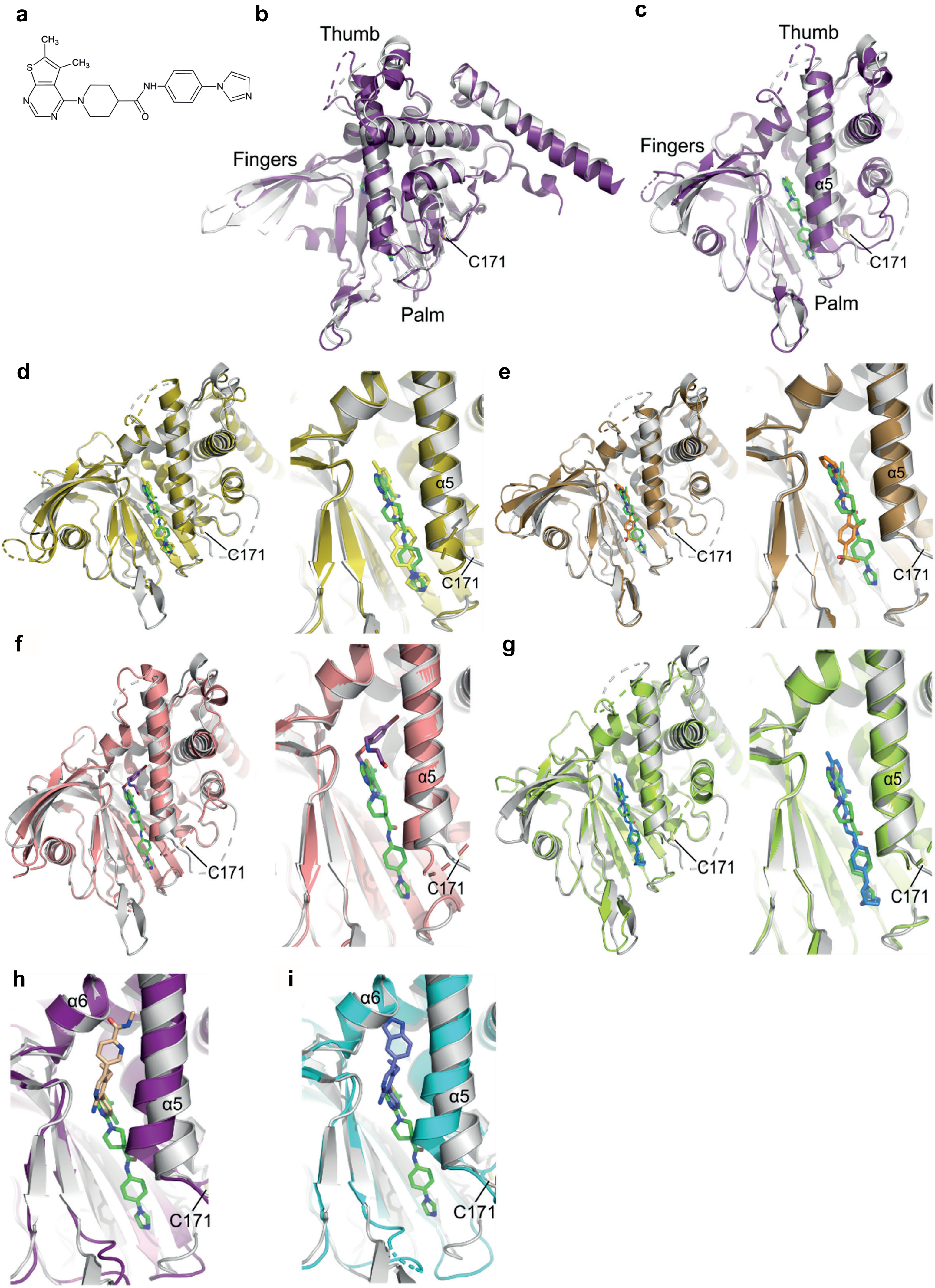
a
CH3
CH3
S
N
N
N
NH
O
N
N
b
Thumb
Fingers
C171
Palm
c
Thumb
Fingers
α5
C171
Palm
d
C171
α5
C171
e
C171
α5
C171
f
C171
α5
C171
g
C171
α5
C171
h
α6
α5
C171
i
α6
α5
C171

### Figure S8

**a** Translation inhibition (assayed by OPP) versus cell viability (USP25/28 inhibitors)

| Compound name | IC50 values (OPP-Protein Synthesis) | IC50 values (WT confluence – Incucyte) | IC50 values (USP28 KO confluence – Incucyte) |
|---|---|---|---|
| CRT519 expt.1 | 0.0250 (µM) | 0.006059 (µM) | 0.002065 (µM) |
| CRT519 expt.2 | 0.0535 (µM) | 0.02129 (µM) | 0.07835 (µM) |
| AZ1 expt. 1 | ~ 9.5 (µM) | 70.99 (µM) | 11.28 (µM) |
| bin-01-68 expt.1 | ~ 29.9 (µM) | 7.515e+099 (µM) | - (µM) |
| bin-01-68 expt.2 | 33.1 (µM) | - (µM) | - (µM) |
| bin-01-07-07 expt.1 | 16.9 (µM) | 338.0 (µM) | 5.188 (µM) |
| bin-01-74 expt. 1 | 1.0 (µM) | 0.8959 (µM) | 0.6079 (µM) |
| AV-11324-34 expt. 1 | ~ 34.0 (µM) | 20.16 (µM) | - (µM) |
| bin-01-76A | 65.5 (µM) | 13.81 (µM) | 194.2 (µM) |
| bin-01-106 | 27.9 (µM) | 73.58 (µM) | 101.1 (µM) |

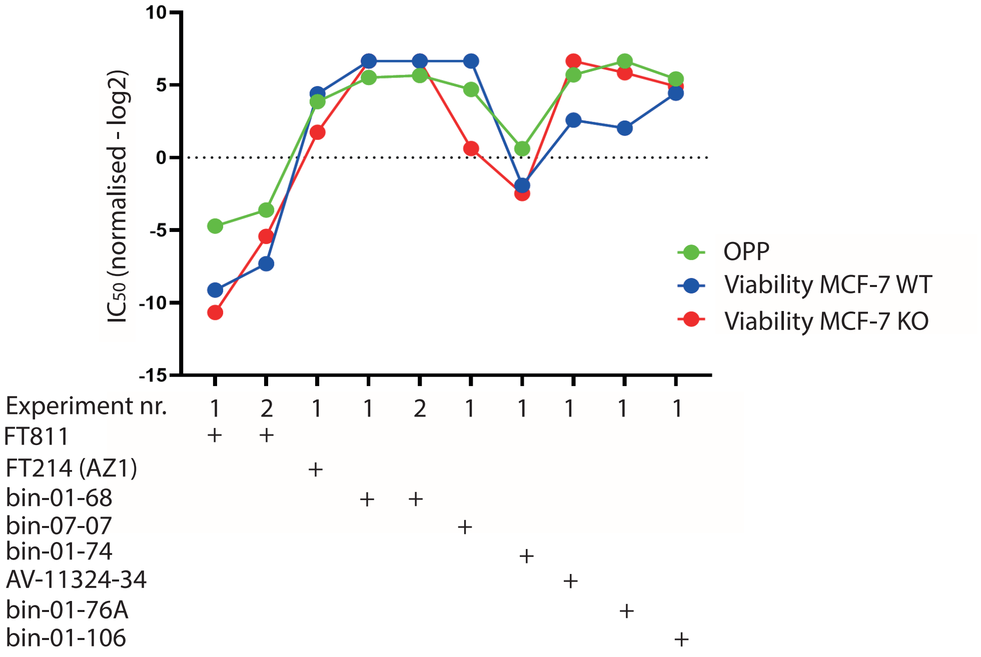


**b**

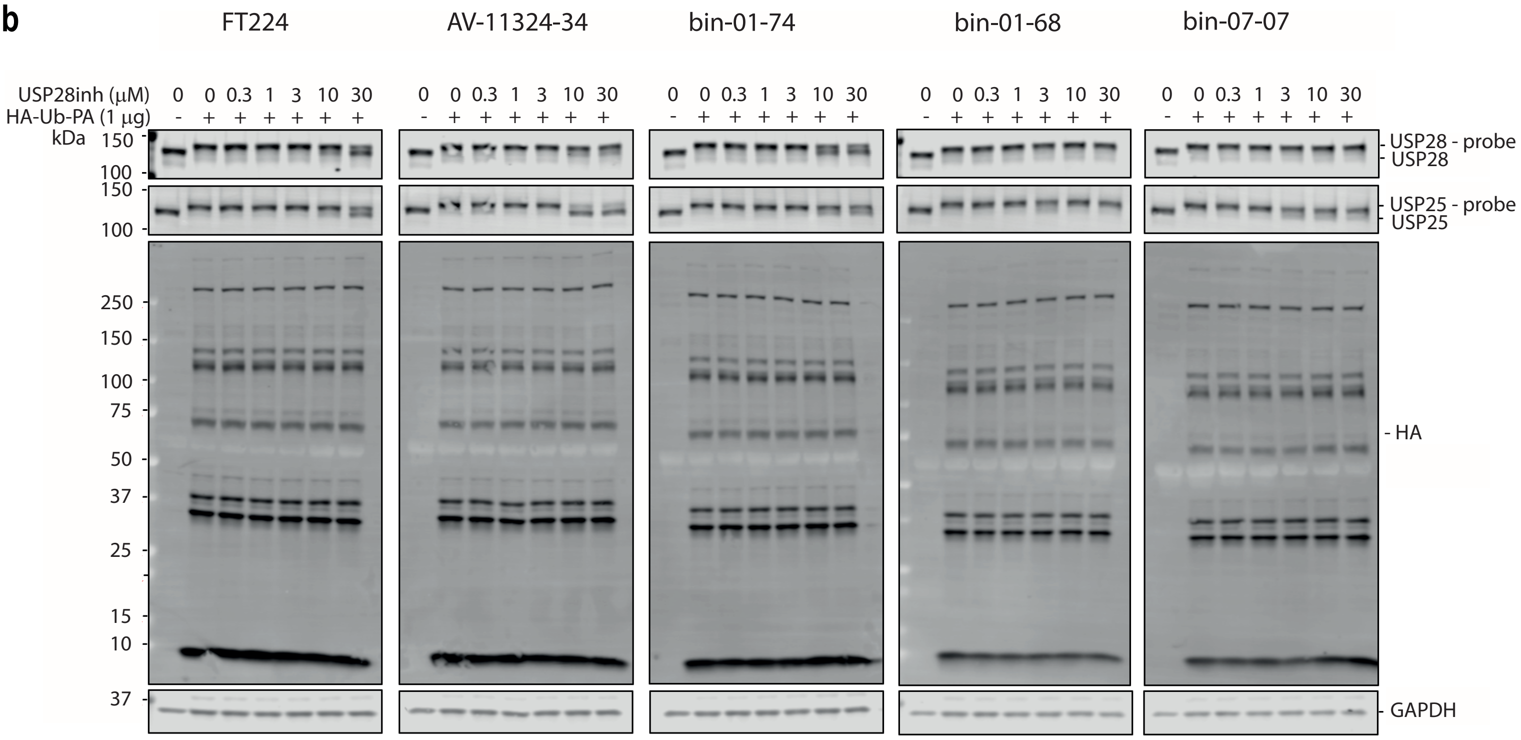

### Figure S9

| Compound ID | Structure | Ub-Rho IC50 (μM) |
|---|---|---|
| bin-01-41 | 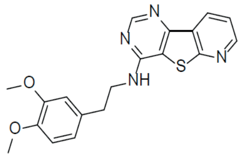 | 5.931 |
| bin-01-50 | 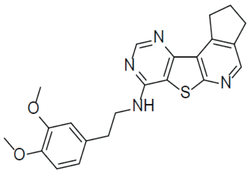 | 2.933 |
| bin-01-55 | 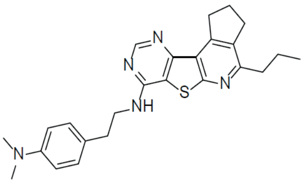 | 0.1995 |
| bin-01-60 | 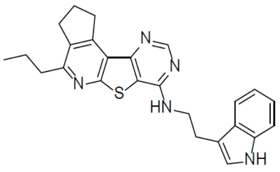 | 1.332 |
| bin-01-70 | 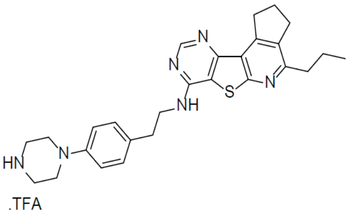 | 0.01669 |

| | | |
|---|---|---|
| bin-01-111 | 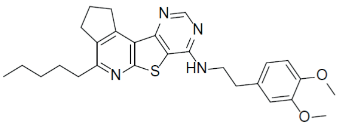 | 0.1003 |
| bin-01-112 | 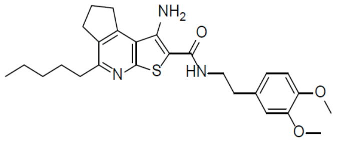 | 0.102 |
| bin-01-68 | 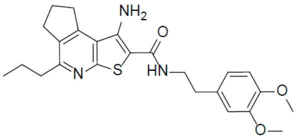 | 0.02887 |
| binh-01-07-07 | 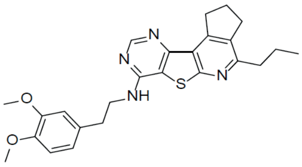 | 0.05803 |
| bin-01-74 | 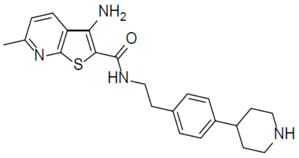 | 0.04031 |
| AV-11324-34 | 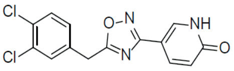 | 0.332 ± 0.07 |

| | | |
|---|---|---|
| bin-01-71 | 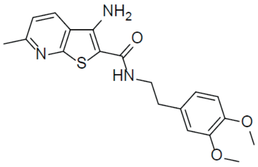 | 0.1386 |
| bin-01-73 | 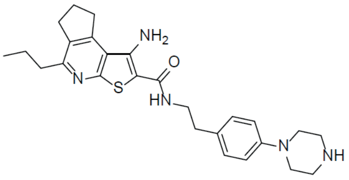 | 0.0312 |
| bin-01-76A | 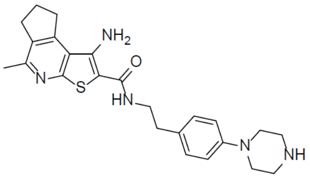 | 0.02209 |
| bin-01-78 | 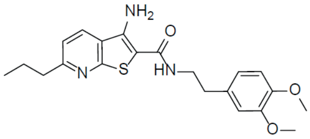 | 0.00674 |
| bin-01-106 | 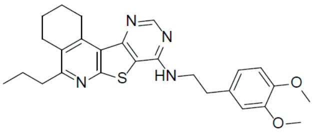 | 0.006892 |

| | | |
|---|---|---|
| AZ1 / FT214 | 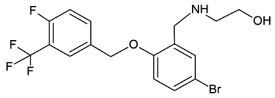 | 0.472 |

| | | |
|---|---|---|
| bin-01-98 | 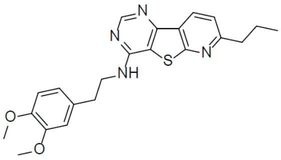 | 0.006876 |

| | | |
|---|---|---|
| FT206 | 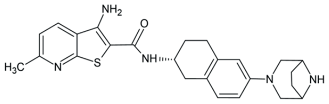 | 0.205 |
| FT224 | 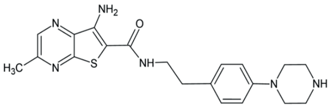 | 0.7 |
| FT529 | 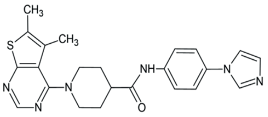 | |
| FT639 | 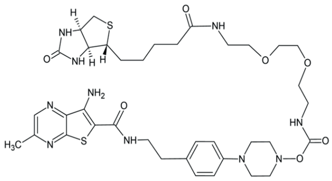 | |
| FT811 | 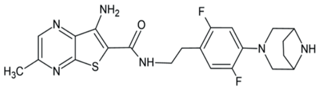 | 1.27 |

### Figure S10

| Compound name | IC50 values (OPP-Protein Synthesis) |
|---|---|
| Cycloheximide (CHX) expt. 1 | 3.592e-008 (µg/mL) |
| Cycloheximide (CHX) expt. 1 | 0.001080 (µg/mL) |
| CRT519 expt.1 | 0.0250 (µM) |
| CRT519 expt.2 | 0.0535 (µM) |
| CRT519 expt. 3 | 0.1991 (µM) |
| AZ1 expt. 1 | ~ 9.5 (µM) |
| AZ1 expt. 2 | 7.8 (µM) |
| bin-01-68 expt.1 | ~ 29.9 (µM) |
| bin-01-68 expt.2 | 33.1 (µM) |
| bin-01-07-07 expt.1 | 16.9 (µM) |
| bin-01-07-07 expt.2 | 12.9 (µM) |
| bin-01-74 expt. 1 | 1.0 (µM) |
| bin-01-74 expt. 2 | 0.7 (µM) |
| AV-11324-34 expt. 1 | ~ 34.0 (µM) |
| AV-11324-34 expt. 2 | Inactive |
| bin-01-98 expt. 1 | 5.7 (µM) |
| bin-01-98 expt. 2 | 5.9 (µM) |
| bin-01-41 | 8.5 (µM) |
| bin-01-50 | 15.5 (µM) |
| bin-01-55 | 33.1 (µM) |
| bin-01-60 | 17.9 (µM) |
| bin-01-70 | 5.8 (µM) |
| bin-01-71 | 111.3 (µM) |
| bin-01-73 | 4.9 (µM) |
| bin-01-76A | 65.5 (µM) |
| bin-01-78 | 23.0 (µM) |
| bin-01-106 | 27.9 (µM) |
| bin-01-111 | 19.0 (µM) |
| bin-01-112 | 67.0 (µM) |

**Figure S11**

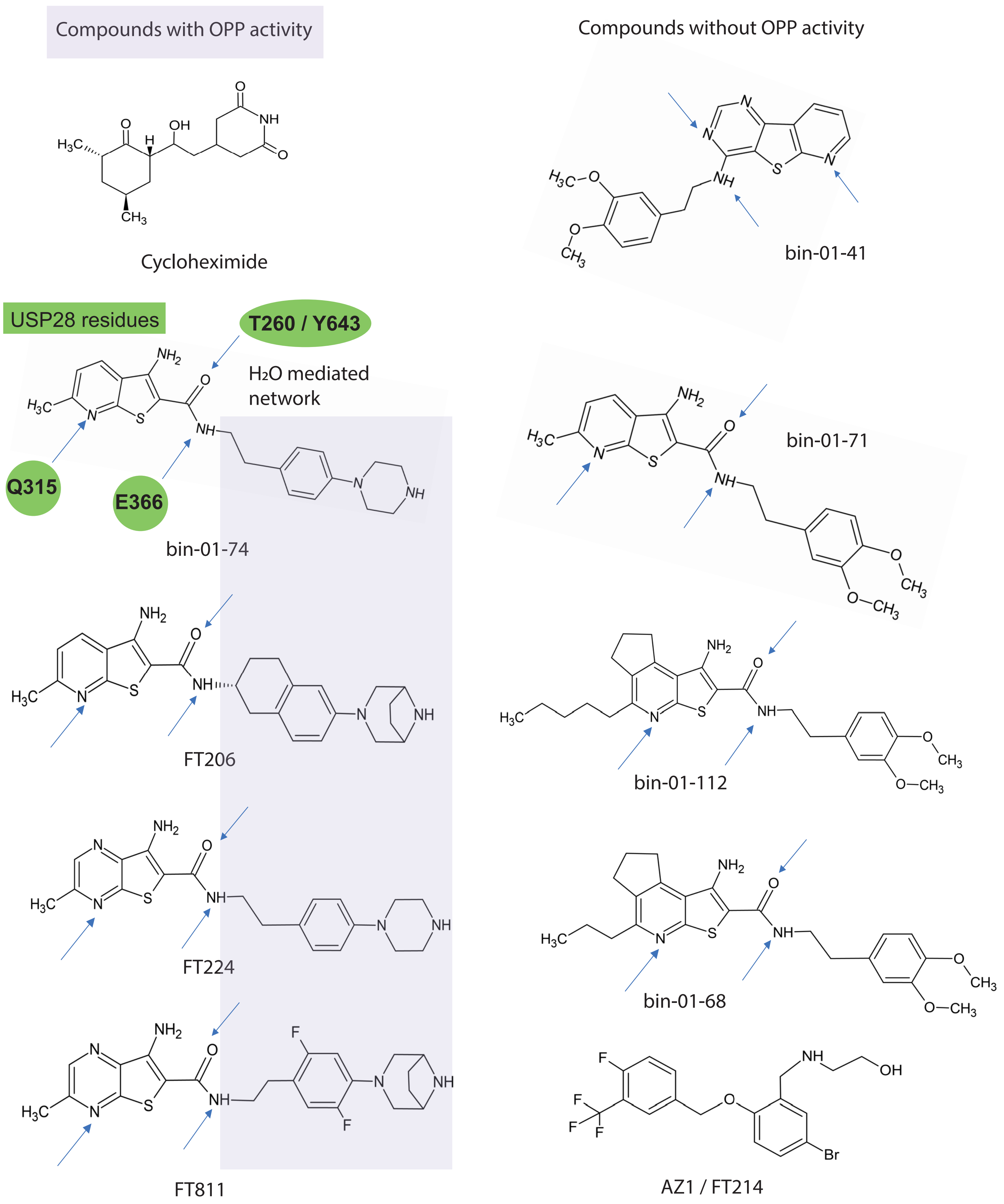
Compounds with OPP activity
Compounds without OPP activity
Cycloheximide
bin-01-41
USP28 residues
T260 / Y643
H2O mediated network
Q315
E366
bin-01-74
bin-01-71
FT206
bin-01-112
FT224
bin-01-68
FT811
AZ1 / FT214

**Figure S12**

**a**

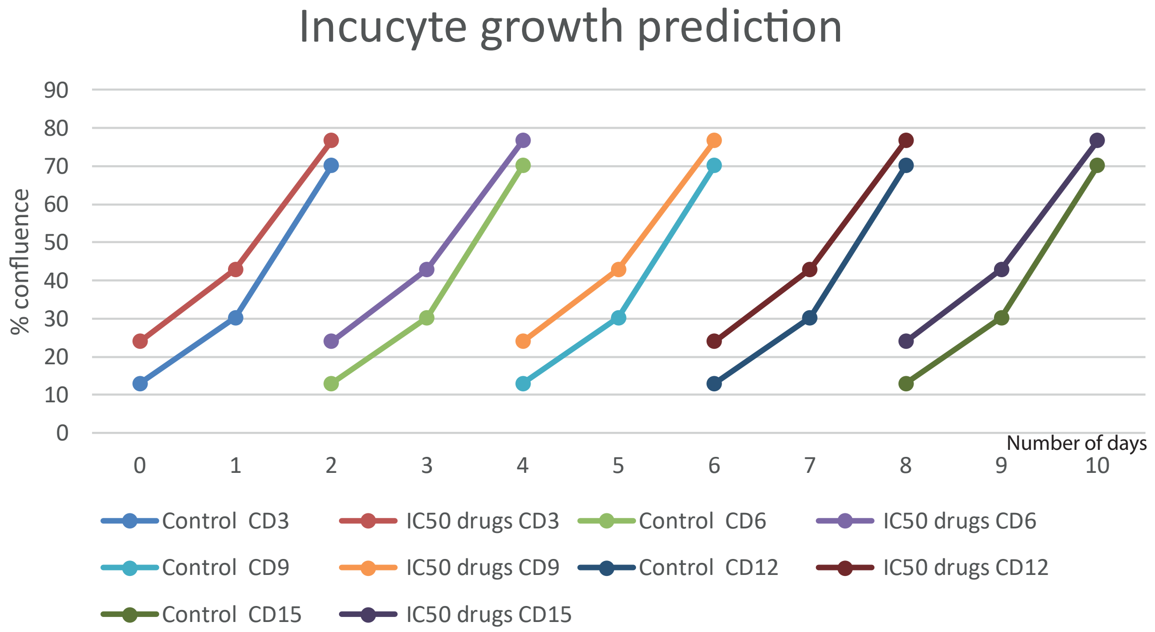


**b**

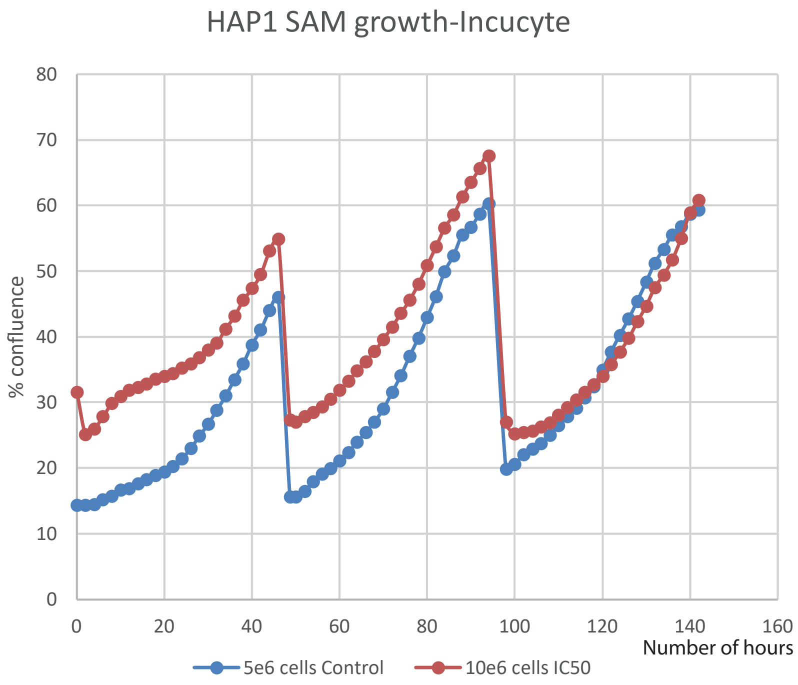


**c**

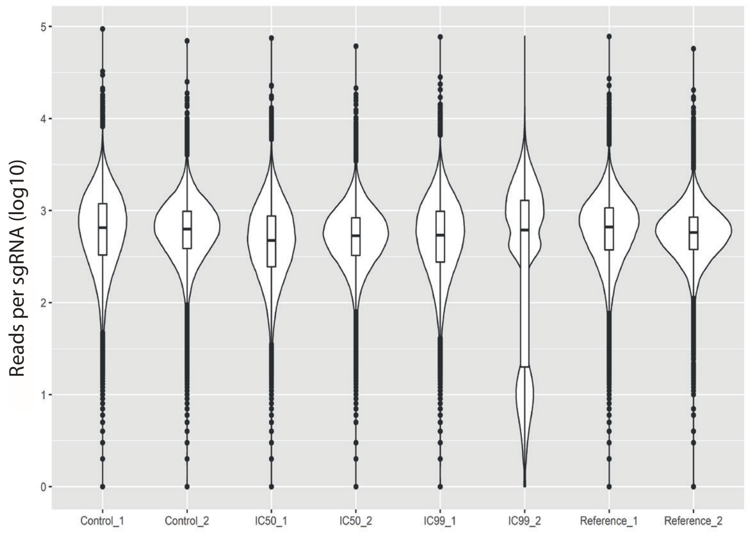


**d**

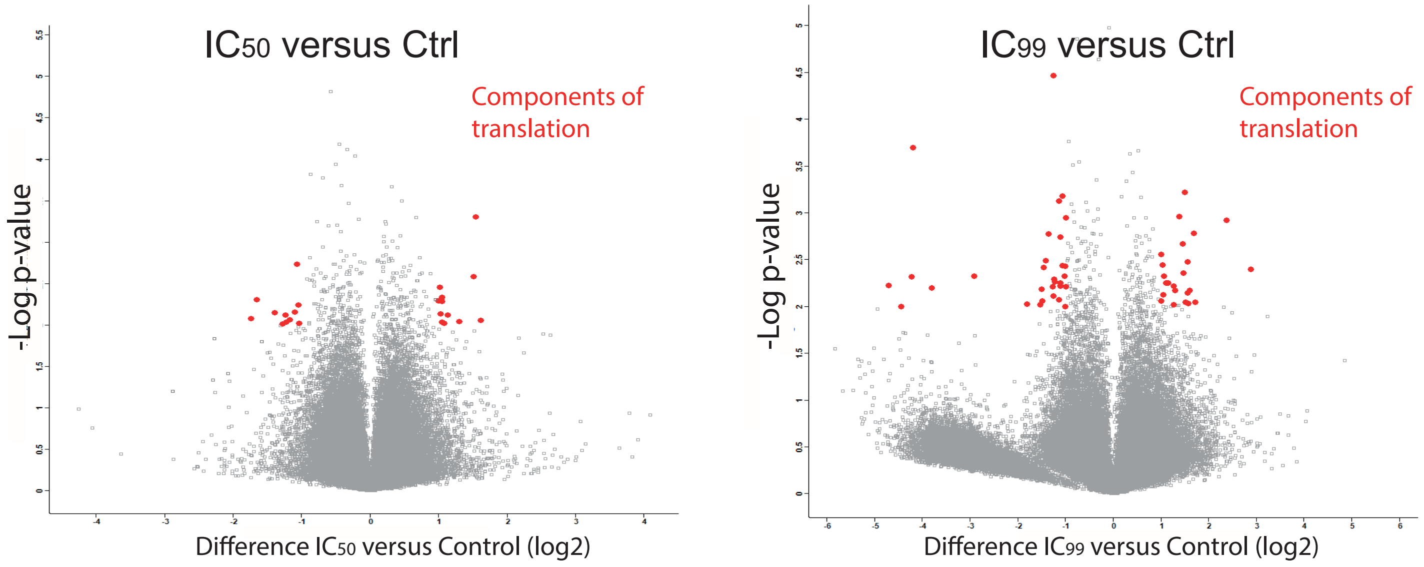

### Figure S13

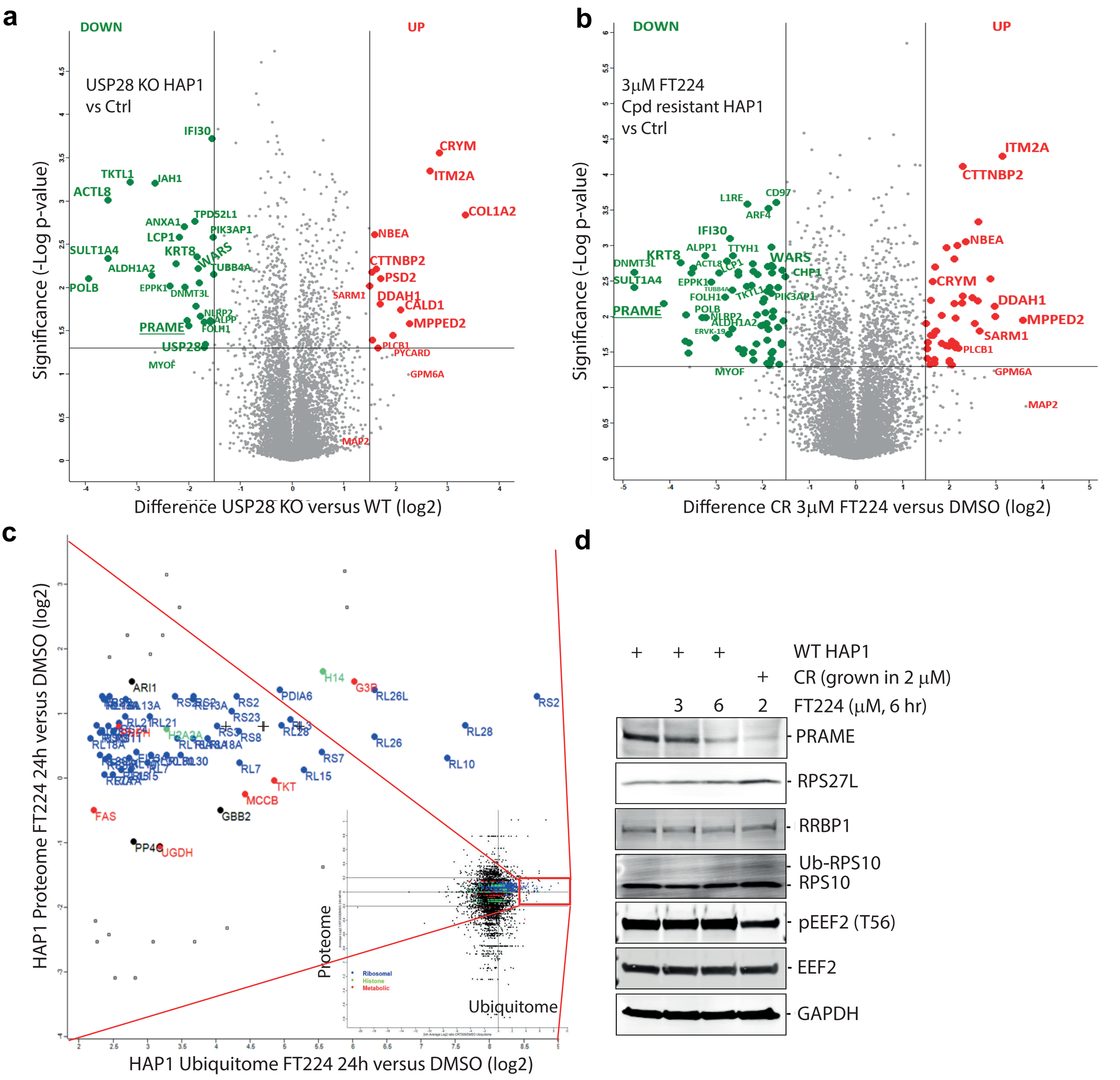


**e**

| Gene | DMSO_0 | DMSO_1 | DMSO_2 | X3uM_0 | X3uM_1 | X3uM_2 | X3uM_6H_ | X3uM_6H_ | X3uM_6H_ | X10uM_6I | X10uM_6I | X10uM_6H_2 |
|---|---|---|---|---|---|---|---|---|---|---|---|---|
| | HAP1 Parental | | | CR (3 μM) | | | Acute (10 μM, 6 hr) | | | CR (10 μM, 6 hr) | | |
| USP28 | 17.8965 | 15.6793 | 16.7918 | 17.9614 | 17.9998 | 19.2743 | 17.4413 | 18.3694 | 17.9934 | 16.8438 | 15.2543 | 16.5302 |
| USP25 | 17.6482 | 14.4997 | 15.4462 | 11.1815 | 11.7475 | 11.3501 | 17.1394 | 18.7551 | 17.1612 | 13.7692 | 11.9734 | 12.2887 |
| MYC | 232.185 | 264.096 | 245.87 | 134.756 | 129.177 | 134.575 | 190.928 | 179.939 | 194.227 | 115.223 | 141.254 | 120.499 |
| KDM1A | 141.202 | 127.608 | 134.255 | 88.8332 | 92.6161 | 90.137 | 117.047 | 120.763 | 114.475 | 77.8439 | 73.0569 | 73.7509 |
| CCND1 | 6.32293 | 6.67791 | 6.29828 | 0.157221 | 0.251844 | 0.209278 | 7.83609 | 7.85377 | 8.28466 | 0.09921 | 0.133723 | 0.07691 |
| CCND2 | 6.50055 | 6.04979 | 6.10736 | 0.352053 | 0.52889 | 0.359043 | 9.525 | 10.5619 | 8.34868 | 0.342 | 0.325545 | 0.244086 |
| CCND2-AS1 | 0.177174 | 0.789349 | 0.159771 | 0.08245 | 0 | 0.156102 | 0.719827 | 0.791217 | 0.552077 | 0 | 0 | 0.089223 |
| CCNE1 | 20.5395 | 19.7792 | 21.2569 | 11.2769 | 12.1786 | 12.976 | 17.857 | 18.2315 | 18.1138 | 16.0559 | 16.9953 | 15.7035 |
| CCNE2 | 15.5291 | 10.8741 | 12.8071 | 24.2588 | 24.4511 | 24.3571 | 19.8338 | 22.6567 | 21.3978 | 36.2913 | 30.4193 | 34.1569 |
| TP53 | 62.4232 | 60.2231 | 60.8689 | 91.9959 | 88.8853 | 94.5538 | 76.1774 | 77.5275 | 75.0685 | 105.85 | 112.101 | 106.809 |
| CDKN1A | 3.28262 | 3.67907 | 3.69081 | 79.6252 | 105.666 | 56.0279 | 6.87506 | 7.21073 | 6.48139 | 167.517 | 208.311 | 144.712 |
| HIF1A | 38.25 | 31.9784 | 35.2071 | 68.4307 | 70.214 | 67.966 | 39.7222 | 43.9223 | 39.4207 | 93.707 | 84.7966 | 85.3878 |
| XBP1 | 66.1002 | 57.6693 | 62.9502 | 105.802 | 109.442 | 99.7244 | 84.0191 | 86.8738 | 86.7626 | 49.5018 | 45.5466 | 44.4802 |
| CHAC1 | 1.61204 | 2.44629 | 1.82051 | 0.76482 | 0.851241 | 0.770566 | 11.0339 | 8.20047 | 11.0733 | 0.116231 | 0.2215 | 0.30205 |
| DDIT3 | 19.4157 | 16.1869 | 18.8676 | 49.0905 | 48.5859 | 49.0305 | 54.186 | 53.9276 | 52.21 | 31.354 | 30.8316 | 31.464 |
| HSPA8 | 556.954 | 477.789 | 514.641 | 768.368 | 709.269 | 786.469 | 121.057 | 121.283 | 121.783 | 223.53 | 205.907 | 236.155 |
| OSTN | 0 | 0 | 0 | 10.6935 | 9.65576 | 10.8476 | 0 | 0 | 0 | 8.06364 | 7.17718 | 6.77668 |
| EPHA5 | 0.029327 | 0.010604 | 0.025116 | 17.7319 | 18.0287 | 17.8888 | 0.031768 | 0.057583 | 0.020993 | 15.0038 | 12.0593 | 15.745 |
| MAP2 | 0.178344 | 0.167965 | 0.17413 | 33.4096 | 32.0515 | 34.6706 | 0.189575 | 0.163597 | 0.182211 | 37.4596 | 33.5677 | 31.3784 |
| ABCB1 | 1.75532 | 1.53722 | 1.8244 | 56.3755 | 56.9954 | 52.2828 | 2.95303 | 3.14748 | 3.10007 | 74.7243 | 72.0455 | 72.006 |
| RPS27A | 1032.36 | 1006.35 | 1027.16 | 1171.82 | 976.214 | 1036.8 | 1015.2 | 1133.02 | 1125.7 | 1033.14 | 965.163 | 1198.25 |

mRNA transcript levels (log2)

Cyclin D1, D2 - upregulated in G1-phase

Cyclin E1, E2 - upregulated in S-phase

Unfolded protein response (UPR)

Processing newly synthesized proteins
Regulator of dendritic growth
Axon guidance during development
Microtubule-assoc. protein
Multidrug resistance pump

**Figure S14**

a

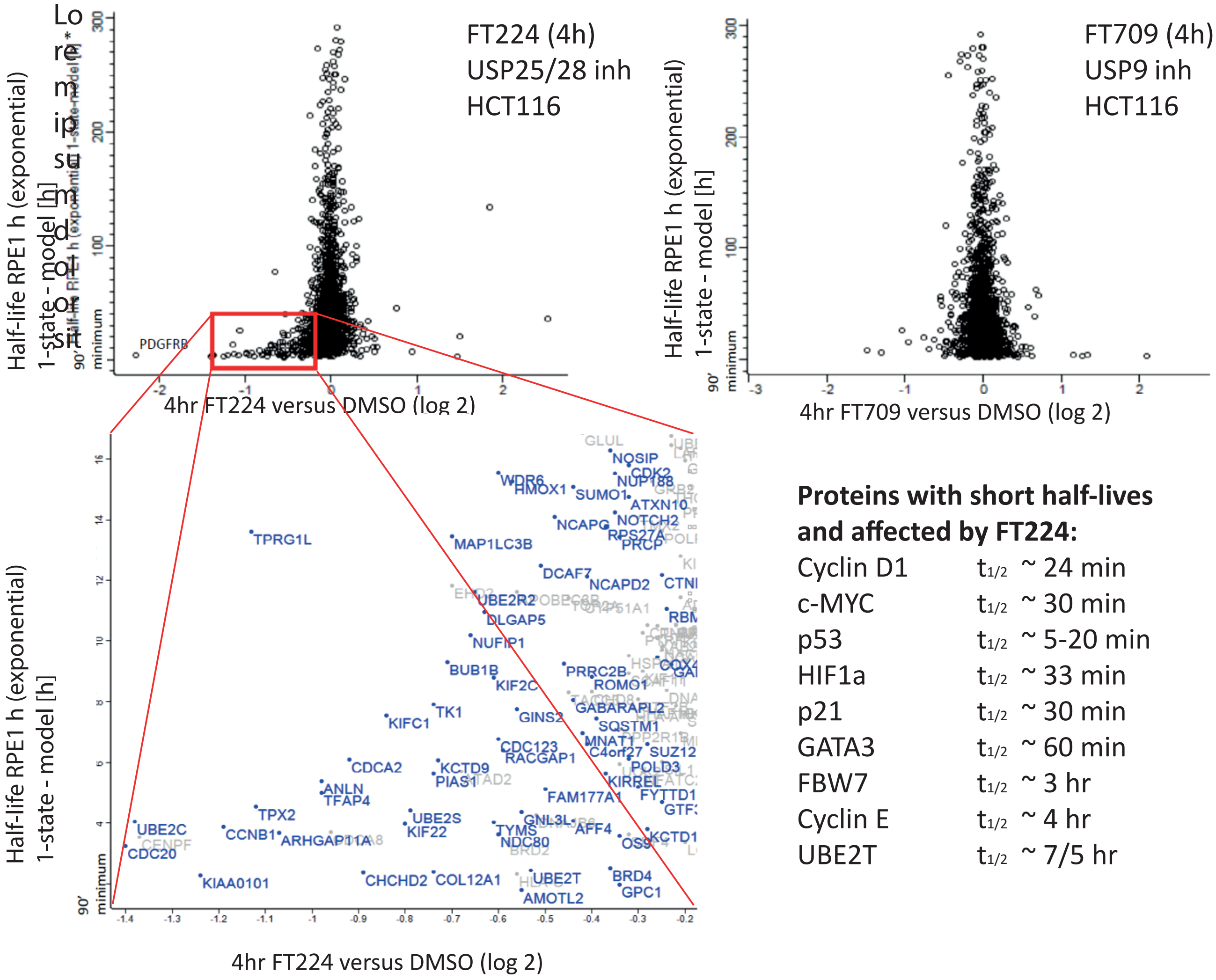


**Proteins with short half-lives and affected by FT224:**

| | |
|---|---|
| Cyclin D1 | $t_{1/2}$ ~ 24 min |
| c-MYC | $t_{1/2}$ ~ 30 min |
| p53 | $t_{1/2}$ ~ 5-20 min |
| HIF1a | $t_{1/2}$ ~ 33 min |
| p21 | $t_{1/2}$ ~ 30 min |
| GATA3 | $t_{1/2}$ ~ 60 min |
| FBW7 | $t_{1/2}$ ~ 3 hr |
| Cyclin E | $t_{1/2}$ ~ 4 hr |
| UBE2T | $t_{1/2}$ ~ 7/5 hr |

b

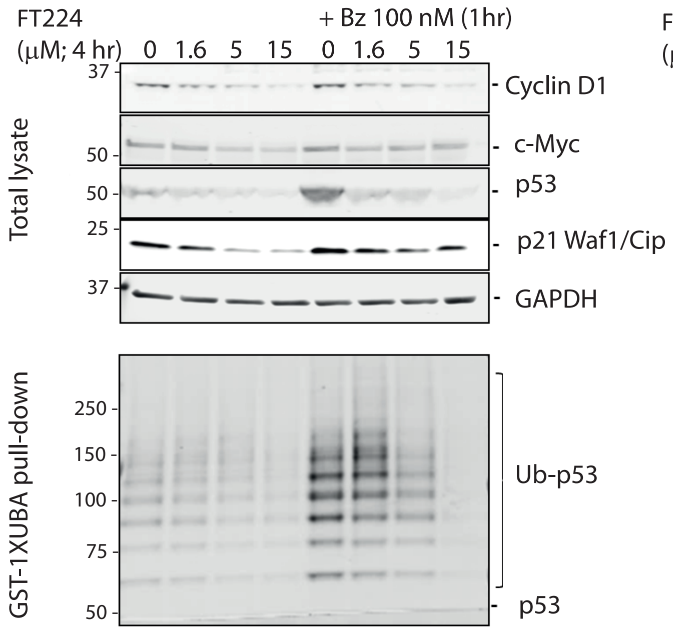


c

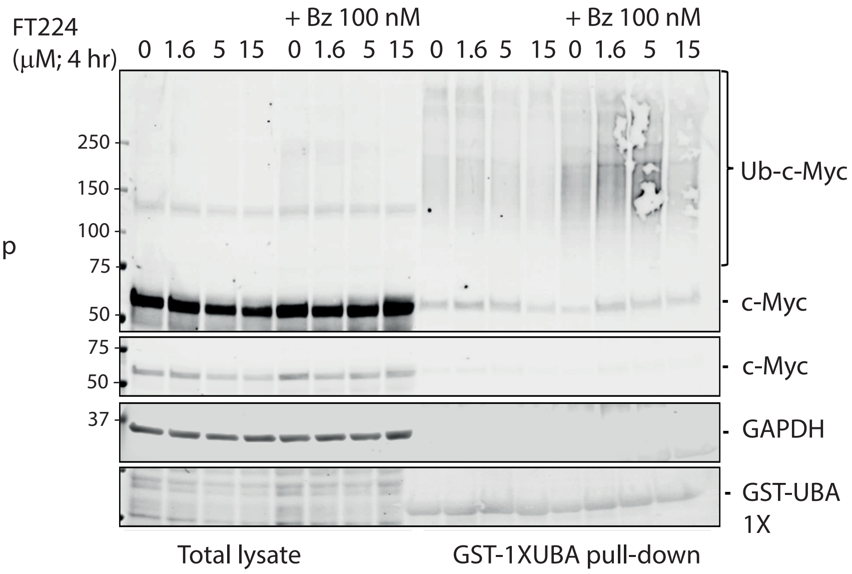


BZ = Bortezomib
CHX = Cycloheximide

### Figure S15

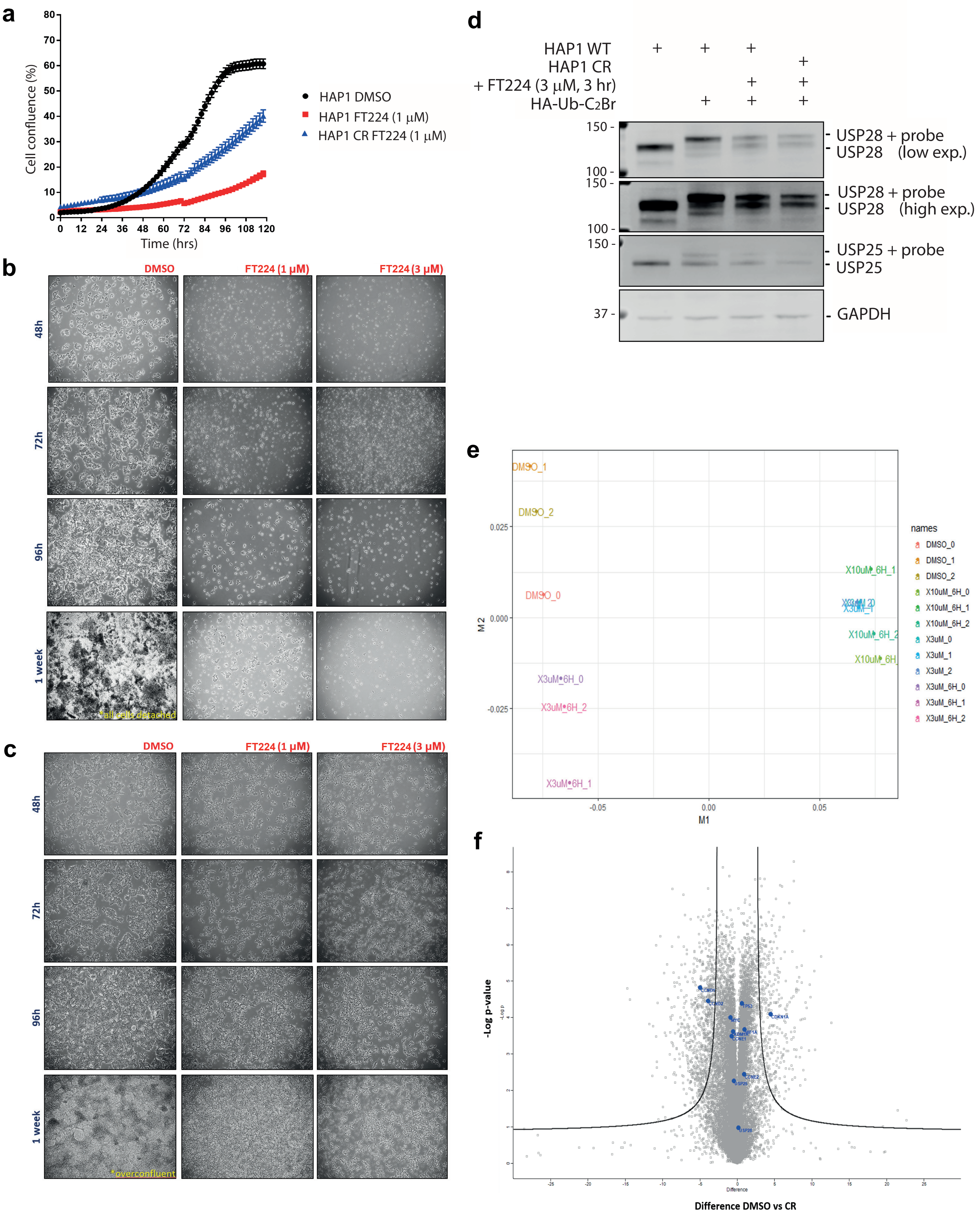
a
Cell confluence (%)
Time (hrs)
HAP1 DMSO
HAP1 FT224 (1 μM)
HAP1 CR FT224 (1 μM)
b
DMSO
FT224 (1 μM)
FT224 (3 μM)
48h
72h
96h
1 week
*all cells detached
c
DMSO
FT224 (1 μM)
FT224 (3 μM)
48h
72h
96h
1 week
*overconfluent
d
HAP1 WT
HAP1 CR
+ FT224 (3 μM, 3 hr)
HA-Ub-C2Br
USP28 + probe
USP28 (low exp.)
USP28 + probe
USP28 (high exp.)
USP25 + probe
USP25
GAPDH
e
names
DMSO_0
DMSO_1
DMSO_2
X10uM_6H_0
X10uM_6H_1
X10uM_6H_2
X3uM_0
X3uM_1
X3uM_2
X3uM_6H_0
X3uM_6H_1
X3uM_6H_2
M1
M2
f
-Log p-value
Difference DMSO vs CR

### Figure S16

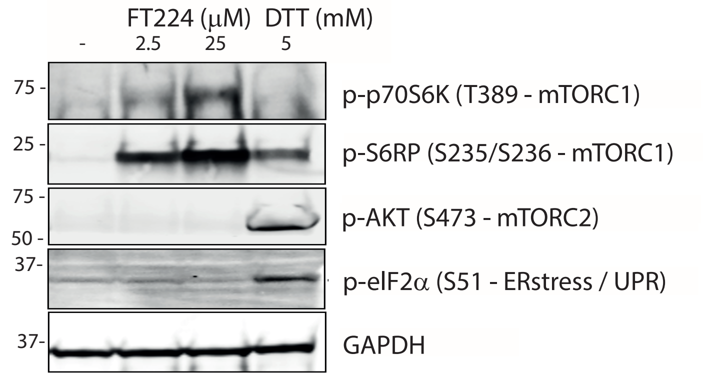


### Figure S17

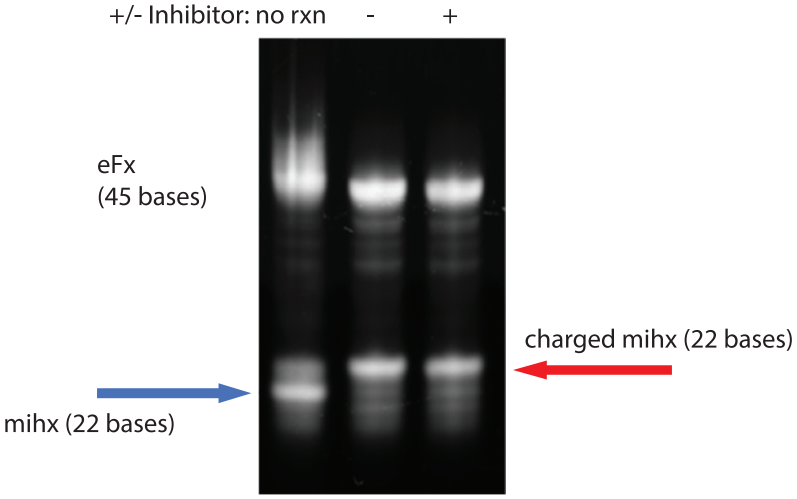

Uncropped blots for Figure 1c

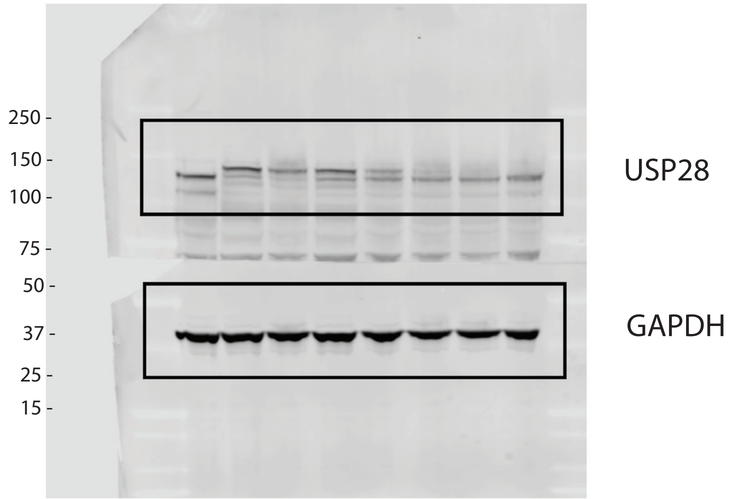


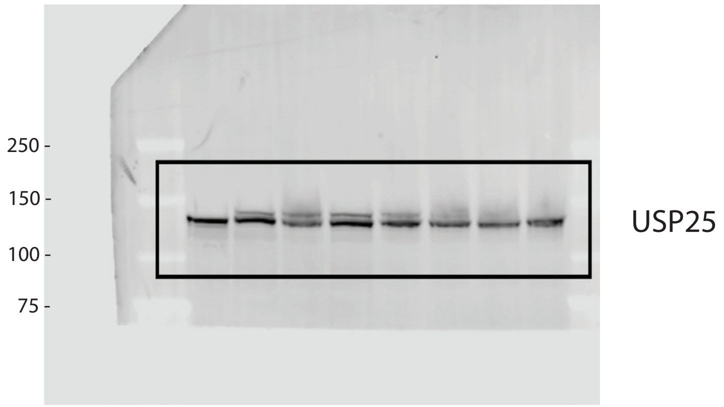

### Figure S19

Uncropped blots for Figure 2c (1)

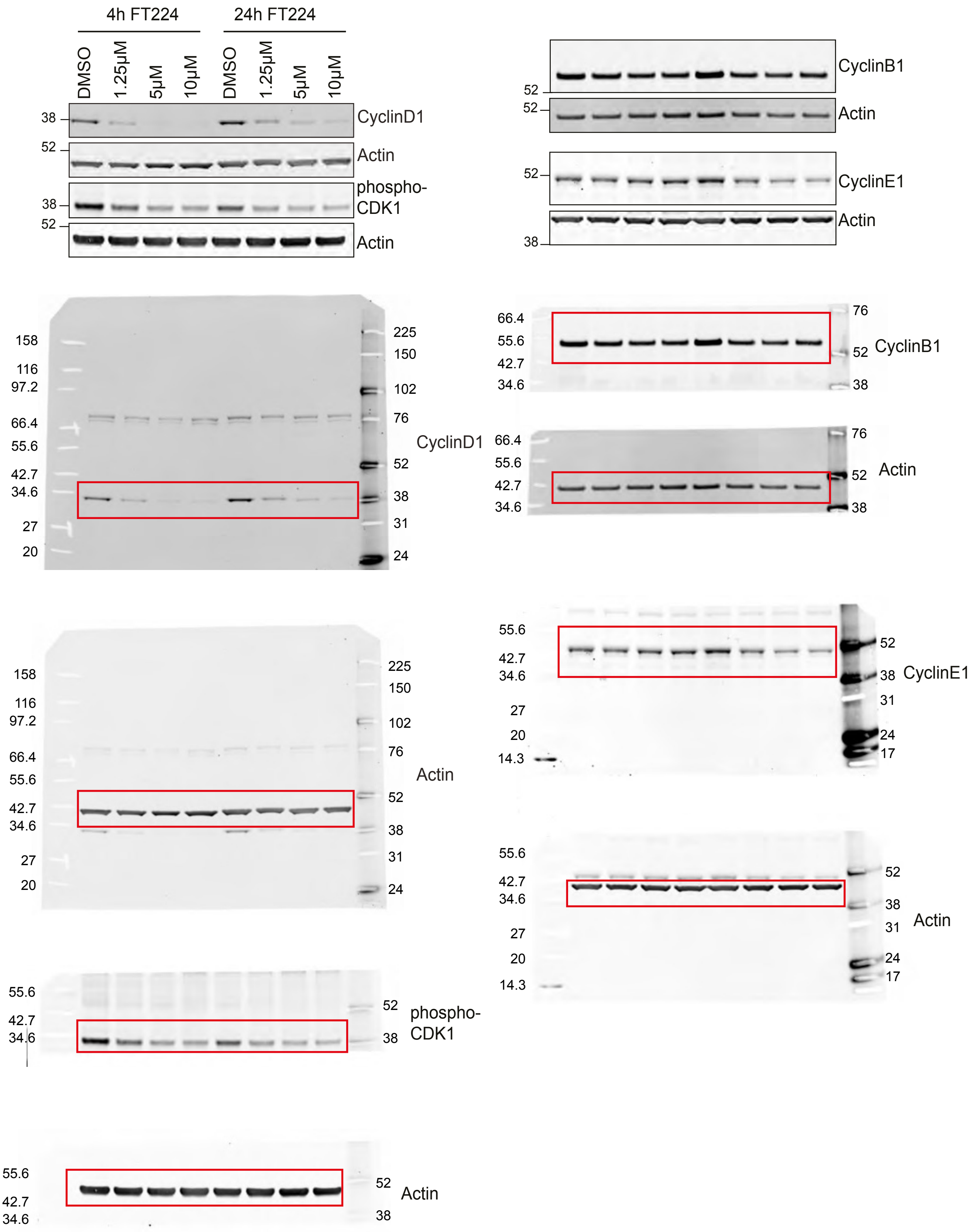

Uncropped blots for Figure 2c (2)

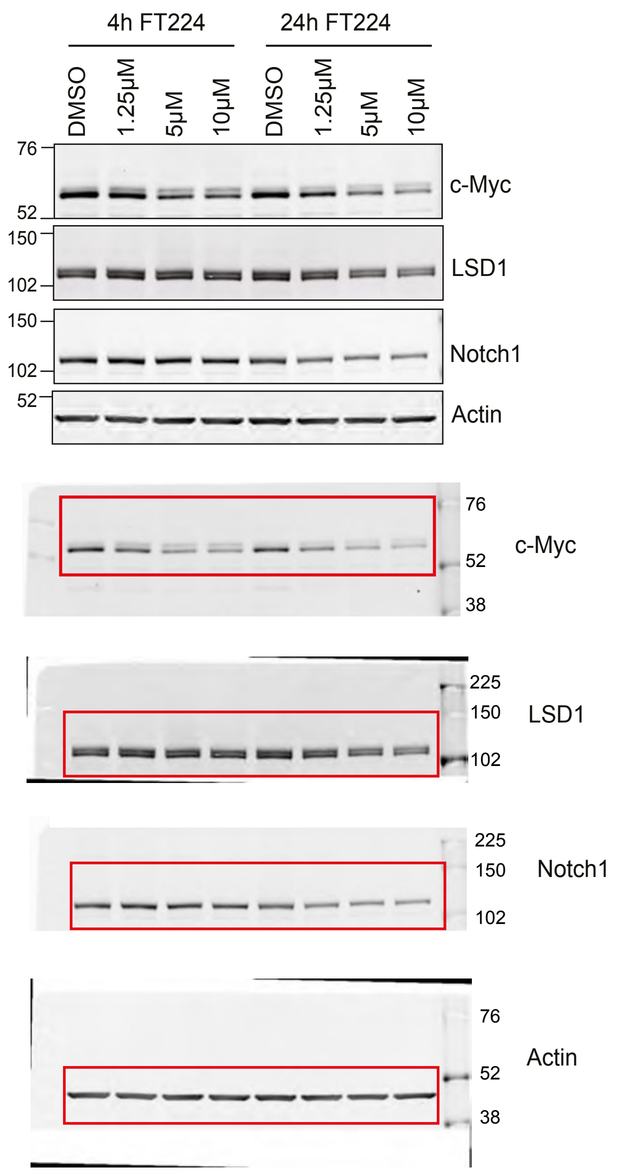

Figure S20

Uncropped blots for Figure 2d (1)

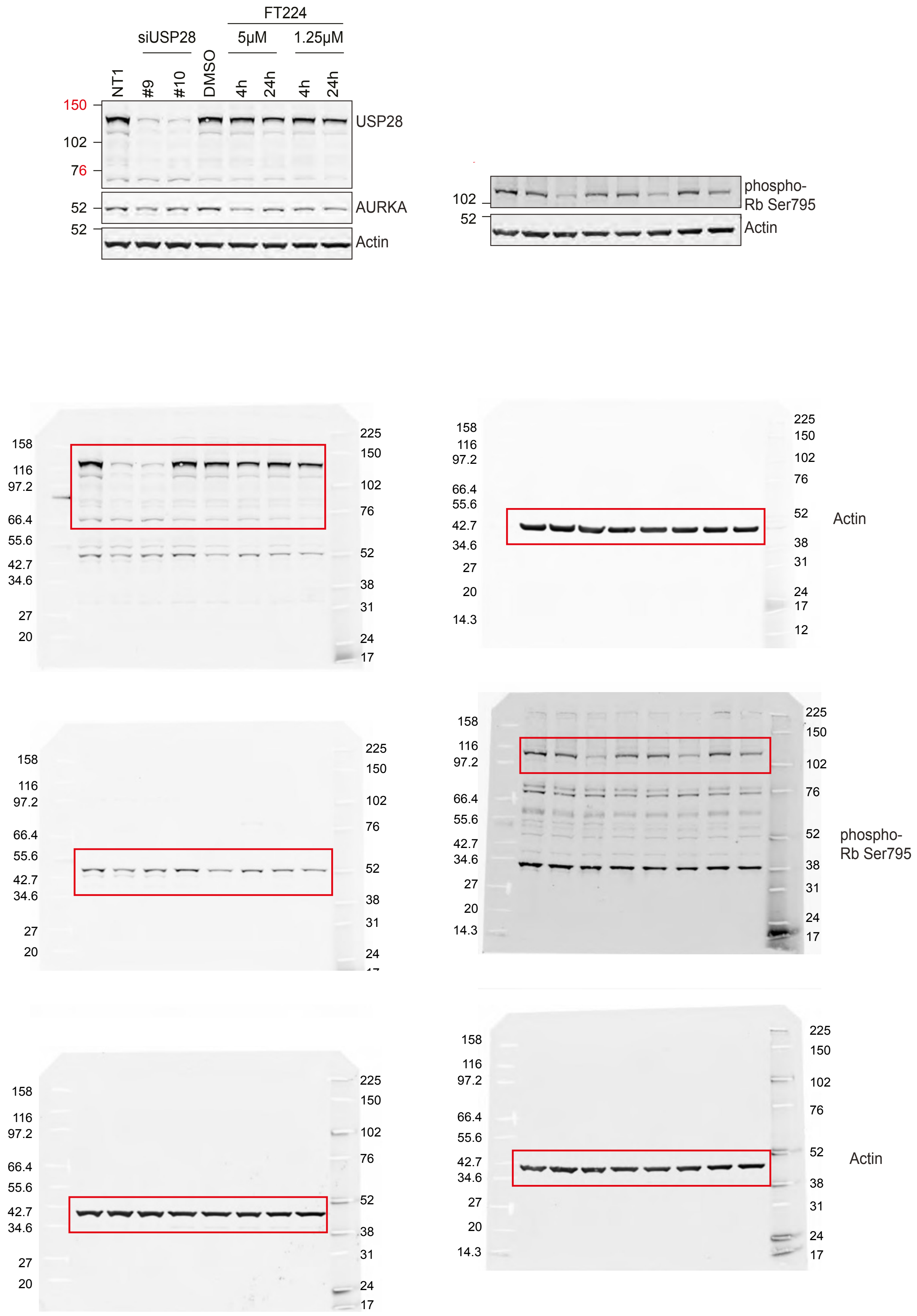

Uncropped blots for Figure 2d (2)

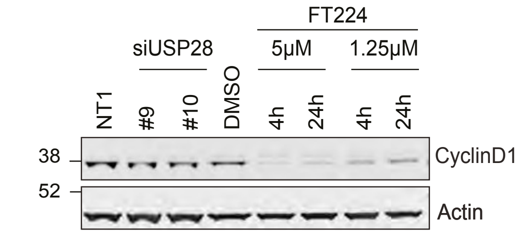


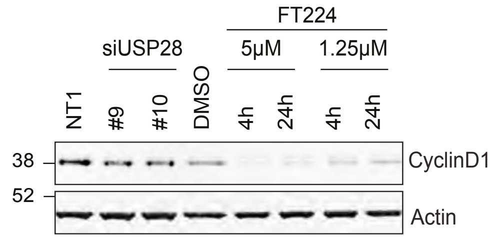


#### Gel 4

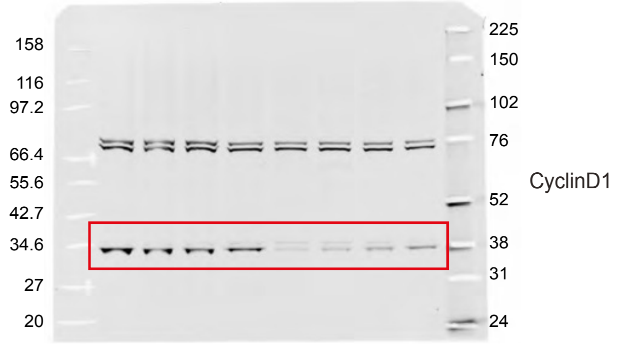


#### Alternative blot - Gel 24 - rerun samples

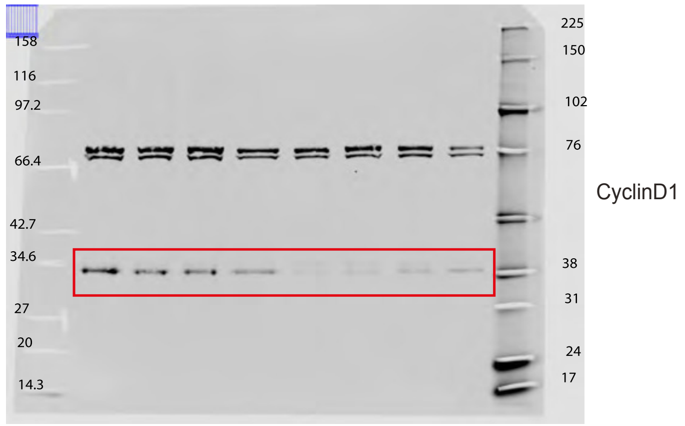


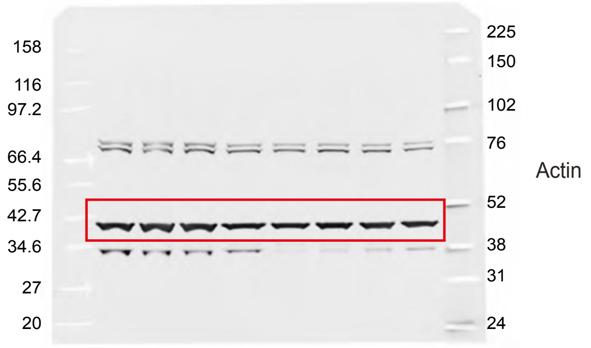


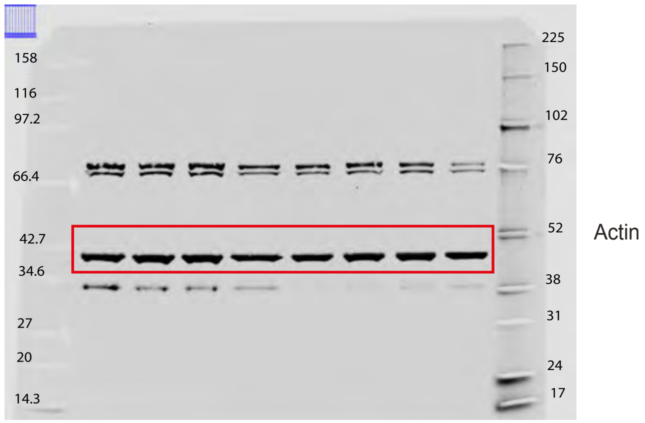

Uncropped blots for Figure 2f,g

Figure 2f

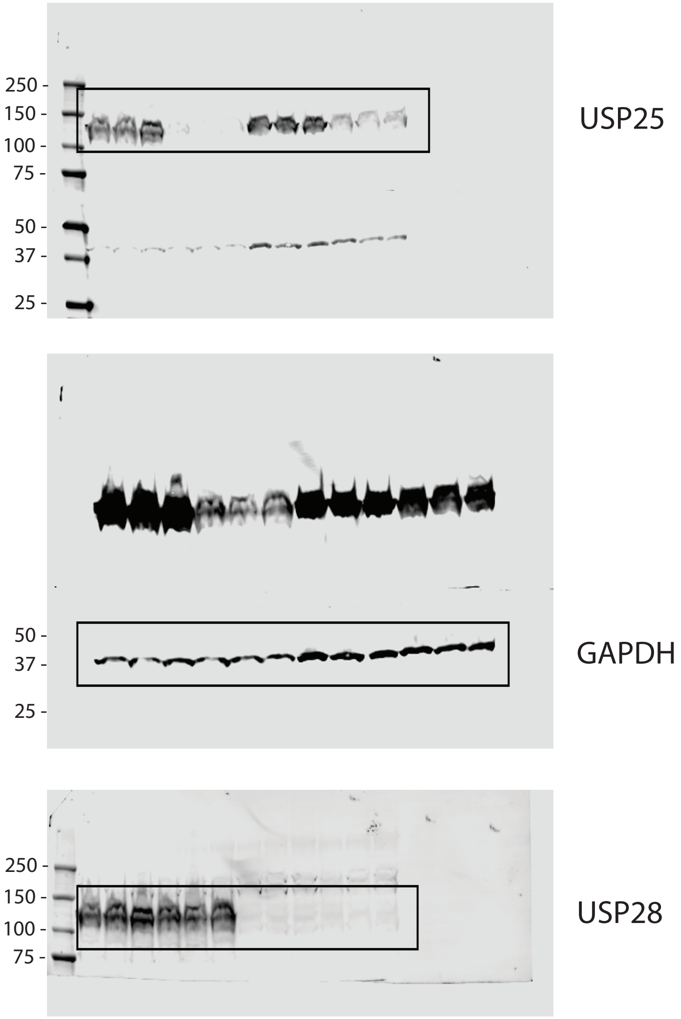


Figure 2g

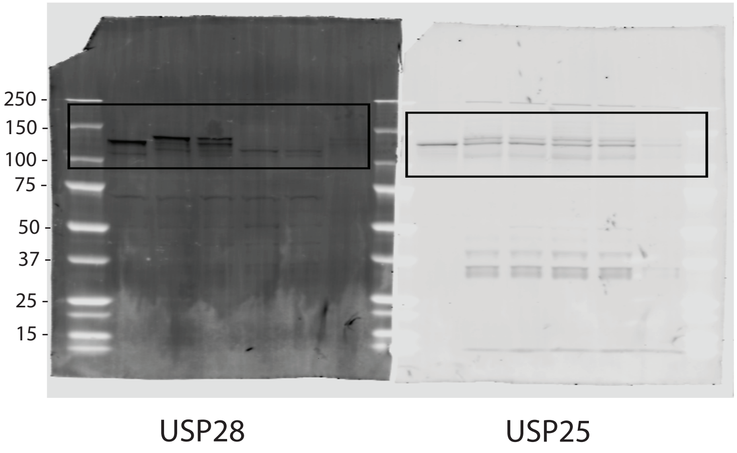

Uncropped blots for Figure 3a,c,d

Figure 3a

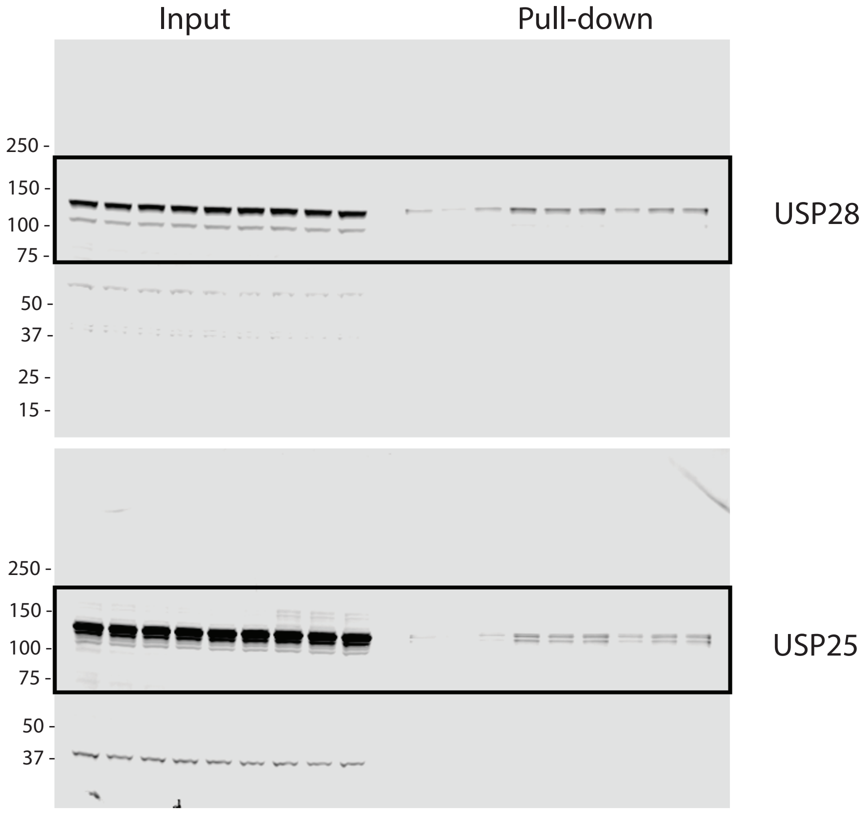


Figure 3c

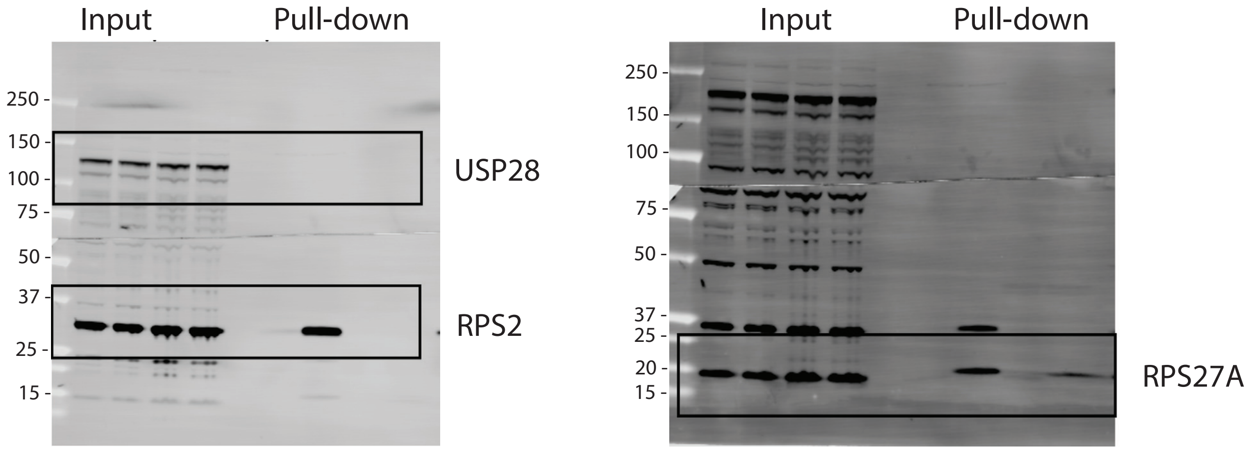


Figure 3d

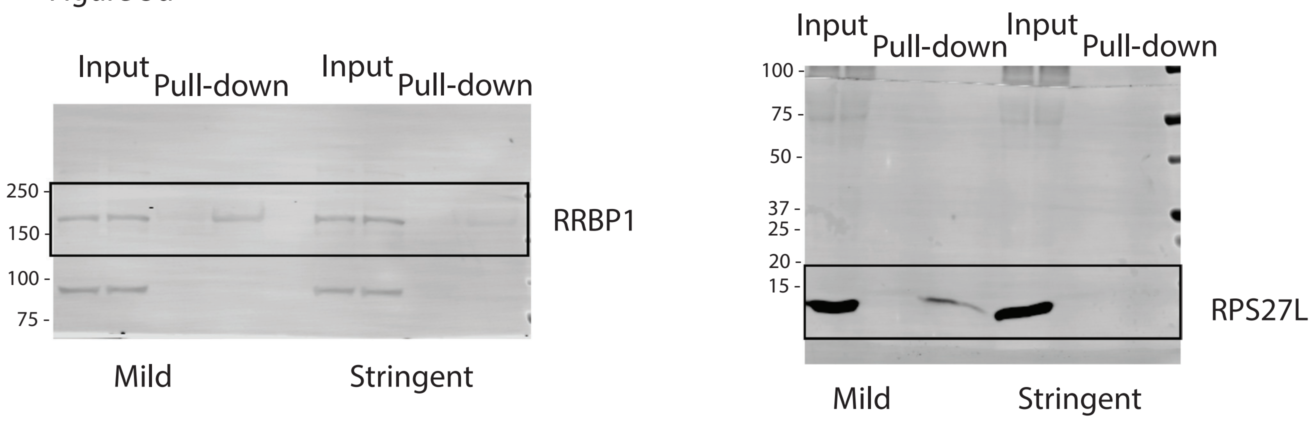

**Figure S23**

Uncropped blots for Figure 4

Figure 4a

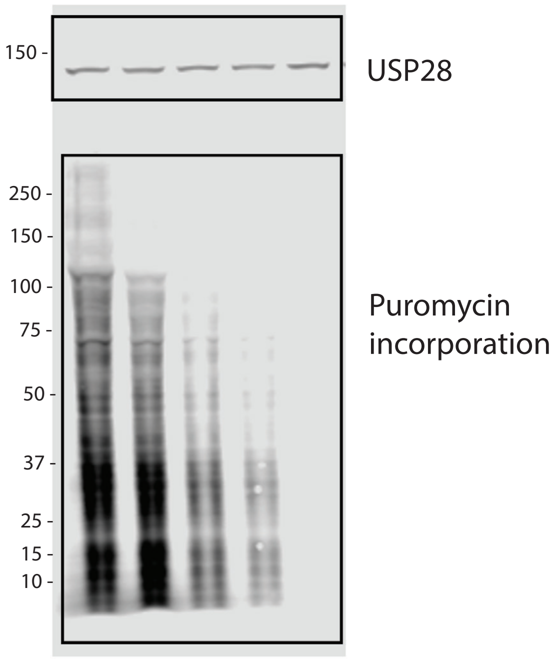


Figure 4b

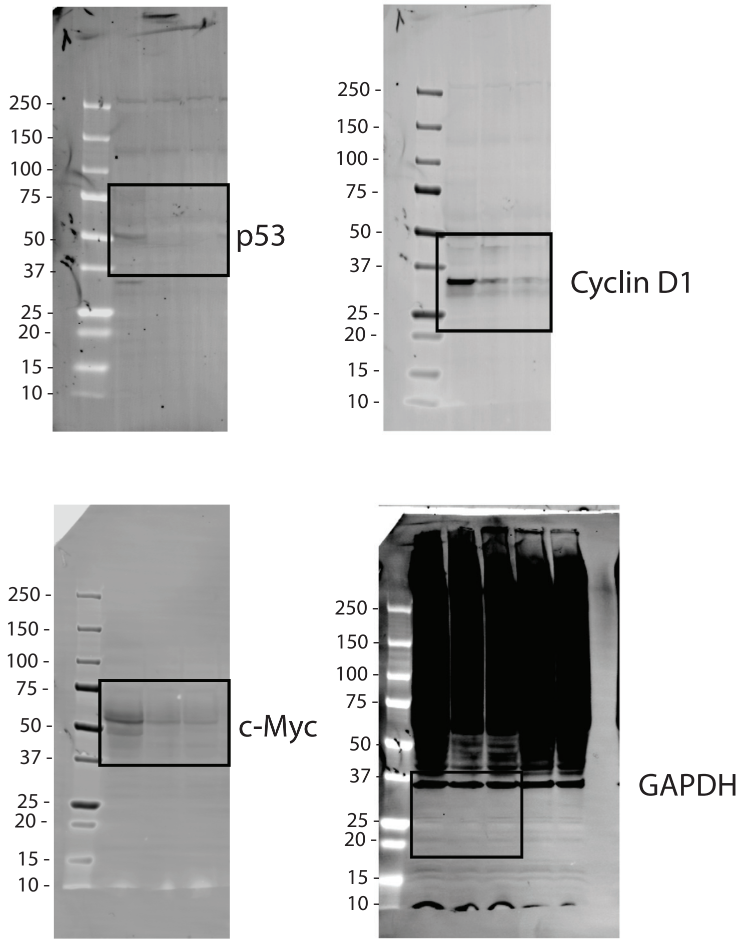


Figure 4f

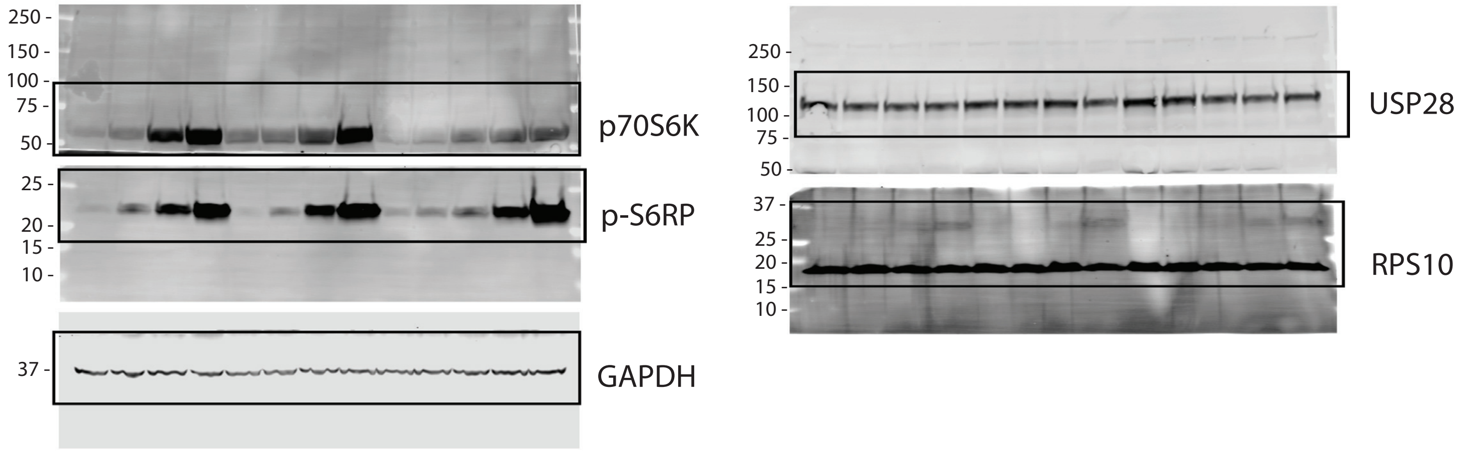

### Figure S24

Uncropped blots for Figure S3a,b

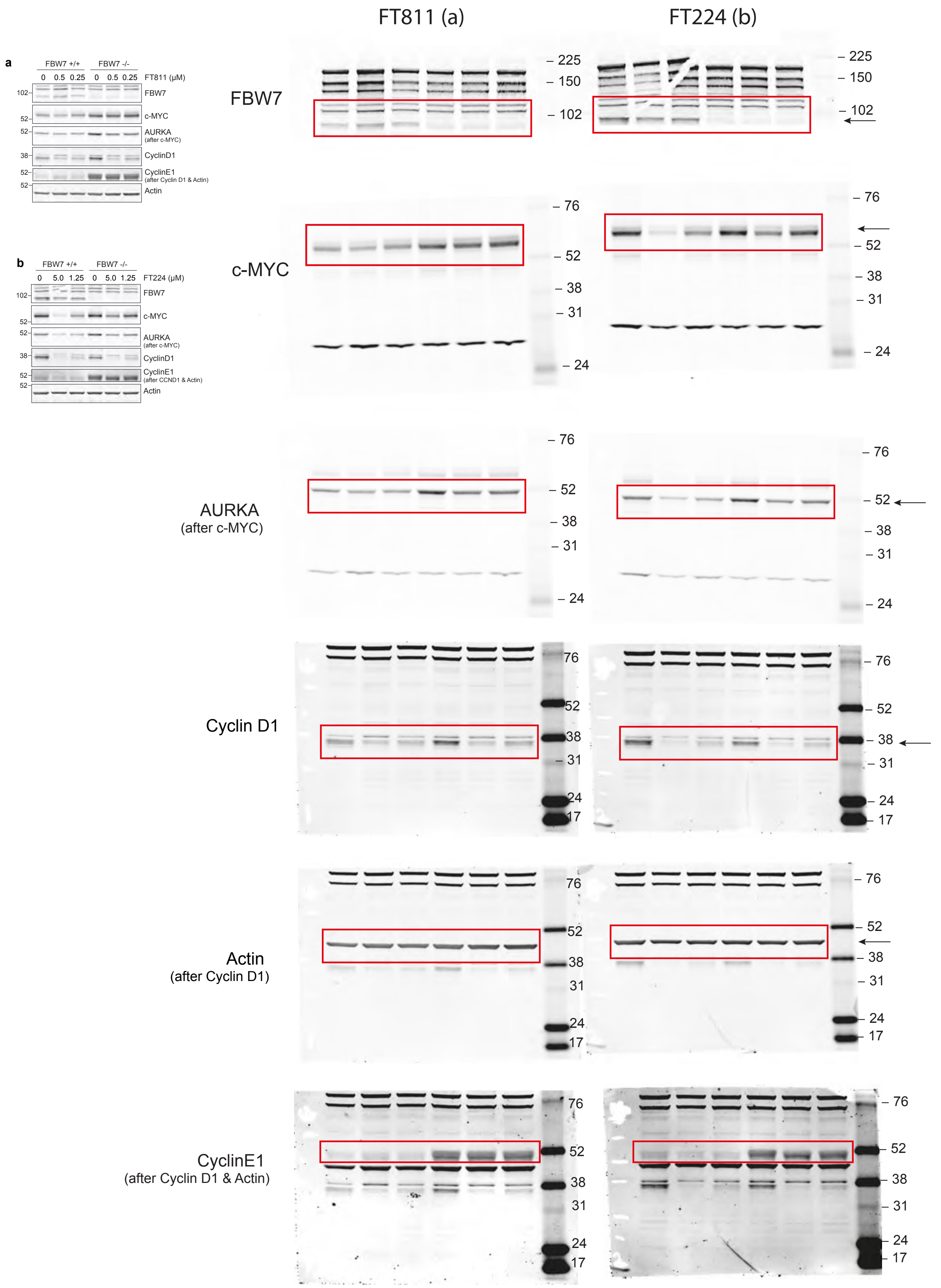

### Figure S25

Uncropped blots for Figure S3e,f

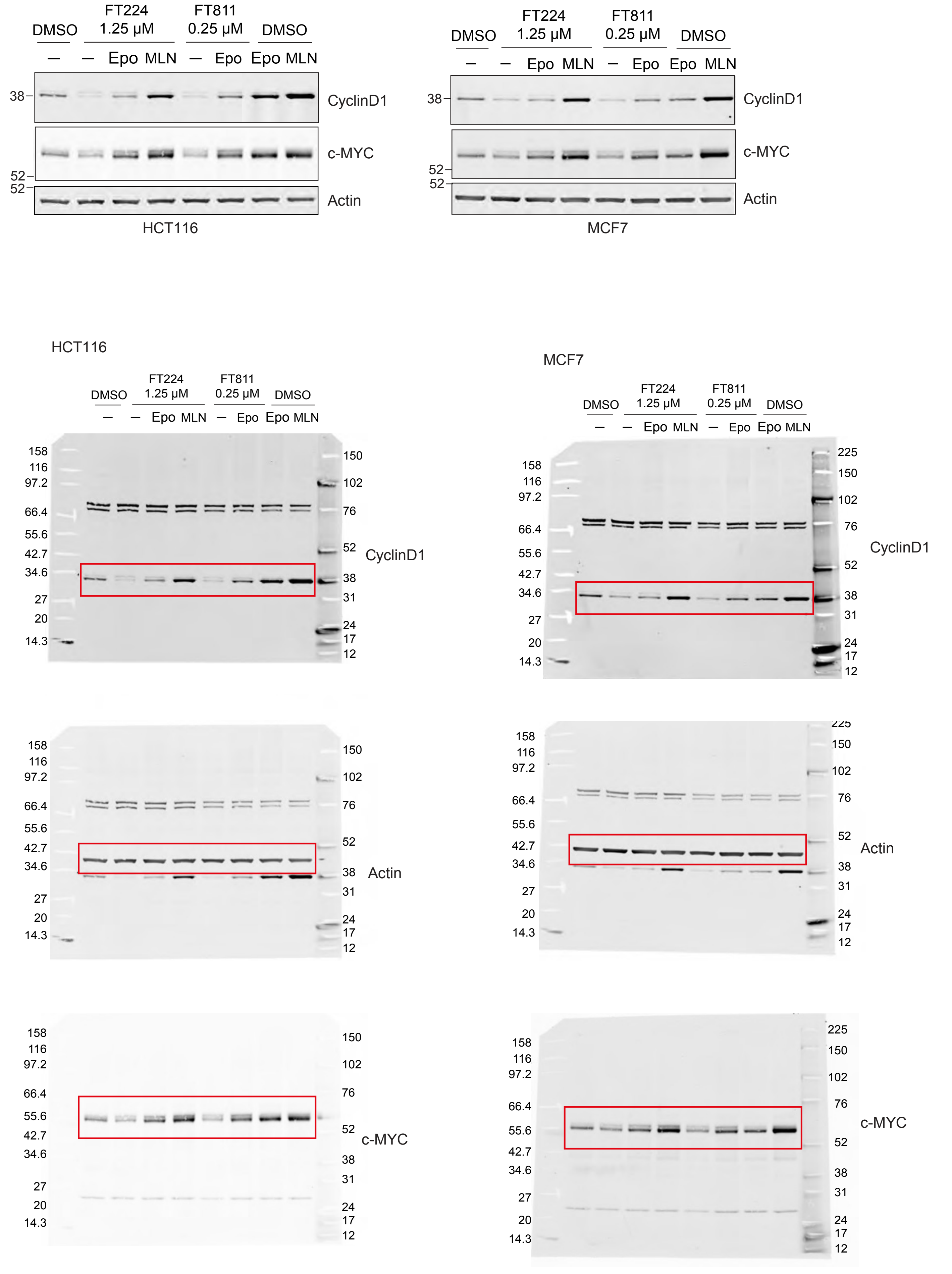

### Figure S26

Uncropped blots for Figure S4

#### Figure S4a

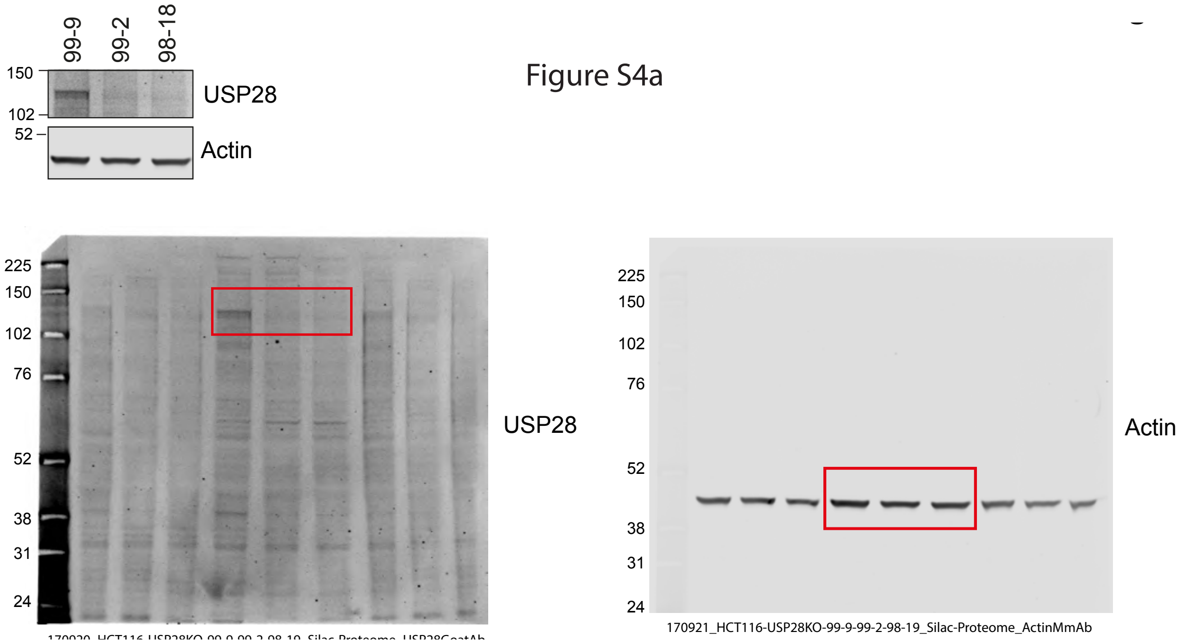


#### Figure S4b

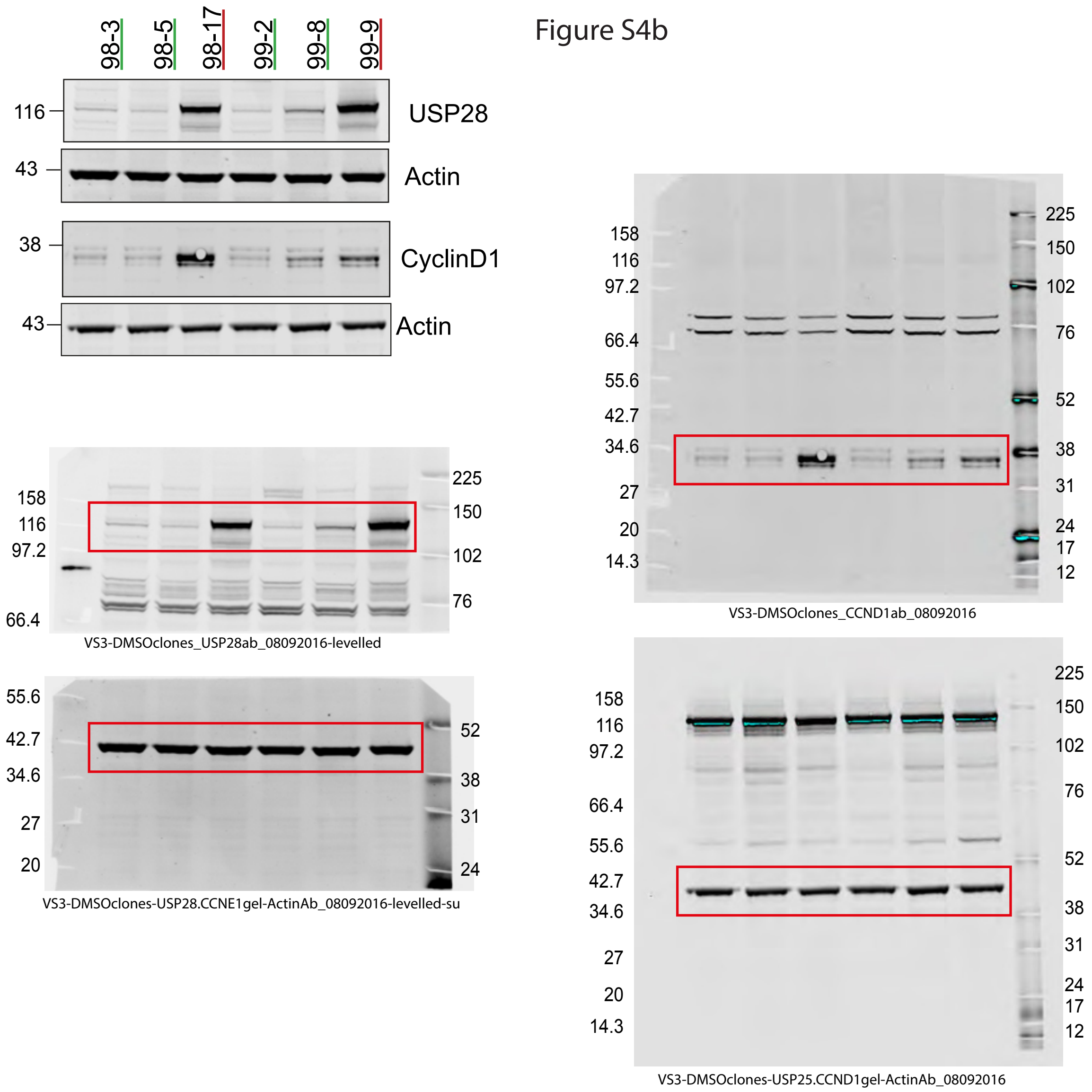

#### Figure S27

### Uncropped blots for Figure S5a, b (1)

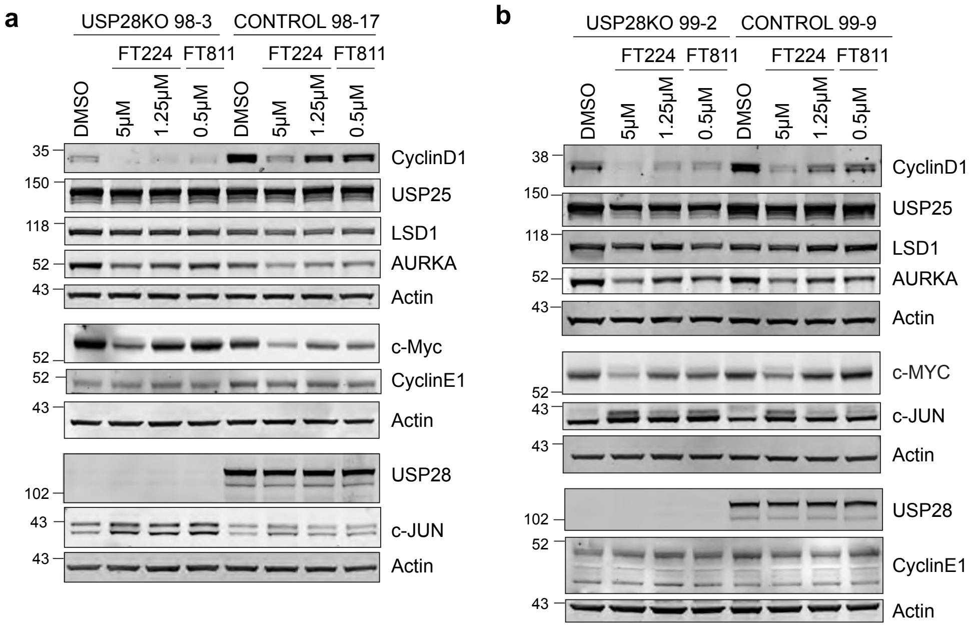


Figure S5a

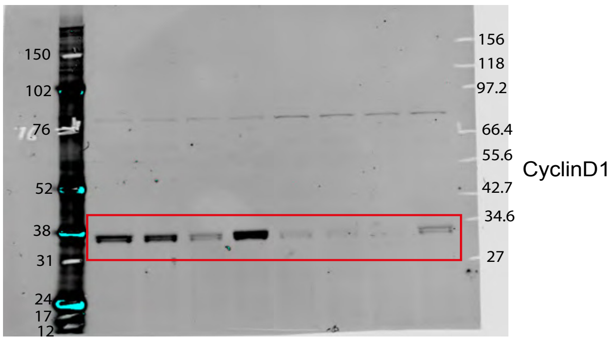


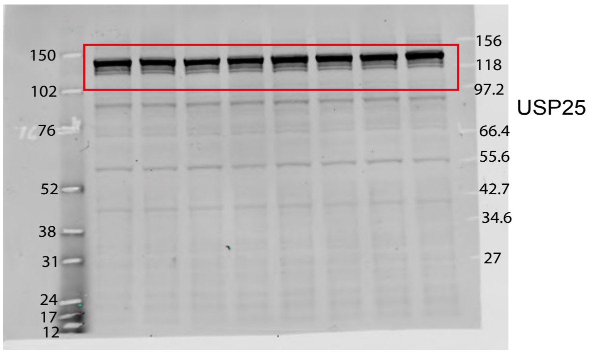


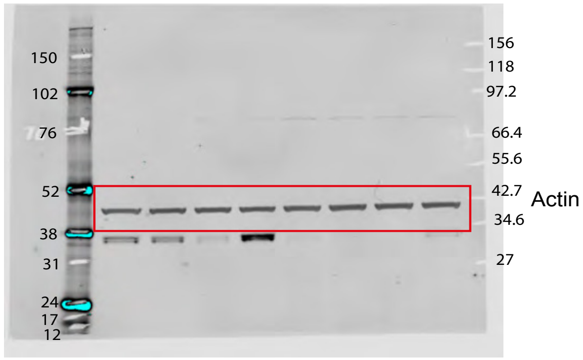


Flipped for figure

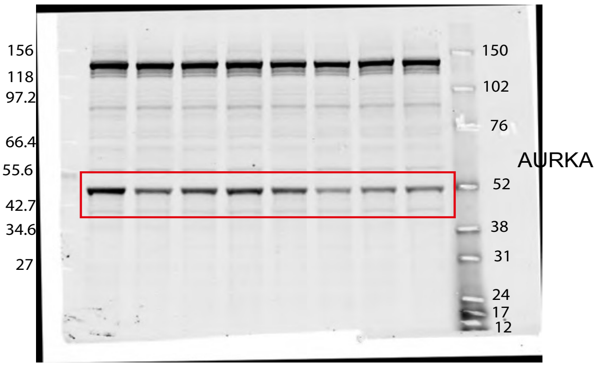


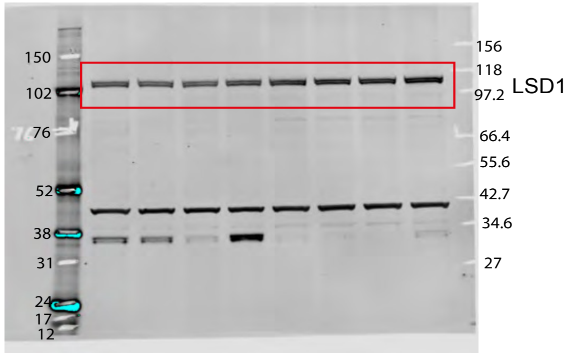


4;

### Uncropped blots for Figure S5a, b (2)

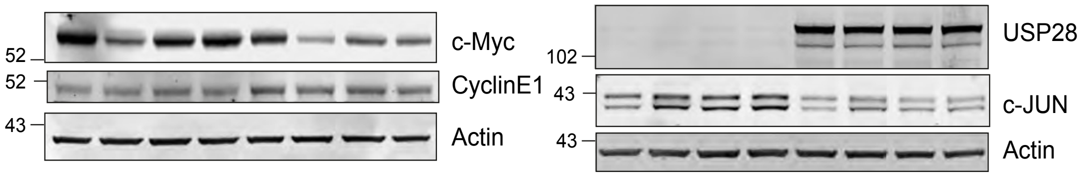


#### Figure S5a

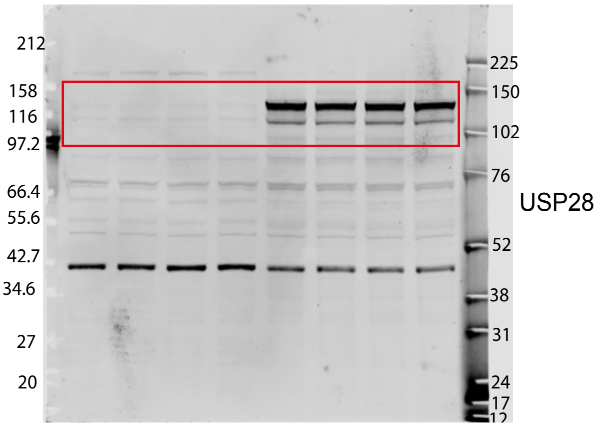


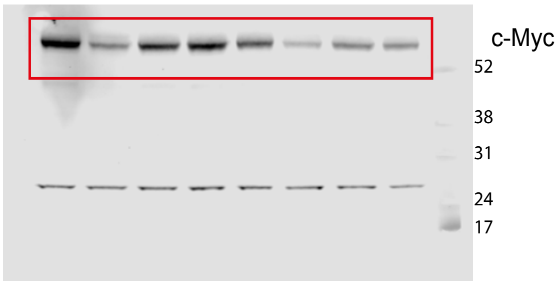


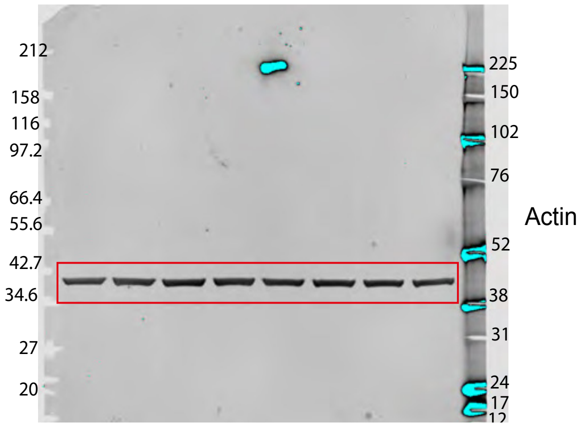


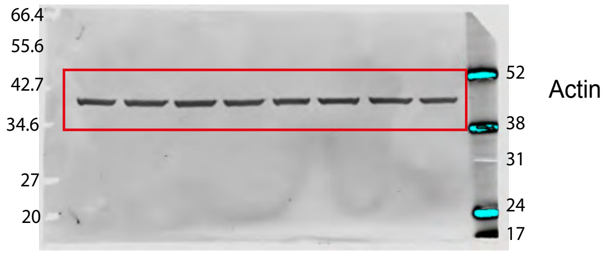


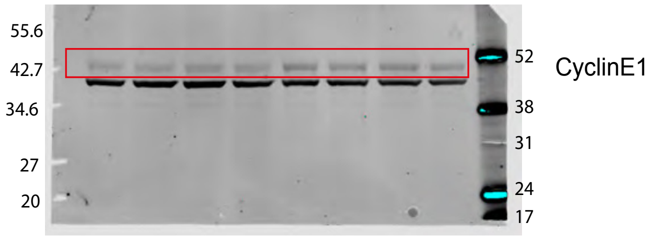


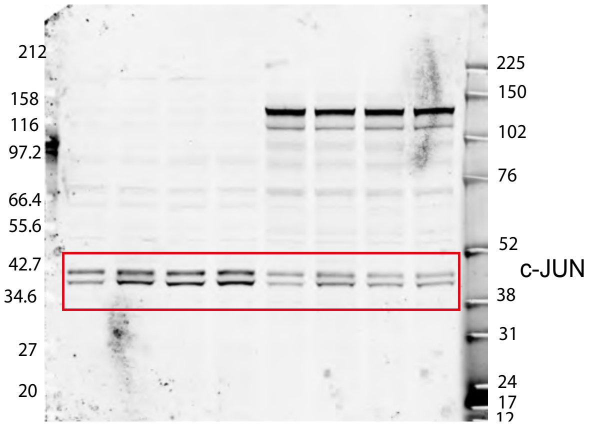

Uncropped blots for Figure S5a, b (3)

Figure S5b

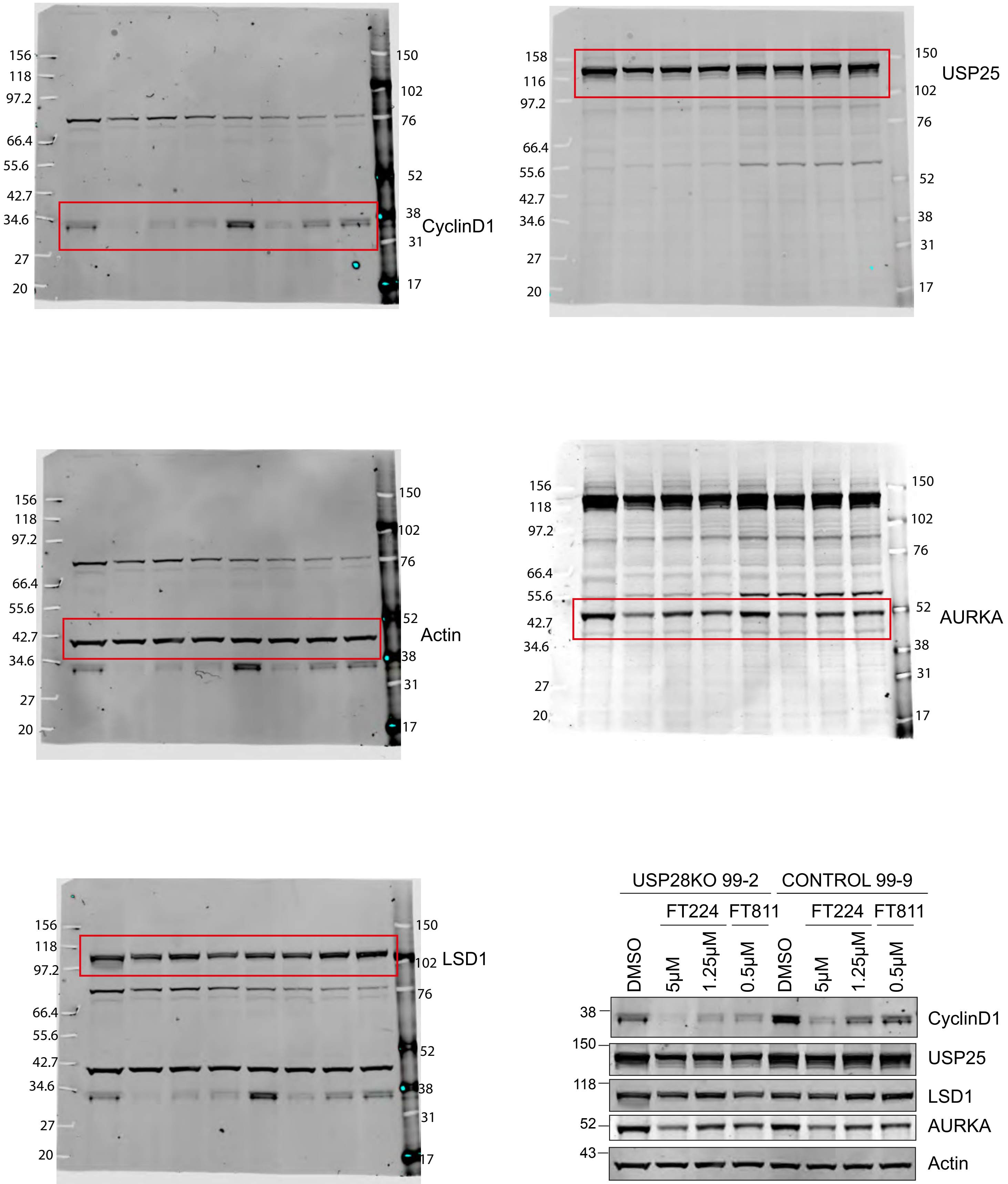

Uncropped blots for Figure S5a, b (4)

#### Figure S5b

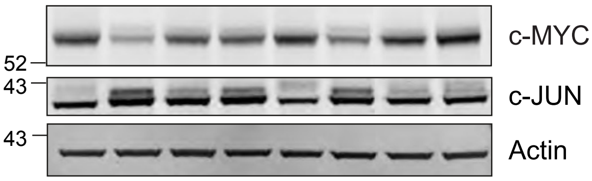


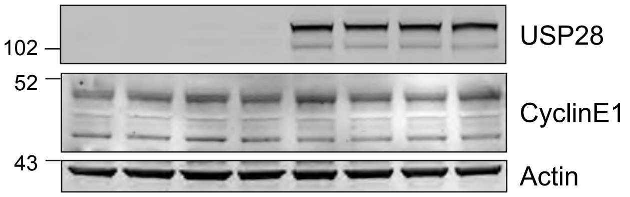


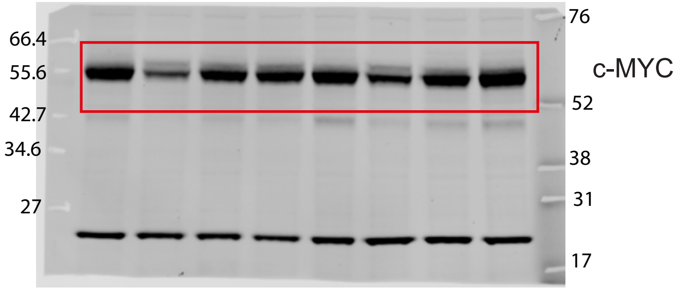


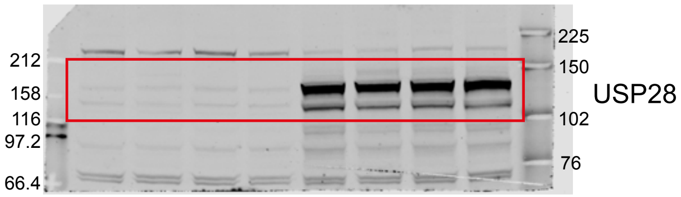


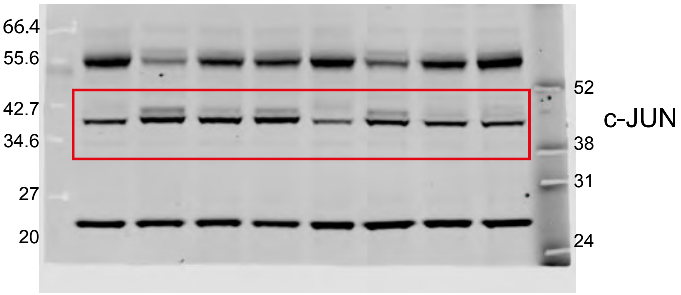


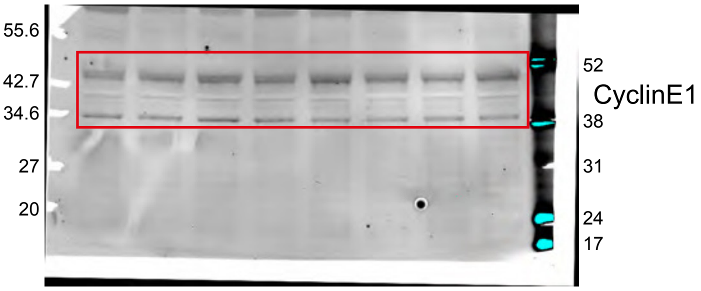


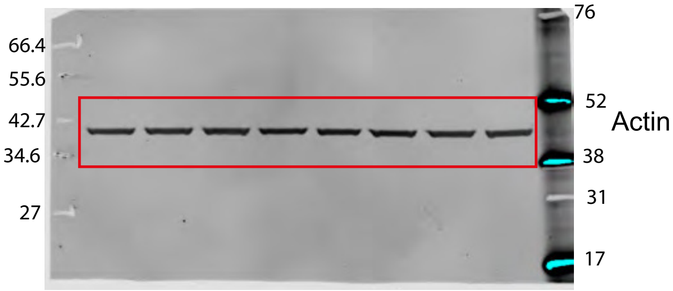


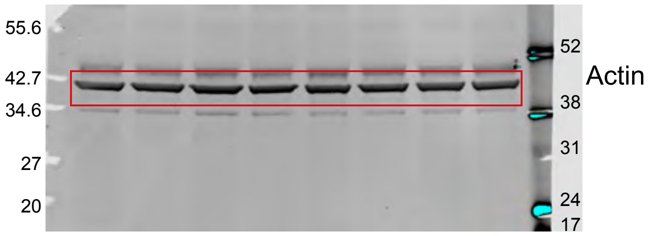

Uncropped blots for Figure S6c,d,f

### Figure S28

Figure S6c

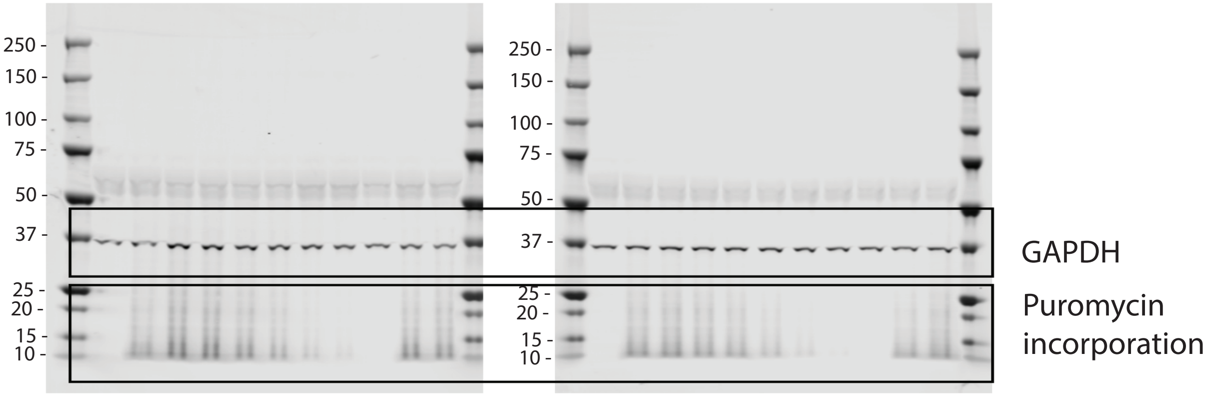


Figure S6d

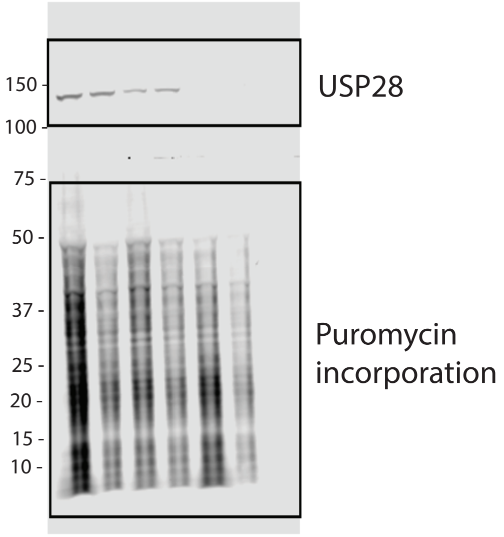


Figure S6f

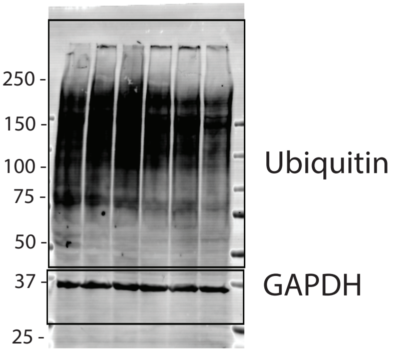

**Figure S29**

Uncropped blots for Figure S8b

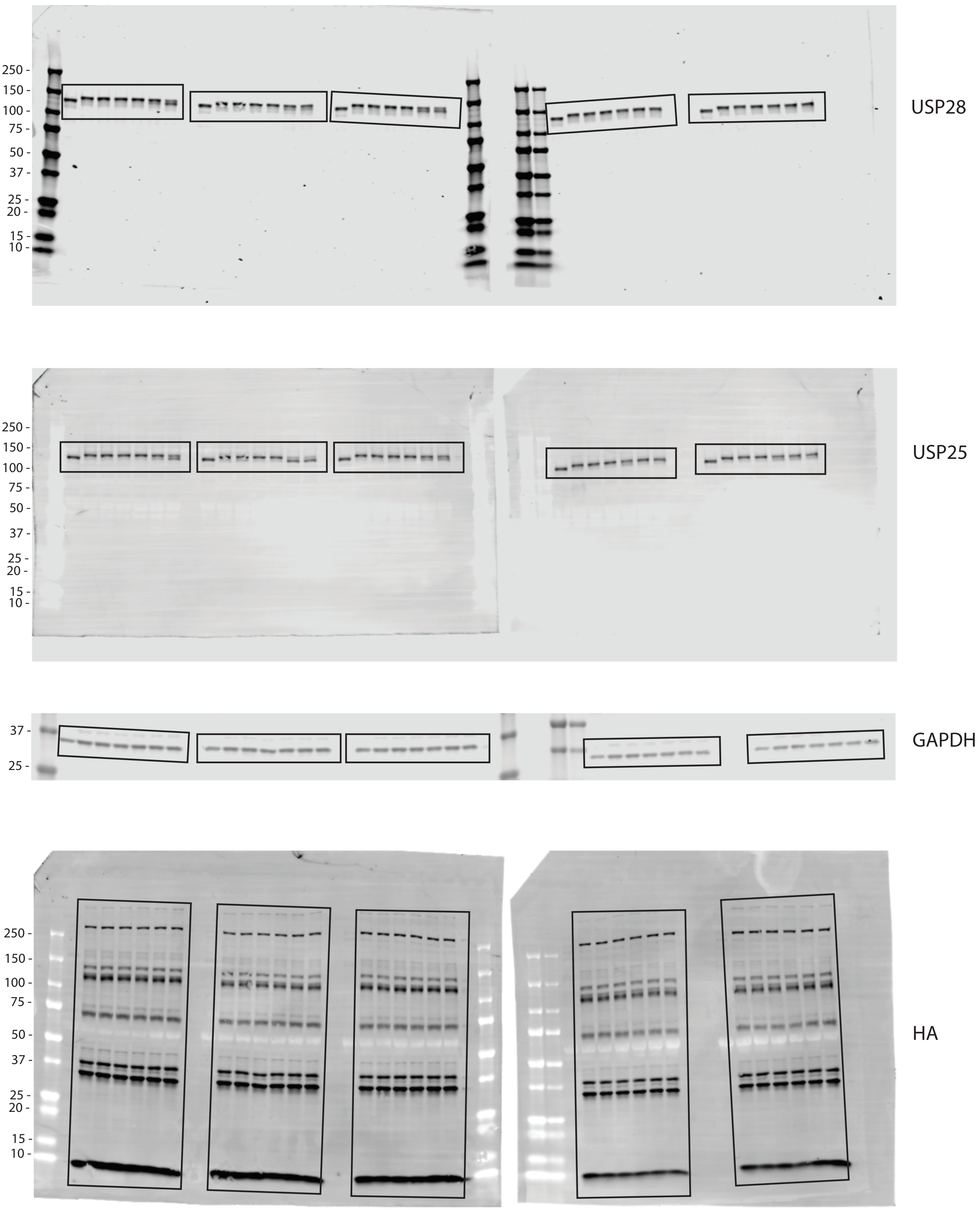

### Figure S30

Uncropped blots for Figure S14

Figure S15b

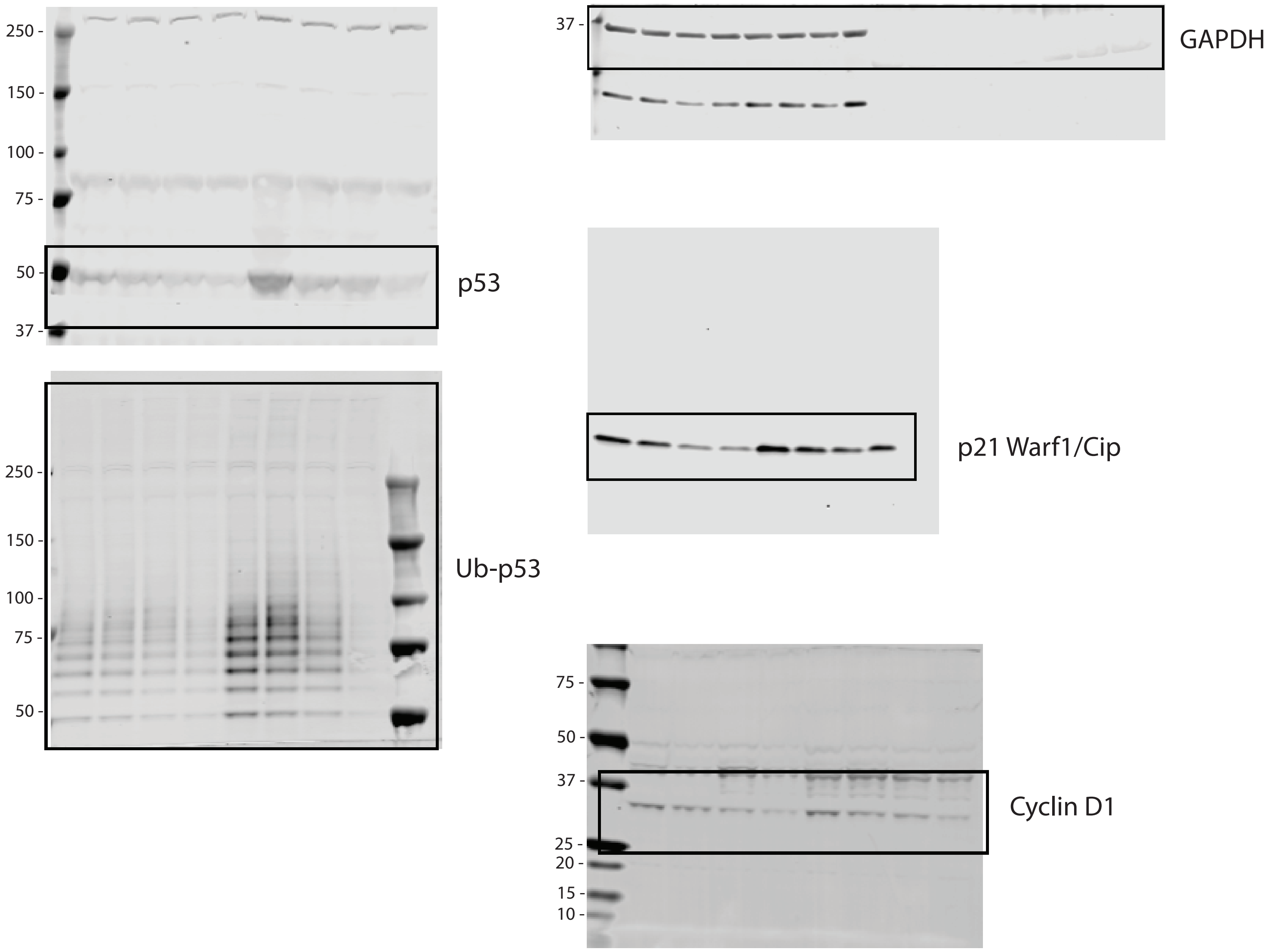


Figure S15c

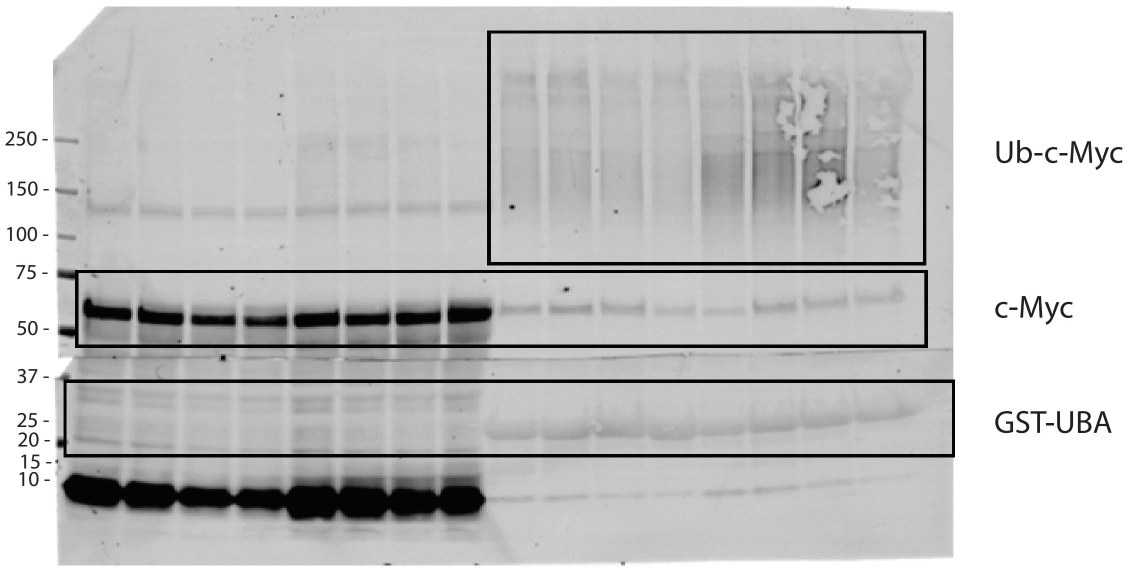

**Figure S31**

Uncropped blots for Figure S15d

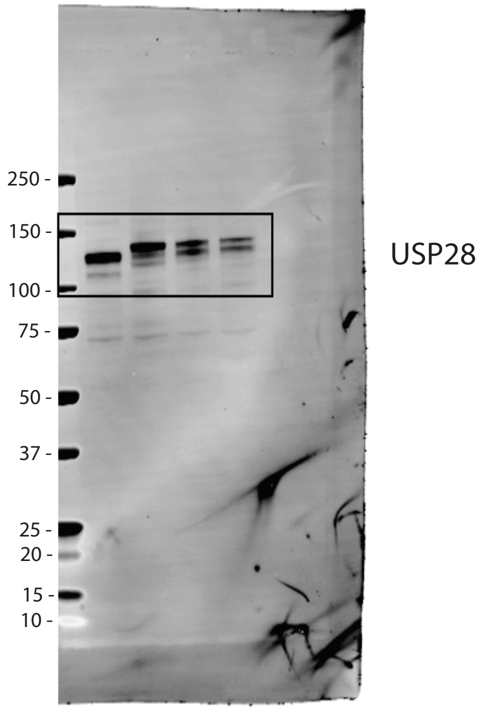


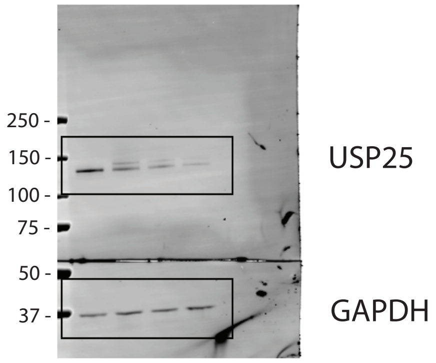

**Figure S32**

Uncropped blots for Figure S16a

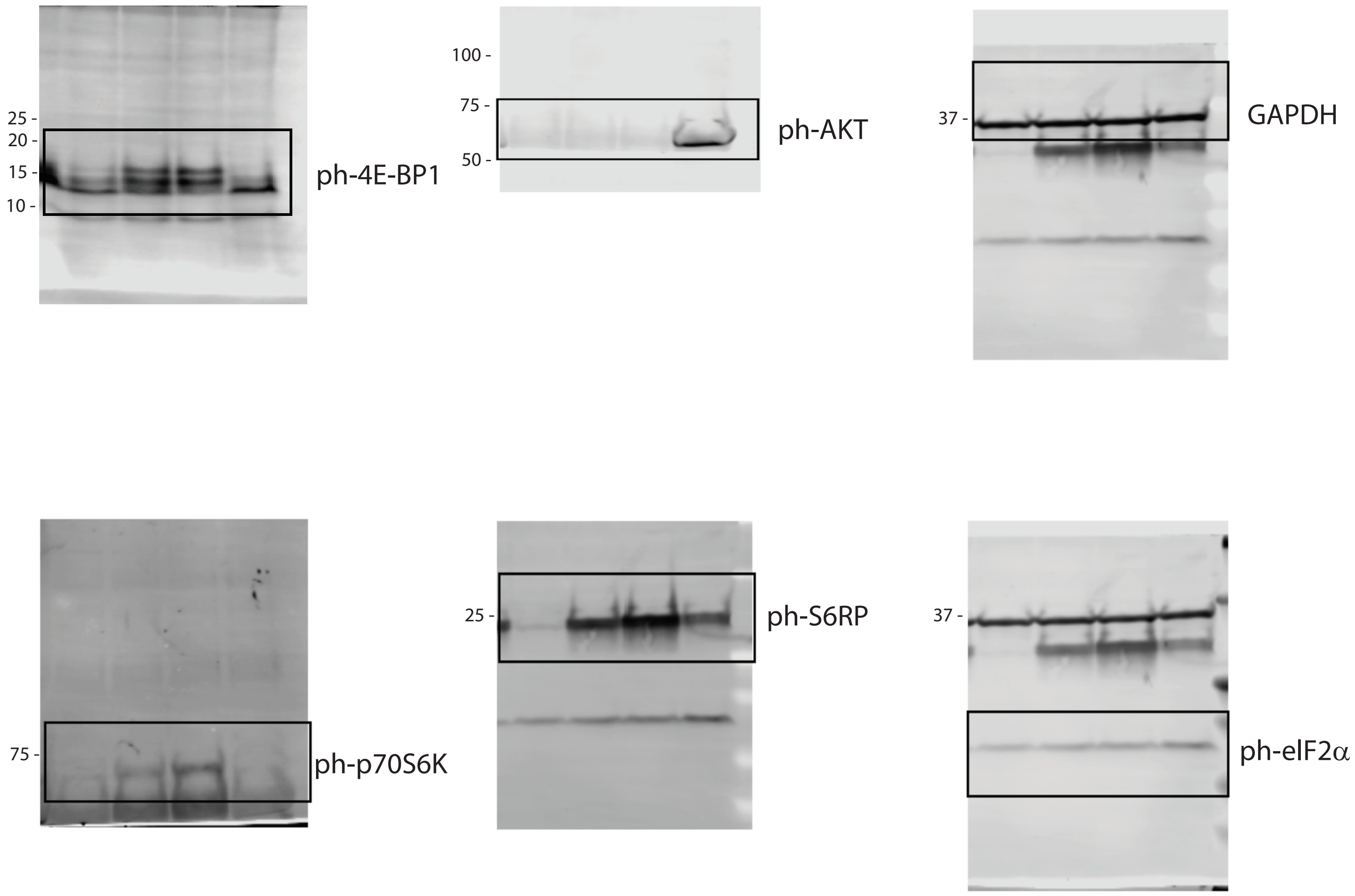
