## SUPPLEMENTAL INFORMATION for "Refined USP25/28 inhibitors with enhanced selectivity towards c-Myc driven squamous lung cancer cells"

**SUPPLEMENTARY INFORMATION**

1. **Material and Methods**

2. **Supplementary Figures**

3. **Supplementary Results & Extended Discussion**

4. **References**

### 1. Material and Methods

**Small molecule inhibitors used in this study (related to Figure S9)**

| Short Name | Original Name | Comments |
|---|---|---|
| FT206 | FT3951206 /CRT0511973 | Reference [1] |
| FT224 | FT3934224 / CRT0510305 | Forma Therapeutics WO2017139778 |
| FT592 | FT3880592 / CRT0422310 | (for USP28 X-ray structure) |
| FT639 | FT3973639-1 | (biotinylated FT224) WO2020033707A1 |
| FT811 | FT3947811 / CRT0511519 | Forma Therapeutics WO2017139779 |
| AZ1 (FT214) | FT3974214 | Reference [2] |
| bin68 | bin-01-68 | Sarah Buhrlage lab, references [3,4] |
| bin07 | bin-01-07-07 | Sarah Buhrlage lab, references [3,4] |
| bin74 | bin-01-74 | Sarah Buhrlage lab, references [3,4] |
| AV34 | AV-11324-34 | Sarah Buhrlage lab, references [3,4] |
| bin98 | bin-01-98 | Sarah Buhrlage lab, references [3,4] |
| bin41 | bin-01-41 | Sarah Buhrlage lab, references [3,4] |
| bin50 | bin-01-50 | Sarah Buhrlage lab, references [3,4] |
| bin55 | bin-01-55 | Sarah Buhrlage lab, references [3,4] |
| bin60 | bin-01-60 | Sarah Buhrlage lab, references [3,4] |
| bin70 | bin-01-70 | Sarah Buhrlage lab, references [3,4] |
| bin71 | bin-01-71 | Sarah Buhrlage lab, references [3,4] |
| bin73 | bin-01-73 | Sarah Buhrlage lab, references [3,4] |
| bin76A | bin-01-76A | Sarah Buhrlage lab, references [3,4] |
| bin78 | bin-01-78 | Sarah Buhrlage lab, references [3,4] |
| bin106 | bin-01-106 | Sarah Buhrlage lab, references [3,4] |
| bin111 | bin-01-111 | Sarah Buhrlage lab, references [3,4] |
| bin112 | bin-01-112 | Sarah Buhrlage lab, references [3,4] |

Chemical synthesis of small molecule inhibitors is described in patents as listed below (compounds with code name 'FT'). Synthesis of the compounds with code names 'bin' and 'AV' has been described previously [3–5]. For the compounds described in patents, we list the molecular formula, molecular weights and references to individual patents as follows:

**FT206:** Molecular Formula: $C_{25}H_{29}N_5OS$, MW: 447.601, Monoisotopic Mass: 447.209281 Da

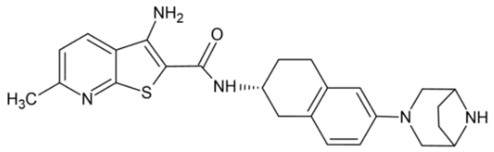


N-((2R)-6-(3,8-diazabicyclo[3.2.1]octan-3-yl)-1,2,3,4-tetrahydronaphthalen-2-yl)-3- amino-6-methylthieno[2,3-b]pyridine-2-carboxamide
WO2020033707 (see, e.g., Example 11-2)
https://patents.google.com/patent/WO2020033707A1/en

**FT224:** Molecular Formula: C20$H_{24}N_6OS$, MW: 396.513, Monoisotopic Mass: 396.173229 Da

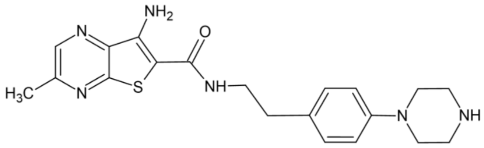


7-amino-3-methyl-N-(4-(piperazin-1-yl)phenethyl)thieno[2,3-b]pyrazine-6-carboxamide
WO2017139778 (see, e.g., Example 79-1)
https://patents.google.com/patent/WO2017139778A1/pt
https://patentimages.storage.googleapis.com/aa/d5/26/7fde2363a2bbe1/WO2017139778A1.pdf

**FT592:** Molecular Formula: $C_{23}H_{24}N_6OS$, MW: 432.546, Monoisotopic Mass: 432.173229 Da

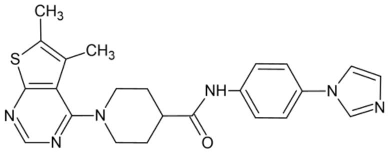


1-(5,6-dimethylthieno[2,3-d]pyrimidin-4-yl)-N-(4-imidazol-1-ylphenyl)piperidine-4-carboxamide
https://pubchem.ncbi.nlm.nih.gov/compound/31409966
Commercially available from Ambinter (https://www.ambinter.com/molecule/20903368)

**FT639** (biotinylated derivative of FT224): Molecular Formula: $C_{37}H_{51}N_9O_6S_2$, MW: 781.992, Monoisotopic Mass: 781.340 Da

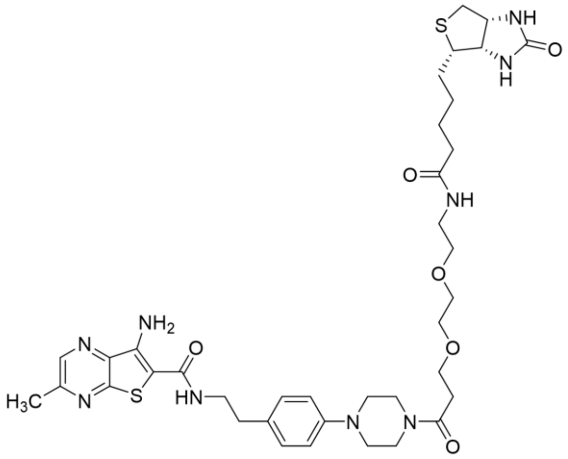

7-amino-3-methyl-N-(4-(piperazin-1-yl)phenethyl)thieno[2,3-b]pyrazine-6-carboxamide modified on the secondary amine of the piperazine moiety with Biotin-PEG2-N-hydroxysuccimid ester
(Conju-Probe: https://conju-probe.com/product/biotin-peg2-nhs/?gad_source=1&gclid=EAIaIQobChMIsrWrlrXYhwMV4plQBh2BTQD2EAAYAiAAEgKrfPD_BwE)
WO2020033707
https://patents.google.com/patent/WO2020033707A1/en

**FT811**: Molecular Formula: $C_{22}H_{24}F_2N_6OS$, MW: Formula Weight: 458.531, Monoisotopic Mass: 458.170036 Da

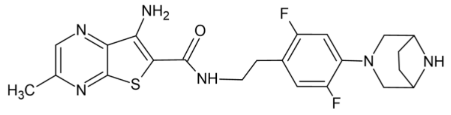

N-(4-(3,8-diazabicyclo[3.2.1]octan-3-yl)-2,5-difluorophenethyl)-7- amino-3-methylthieno[2,3-b]pyrazine-6-carboxamide
WO2017139779 (see, e.g., Example 17-1)
https://patentscope.wipo.int/search/en/detail.jsf?docId=WO2017139779

**Cell lines, antibodies and reagents**

The human LSCC (NCI-H520) line was provided by the Francis Crick Institute Cell Services and cultured in RPMI-1640 medium supplemented with 10% FBS, 1% penicillin/streptomycin, 2 mM glutamine, 1% NEEA, and 1 mM Na pyruvate as described previously [1]. HCT116 and MCF7 cell lines were cultured in high-glucose DMEM medium supplemented with 10% FBS, 1% penicillin/streptomycin and 2 mM glutamine. HAP1 cells were cultured in Iscove's Modified Dulbecco's Medium (IMDM) supplemented with 10% FBS, 1% penicillin/streptomycin and 2 mM glutamine as described [6]. FT224 compound-resistant (CR) HAP1 cells were generated as follows: HAP1 cells were cultured for three months with weekly stepwise increases in FT-224 concentration, starting at 1 μM and reaching 3 μM with a stable growth rate. HAP1 *USP28* knockout (KO) were generated by Horizon Inc. (Horizon Cat # C631) and generously provided by Dr Sebastian Nijman. MCF7 *USP28* knockout (KO) cells were generated and kindly provided by Dr Joss Chapman (Oxford, UK). HCT116 *USP28* knockout (KO) cells were generated using CRISPR-Cas9 with one of two independent USP28-specific sgRNAs targeting exon 5 and 7 respectively (**Figure S4**). sgRNAs were cloned into pSpCas9(BB)-2A-GFP (PX458; Addgene #4813846) for transfection into HCT116 Flp-In TREX cells (gift from Stephen Taylor, University of Manchester), followed by FACS and single cell dilution. HCT116 *FBW7* KO cells and matched parentals were a gift from the Vogelstein laboratory (John Hopkins, USA). For knockdown experiments shown in Figure 2d, MCF7 cells were transfected twice over 120 hours with USP28 siRNA #9 and #10 from Ambion Silencer Select Pre-designed siRNA (Thermo Fisher Scientific; 5'-GUGAUUGCUUUAUACCGA and 5'-GAUUAUAGUUUGUUCCGAA-3') at 40 nM final concentration using Oligofectamine (Thermo Fisher Scientific 12252011). For all other knockdown experiments, the following siRNA reagents were used: USP28 siRNA on-TARGET plus SMART plus (Dharmacon #L-006076-00) and USP25 siRNA on-TARGET plus SMART plus (Dharmacon #L-006074-00). For cell transfection experiments, the indicated siRNA sequences were mixed with Lipofectamine RNAimax (Invitrogen #13778-150) following the manufacturer's instructions as described previously [7]. All cells were tested Mycoplasma-negative and were maintained at 37°C with 5% CO2. Antibodies used in this study are listed in Table 2. Arginine/lysine free DMEM "light" medium, L-lysine (Lys0), L-arginine (Arg0), L-lysine-2H4 (Lys4), L-arginine-U-13C6 (Arg6), L-lysine-U-13C6-15N2 (Lys8) and L-arginine-U-13C6-15N4 (Arg10) were purchased from Silantes.

**Activity-based protein profiling (ABPP) assays**

Molecular active site probes based on the ubiquitin scaffold were generated and used essentially as described [8,9]. In brief, HA-tagged Ub propargyl probes were synthesized by expressing the fusion protein HA-Ub75-intein-chitin binding domain in Escherichia coli BL21 strains. Bacterial lysates were prepared, and the fusion protein was purified over a chitin binding column (NEB labs, UK). HA-Ub75-thioester was obtained by incubating the column material with mercaptosulfonate sodium salt (MESNa) overnight at 37°C. HA-Ub75-thioester was concentrated to a concentration of ~1 mg/mL using 3000 MW filters (Sartorius) and then desalted against PBS using a PD10 column (GE Healthcare). After diluting the HA-Ub-MESNa stock to 1.5 mg/mL, 250 mM propargylamine (or bromo-ethylamine) was added to HA-Ub-

MESNa (0.5 mL), at a pH range of 8–9 for 20 min at ambient temperature, to generate HA-Ub-PA (or HA-Ub-$C_2Br$). This was followed by a desalting step against phosphate buffer pH 8 as described above. Ub probe material was concentrated to ~1 mg/mL, using 3000 MW filters (Sartorius), and kept as aliquots at –80°C until use. Activity-based protein (ABPP) assays were performed as described previously described [8,9]. In brief, HAP1, HCT116 or MCF7 cell extracts were prepared using glass-bead lysis in 50 mM Tris pH 7.4, 5 mM $MgCl_2$, 0.5 mM EDTA, 250 mM sucrose, 1 mM DTT as described previously [6,10,11]. In brief, for experiments with crude cell extracts, ~50 µg of crude cell extract material was incubated with different concentrations of USP25/28 inhibitor compounds (FT224 – **Figure 1e**; FT811 and FT214/AZ1 – **Figure S1a, b;** FT224, AV-11324-34, bin-01-74, bin-01-68 and bin-0707 – **Figure S8b**) for 1 hr at 37°C, followed by the addition of 1 µg HA-UbPA and incubation for 30 min at 37°C. Samples were then subsequently boiled in reducing SDS-sample buffer, separated by SDS-PAGE and analysed by Western blotting using anti-HA (Roche, 1:2000), anti-USP28 (Abcam, 1:1000), anti-USP25 (Abcam, 1:1000), anti-GAPDH (Invitrogen, 1:1000), or beta Actin (Abcam, 1:2000) antibodies.

**OPP and SunSet assays to measure newly synthesized proteins**

The O-propargyl-puromycin (OP-Puro / OPP) labelling assay has been performed essentially as described previously [12,13]. In brief, HAP1 cells exposed to USP25/28 inhibitor for 4 hrs were incubated with 50 µM of OP-Puro in 10% fetal bovine serum (FBS) / Dulbecco's Modified Eagle Medium (DMEM) for 1 hour. Subsequently, cells were harvested and washed with ice-cold PBS. Cells were then fixed with 1% paraformaldehyde in PBS for 15 min on ice, washed in PBS, followed by permeabilization using permeabilization buffer (3% FBS and 0.1% saponin in PBS) for 5 min at room temperature. The Click chemistry reaction was performed by incubating cells with Click solution (1 mM CuSO4, 5 µM azide-fluor 488, 150 mM Tris, pH8.5 and 100 mM ascorbic acid adding before use) for 30 min at room temperature. The cells were washed three times in permeabilization buffer and resuspended in PBS for confocal microscopy imaging. The SunSet assay was performed essentially as described [14,15].

**Western blotting assays**

1-5x$10^6$ cells were lysed in ice-cold lysis buffer (20 mM Tris HCl, pH 7.5, 5 mM MgCl2, 50 mM NaF, 10 mM EDTA, 0.5 M NaCl, and 1% Triton X-100) or RIPA lysis buffer (10 mM Tris-HCl pH 7.5, 150 mM NaCl, 1% NP40, 0.1% SDS, 1% sodium deoxycholate) that was completed with protease, phosphatase, and kinase inhibitors (cOmplete Protease Inhibitor Cocktail tablets, Roche; protease inhibitor and phosphatase inhibitor cocktails I and II, Sigma ) following the manufacturer's instructions. Protein extracts were mixed with reducing SDS-Laemmli sample buffer (1x final concentration) and separated by SDS-PAGE, transferred to a nitrocellulose membrane, and blotted with antibodies using blotting conditions as described in **Table 2**. Primary antibodies were detected using secondary goat antibodies against mouse (IRDye 800CW) or rabbit (IRDye 680CW) IgGs (dilution: 1:20,000) and visualized with the Odyssey infrared imaging system (LI-COR, Biosciences). Uncropped Western blots for all main and supplementary figures are provided in **Figures S18-32**.

**FT639 Chemoproteomics and pulldown assays**

~500μg HAP1 cells were lysed using RIPA (50 mM Tris-HCl (pH 7.4), 150 mM NaCl, 1% Triton X-100, 0.5% sodium deoxycholate and 0.1% SDS) or glass bead lysis buffer (described above) supplemented with 0.2% NP-40, as indicated, and supplemented with phosphatase inhibitors. Lysates were homogenized by passing through a needle 10 times to ensure efficient cell disruption. Clarified lysates were collected by centrifugation. For chemical proteomics profiling, lysates were incubated with and without the FT639 at the indicated concentrations for three hours. In addition to beads-only controls, for some experiments, two control conditions were used: (i) DMSO-treated lysates with streptavidin beads alone, and (ii) competition assays in which lysates were pre-incubated with excess unbound compound for 30 minutes at room temperature before addition of FT639. ~50 μg Beads (Dynabeads™ MyOne™ Streptavidin C1) were washed four times with lysis buffer (glass beads lysis buffer + 0.2% NP-40 (GBN) for the mild conditions or RIPA for the stringent conditions). Incubation with inhibitor-treated or control lysates was performed under rotation for 2 hours at ambient temperature. After four washes with lysis buffer (glass beads lysis buffer + 0.2% NP-40 (GBN) for the mild conditions or RIPA for the stringent conditions), bound proteins were eluted twice with 55 µL of 3X SDS-Laemmli sample buffer (boiled for 5 minutes), separated by SDS-PAGE and analysed by immunoblotting. For proteomics analysis, the sample preparation was performed as previously reported [7]. Briefly, combined eluates were subjected to two rounds of chloroform-methanol precipitation to remove detergents and concentrate proteins. The resulting protein pellets were resuspended and digested in-solution with trypsin overnight at room temperature. Peptides were desalted and analysed by nano-liquid chromatography-tandem mass spectrometry (LC-MS/MS) for protein identification and quantification.

**Analysis by quantitative mass spectrometry**

*ABPP-MS assays* (**Figure 1a**). Mass spectrometry based analysis of HA-Ub active site probe pulldown experiments in presence of different concentrations of the inhibitor FT206 were performed essentially as described [8,9] with some modifications. In brief, anti-HA immunoprecipitated material from ~500 µg to 1 mg of MCF7 cell crude extract prepared as described for ABPP assays (see above) was subjected to in-solution trypsin digestion and desalted using C18 SepPak cartridges (Waters) based on the manufacturer's instructions. Digested samples were subjected to analysis by nano-UPLC-MS/MS using a Dionex Ultimate 3000 nano UPLC with EASY spray column (75 µm × 500 mm, 2 µm particle size, Thermo Scientific) with a 60 min gradient of 0.1% formic acid in 5% DMSO to 0.1% formic acid to 35% acetonitrile in 5% DMS at a flow rate of ~250 nL/min (~600 bar / 40 °C column temperature) coupled to an Orbitrap Q Exactive High Field (QE HF) mass spectrometer as described previously [16]. Protein identification and label-free quantitation were performed using MaxQuant Software (V1.5.3.8). Analysis of variance (ANOVA) (multiple comparison; original FRD method of Benjamini and Hochberg) was performed using GraphPad Prism software (version 7.0). Quantitative proteomics data were analysed using Perseus software (version 1.5).

*SILAC quantitation proteomics* (**Figure 2a, S2**). Stable isotope labelling of amino acids in cell culture (SILAC)-based quantitative proteomics was performed in HCT116 wildtype, *FBW7*$^{-/-}$ (KO) or *FBW7*$^{+/+}$ (wild-type) cells. Cells were cultured in arginine/lysine free DMEM "light" medium supplemented with L-lysine (Lys0) and L-arginine (Arg0) (FBW7 KO cells), "medium" with L-lysine-$^{2}H_4$ (Lys4) and L-arginine-U-$^{13}C_6$ (Arg6) (wildtype or FBW7 KO cells) or "heavy" with L-lysine-U-$^{13}C_6$-$^{15}N_2$ (Lys8) and L-arginine-U-$^{13}C_6$-$^{15}N_4$ (Arg10) (wildtype cells) for three weeks. Subsequently, wildtype or FBW7 KO cells were treated either with DMSO or 5 µM FT224 for 4 hrs according to the experimental outline shown in **Figure S2a**. Cells were collected, lysed in 50 mM Tris, pH 6.8, 2 % SDS and 10 % glycerol. Relative protein concentrations of the lysates were determined using a bicinchoninic acid assay (Thermo Fisher Scientific). Light-, medium-and heavy labeled lysates were combined in a 1:1:1 ratio composed of 200 µg of each of the following – A:B:D and A:C:D:

| SAMPLE | GENOTYPE | LABEL | TREATMENT |
|---|---|---|---|
| A | FBW7 $^{-/-}$ | Light | DMSO 0.05% final, 4hrs |
| B | FBW7 $^{-/-}$ | Medium | FT224 5 µM 4hrs |
| C | FBW7 $^{+/+}$ | Medium | DMSO 0.05% final, 4 hrs |
| D | FBW7 $^{+/+}$ | Heavy | FT224 5 µM 4hrs |

Samples were kept at -80°C until analysis. A fractionation approach to enhance the cellular (deep) proteome coverage was performed by off-line high-pH reverse-phase HPLC prefractionation of digested material to 50 fraction that were concatenated down to 10 fractions as reported previously [17]. Mass spectrometry analysis was performed using nano-LC-MS/MS on U3000 nano UPLC coupled to an orbitrap Q Exactive HF mass spectrometer (Thermo Fisher) as reported previously [16]. All raw MS files from the biological replicates of the SILAC proteome experiments were processed using MaxQuant software (version 1.5.3.8) as reported [16]. In brief, MS/MS data was searched against the Uniprot database (uniprotHumanUP000005640.fasta, retrieved July 2015) and the default settings [18]. The minimum required peptide length was set to six amino acids, and two missed cleavages were allowed. Cysteine carbamidomethylation was set as a fixed modification, whereas oxidation and N-terminal acetylation were considered as variable modifications. ProteinGroup text files were further processed using Excel, and the $\log_2$ transformed normalized ratios were plotted using JMP software (version 13.0.0) (**Figure S2b**).

*Chemoproteomics with FT639* (**Figure 3b**). Peptide material was subjected to LC-MS/MS analysis as described previously [9]

*TMT proteomics in HAP1 cells adapted to FT224* (**Figure S13a,b**). Proteomic analysis of HAP1 wild-type (WT) and USP28 knockout (KO) cells treated with FT224 under acute and chronic conditions was performed using tandem mass tag (TMT) labelling (Thermo Fisher Scientific, Cat. No. 901110) according to the manufacturer's instructions. Peptides were fractionated by off-line high-pH reverse-phase HPLC (50 fractions concatenated into 10) [17] and analysed on a Dionex Ultimate 3000 nano UPLC with EASY spray column (75 µm × 500 mm, 2 µm particle size, Thermo Scientific) coupled to a Q Exactive HF mass spectrometer as reported [16].

*Proteomics and ubiquitomics analysis of HAP1 cells exposed to FT224* (**Figure S13c**). Proteomics, ubiquitomics and label-free quantitative analysis (LFQ) were performed essentially as described previously [7].

*Proteomics analysis of short-lived proteins after treatment with FT224 or FT709* (**Figure S14**). Label-free quantitative proteomics with FT709 was performed essentially as described previously [17].

The mass spectrometry proteomics data have been deposited to the ProteomeXchange Consortium via the PRIDE [19] partner repository with the dataset identifier PXD075950.

### CRISPR screen

The Human CRISPR Activation Library (SAM v1) was a gift from Feng Zhang (Addgene # 1000000057). Lentivirus was produced for the sgRNA library, the Cas9-VP64 fusion plasmid (Addgene # 61425) and the MS2-P65-HSF1 plasmid (Addgene # 89308) by co-transfection with second-generation lentiviral packaging vectors pMD2.G (Addgene, # 12259) and psPAX2 (Addgene, # 12260) into HEK293T cells. Virus titre was determined for optimizing transduction at MOI<0.3 in HAP1 cells in T175 tissue culture flask format. For the screen, HAP1 cells were transduced first with lentivirus containing dCas9-VP64 complex in combination with lentivirus containing transcriptional activators p65 and HS1 that make up the SAM system, positive cells were selected using blasticidin, hygromycin, and zeocin resistance. Cells were then transduced at MOI < 0.3, so that a single cell carried one sgRNA, with lentiviruses containing SAM library sgRNAs. Two biological replicates with at least 150 × 106 HAP1 cells were infected in 12-well plates at 2000 rpm for 2 hrs at 37°C, to achieve a 500-fold coverage of the library. After zeocin selection, cells were treated with FT224 at $IC_{50}$ and $IC_{99}$ concentrations (1μM or 3μM, respectively) or DMSO control up to ~150 hours. Genomic DNA extraction was performed using the QIAamp Blood Maxi kit (# 51194) according to the manufacturer's protocol. The virally integrated CRISPR guide sequences were amplified by PCR and sgRNA sequences were identified by next generation sequencing (NGS) using the Illumina HiSeq 4000 platform (Oxford Genomics Centre, Wellcome Trust Centre for Human Genetics). Gene rankings were generated using the MAGeCK algorithm [20,21]. CRISPR data will be uploaded to the Gene Expression Omnibus (GEO) – NCBI – NIH database.

### Protein expression and purification for crystallography

A plasmid containing insert-deleted apo form human USP28 catalytic domain (residues 108-399[GSGSGS]580-738) was transformed into BL21(DE3) *E.coli* cells using standard methods. Ampicillin was used in cultures at 100 μg $mL^{-1}$. Growth was carried out in 2 L baffled flasks at 37C in LB media with the culture being cooled to 18 °C and, at an $OD_{600}$ of ~0.6 A, induced with 0.25 mM IPTG. After overnight growth, the cells were spun out at 3300 xg and frozen at -20C. The pellet was taken up in lysis buffer (50 mM HEPES pH 8, 300 mM NaCl, 10 % Glycerol, 5 mM Imidazole, 1 mM TCEP, 1 mM AEBSF, DNase) and soluble protein released by cell disruption at 30KPSI. Cell debris were removed by a 49,000 xg spin over 60 mins at 4 °C. The clarified supernatant containing USP28 protein was loaded onto a 5 mL HisTrap Excel column and washed using wash buffer (50 mM HEPES, pH 8, 300 mM NaCl, 10% Glycerol, 10mM Imidazole, 1mM TCEP). Elution of His-tagged USP28 protein was achieved using a step to 250 mM Imidazole. The pooled sample from NiIMAC elution was diluted 1 in 10 using buffer A (20 mM HEPES pH 8, 10 % glycerol, 1 mM TCEP) and loaded onto an anion exchange 6 mL Resource Q column. After washing, the protein was then eluted over a 20 cv gradient to 1 M NaCl. The resulting dominant USP28 peak was pooled and concentrated. The USP28 sample was subsequently purified by size-exclusion chromatography using a Superdex S75 column equilibrated in buffer (50 mM HEPES pH 8, 100 mM NaCl, 10 % glycerol, 2 mM TCEP) and the main USP28 containing peak collected and concentrated to 12mg $mL^{-1}$.

**X-ray crystallography of USP28-FT592 inhibitor complex**

A crystal of insert-deleted apo form human USP28 catalytic domain was grown in PACT condition E3 (20 % w/v PEG 3350, 0.2 M sodium iodide) at 20C, soaked in 5mM FT592 overnight at 20C, and flash-frozen in liquid nitrogen. Data were collected at 100K on a Rigaku Micromax-007 HF X-ray generator with Varimax HF Optics and a Rigaku Saturn 944 CCD detector. The crystal belonged to space group H32 with unit cell dimensions a=b= 107.8, c= 274.7 Å, and 1 molecule in the asymmetric unit, and data were processed to 3.4 Å resolution using StructureStudio (Rigaku). The structure of USP28 in complex with FT592 was solved by molecular replacement using PHASER [22] and human USP7 (PDB code= 1NBF) as the search model. The model was rebuilt using COOT [23] and refined using REFMAC5 [24]. Data collection and refinement statistics are shown in Table 1. The coordinates and diffraction amplitudes for the USP28-FT592 complex are deposited in the Protein Data Bank (PDB code ID 29GK).

**Cell growth, morphology and proliferation assays**

5,000 cells were seeded in 96-well plates, and cells were imaged in an IncuCyte Zoom Imager (Essen Bioscience) for live-cell imaging during the indicated times, at the indicated intervals, in the phase contrast channel. Cell growth was determined by measuring the percent confluence over time (as in **Figure 6, 7**, and **S15**). End point cell viability was measured using the Resazurin assay (Alamar Blue; Thermo Fisher #DAL1025), following the manufacturer's instructions. (as in **Figure 2e**).

**Statistical analysis**

Data are represented as mean ± SEM. Statistical significance was calculated with the unpaired two-tailed Student's t test, one-way or two-way ANOVA followed by multiple comparison test using GraphPad Prism software. A p value that was less than 0.05 was considered to be statistically significant for all data sets. Significant differences between experimental groups were: *$p < 0.05$, **$p < 0.01$, or ***$p < 0.001$ or ****$p < 0.0001$. Biological replicates represent experiments performed on samples from separate biological preparations; technical replicates represent samples from the same biological preparation run in parallel.

**Microhelix acylation assay**

Assays were performed as described in Goto et al. [25] with minor alterations. Briefly, 6 µL solutions of mihx (41.67 µM), eFx (41.67 µM) in HEPES-KOH (pH 7.5, 83.3 mM) were heated at 95 °C for 2 minutes, cooled at room temperature for 5 minutes and $MgCl_2$ (3 M, 2 µL) added. After a further 5 minutes at room temperature the solution was cooled on ice for 5 minutes before addition of ClAc-L-Tyr-OCM [26] (50 mM in DMSO, 1 µL) and either 1 µL DMSO ("-" sample) or FT224 ("+" sample; 10 mM in DMSO, 1 µL). After thorough mixing, samples were incubated on ice for 45 minutes. Final concentrations of components in the reaction were as follows: HEPES-KOH (pH 7.5, 50 mm), eFx (25 µM), mihx (25 µM), $MgCl_2$ (600 mM), ClAc-L-Tyr-OCM (5 mM), FT224 (1 µM, where present). Following incubation, RNA was precipitated and analysed by 20% denaturing urea-acid-PAGE as described in Goto et al., [25].

### 2. Supplementary Figures

**Figure S1: USP28 inhibitors with various chemical scaffolds differentially affect USP28 versus USP25**

**a** Chemical structures of USP25/28 inhibitors FT811 and AZ1.

**b** FT811 (left panel) and AZ1 (right panel) potencies towards USP28 and USP25 assessed by the ABPP assay. 50μg MCF-7 cell extracts were exposed to FT224 inhibitor at the indicated concentrations for 30 min at 37°C, followed by adding HA-Ub-PA probe for 30 min, separation by SDS-PAGE and analysis by western blotting.

**Figure S2: FBW7-dependent and independent effects of the USP25/28 inhibitor FT224 on the HCT116 cell proteome**

**a** Stable isotope labeling of amino acid in cell culture (SILAC) based proteomic experimental setup using colorectal cancer HCT116 wildtype ($FBW7^{+/+}$) or FBW7 KO ($FBW7^{-/-}$) exposed to either 5 μM FT224 or DMSO for 4 hrs. Cells were harvested and lysed in SDS-Lysis buffer (50 mM Tris pH 6.8, 2% SDS, 10% glycerol). Lysates from light, medium and heavy labelled samples, were combined at a 1:1:1 ratio to obtain two triplexed samples that were processed for mass spectrometry.

**b** Scatter plot analysis comparing HCT117 FBW7 (Y-axis) versus KO (X-axis) treated with 5 μM FT224 or DMSO for 4 hrs. Differentially abundant proteins are indicated in orange, green, red, blue, teak, olive-green and purple, respectively. Proteins that are de-enriched in FT224-treated cells independently of FBW7 status include CCND1 and CCNB1 (both shown in bold) and are found in the left hand bottom quadrant (teal). Data shown are from one of two independent experiments.

**Figure S3: USP25/28 inhibitor effects on substrates of the FBW7-USP28 axis**

**a** Treatment of FBW7 wildtype and knockout HCT116 cells for 4 hrs with FT811 decreases USP28 substrate levels, independently of FBW7 status.

**b** Neither FT224, nor FT811 treatment of MCF-7 cells affects Cyclin D1 (CCND1) mRNA levels. Shown is the mean of two individual experiments except for the last datapoint.

**c, d** c-Myc and Cyclin D1 protein levels are decreased after 4 hrs treatment with FT811 or FT224 treatment and rescued by cotreatment with epoxomicin (Epo, 50 nM) or MLN4924 (MLN, 1 μM) for 4 hrs in HCT116 (**c**) and MCF-7 (**d**) cells.

**Figure S4: Generation of HCT116 USP28 KO cells**

**a** CRISPR strategy to generate USP29 KO clones using two distinct (#98 and #99) sgRNAs targeting exons 5 and 7 within the USP catalytic domain. Western blot shows successful loss of USP28 protein expression. Shown are two edited alleles for one sgRNA 98 targeted clone (98-19) and a single sequence for one sgRNA 99 targeted clone (99-2).

**b** Representative western blot showing the loss of USP28 protein expression in a panel of USP28KO clones is associated with a decrease in Cyclin D1 levels.

**c - e** Scatter plots showing a correlation between USP28 and Cyclin D1 protein abundance in clonal cells targeted with either sgRNA #98 or #99. Each graph shows a quantitation of a distinct western blot.

**Figure S5: Characterisation of HCT116 USP28 KO cells**

**a** Western blot showing a panel of cell cycle proteins and reported USP28 substrates in USP28 KO (98-3) and control (98-17) HCT116 clonal cells treated with FT224 or FT811 for 4 hrs.

**b** Western blot showing a panel of cell cycle proteins and reported USP28 substrates in HCT116 USP28 KO 99-2 and control (99-9) clonal cells treated with FT224 or FT811 for 4 hrs.

**Figure S6: USP28 inhibitor off-target in the translation machinery**

**a** Reticulocyte lysate luciferase assay for assessing inhibition of protein synthesis as described in the methods section.

**b** FT224, but not FT214 (AZ1) inhibitor, interferes with protein synthesis as measured by the OPP assay.

**c** FT224 affects protein synthesis in wildtype HAP1 and USP28 KO cells as measured by puromycin incorporation, SDS-PAGE and immunoblotting.

**d** FT224 affects protein synthesis in wildtype HAP1, compound resistant (CR – **Figure S15**) and USP28 KO cells.

**e** Quantitation of (**d**).

**f** FT224 treatment depletes conjugated ubiquitin levels in HAP1 cells as assessed by SDS-PAGE and anti-ubiquitin western blotting.

**Figure S7: USP28-inhibitor structure characterization**

**a** Chemical structure of FT592 (N-(4-(1H-imidazol-1-yl)phenyl)-1-(5,6-dimethylthieno[2,3-d]pyrimidin-4-yl)piperidine-4-carboxamide). Figure prepared using ChemDraw Prime 23.0.1.10.

**b** Superposition of human USP28 in complex with FT592 in grey on apo form human USP28 in purple (PDB code 6HEH). FT592 is shown as a stick representation with carbon atoms colored green. The thumb, palm and fingers subdomains of the catalytic domain, catalytic cysteine (C171), and α5 are highlighted.

**c** View in panel **b** rotated on the y-axis by 50°.

**d** Left: superposition of human USP28 in complex with FT592 in grey (FT592 carbon atoms in green) on human USP28 in complex with FT206 in gold (FT206 carbon atoms in yellow; PDB code 8P1Q); right: close-up view of the inhibitor binding pocket highlighting α5 and the catalytic cysteine, C171.

**e** Left: superposition of human USP28 in complex with FT592 in grey (FT592 carbon atoms in green) on human USP28 in complex with Vismodegib in brown (Vismodegib carbon atoms in orange; PDB code 8P14); right: close-up view of the inhibitor binding pocket highlighting α5 and the catalytic cysteine, C171.

**f** Left: superposition of human USP28 in complex with FT592 in grey (FT592 carbon atoms in green) on human USP28 in complex with AZ1 in salmon (AZ1 carbon atoms in purple; PDB code 8P1P); right: close-up view of the inhibitor binding pocket highlighting α5 and the catalytic cysteine, C171.

**g** Left: superposition of human USP28 in complex with FT592 in grey (FT592 carbon atoms in green) on human USP28 in complex with an unpublished inhibitor in lime (unpublished inhibitor carbon atoms in blue; PDB code 7TUO); right: close-up view of the inhibitor binding pocket highlighting α5 and the catalytic cysteine, C171.

**h** Superposition of human USP28 in complex with FT592 in grey (FT592 carbon atoms in green) on human USP7 in complex with GNE6776 in purple (GNE6776 carbon atoms in light brown; PDB code 5UQX) highlighting α5 and the catalytic cysteine, C171, in USP28.

**i** Superposition of human USP28 in complex with FT592 in grey (FT592 carbon atoms in green) on human USP7 in complex with GNE6640 in cyan (GNE6640 carbon atoms in violet; PDB code 5UQV) highlighting α5 and the catalytic cysteine, C171, in USP28.

Figure prepared using PyMOL (The PyMOL Molecular Graphics System, Version 2.5.8, Schrödinger, LLC.).

**Figure S8: USP25/28 inhibitors tested for protein translation activity and USP25/28 activity labelling**

**a** Upper panel – $IC_{50}$ values; lower panel – OPP values. Table and associated graph showing $IC_{50}$ values for translation inhibition (OPP assay) and cell viability for MCF-7 WT and USP28 KO cells treated with USP25/28 inhibitors.

**b** ABPP assay in HAP1 cell extracts. USP28 inhibitors FT224, AV-11324-34, bin-01-74, bin-01-68 and bin-07-07 were exposed to ~50μg cell extracts for 1 hr at room temperature, followed by HA-UbPA probe labelling for 45' at 37°C, separation by SDS-PAGE and analysis by immunoblotting.

**Figure S9: Chemical structures and Ub-Rhodamine assay based $IC_{50}$ values of refined USP25/28 inhibitors**

**Figure S10: Compound names and $IC_{50}$ values for the OPP assay**

Left column: compound name; right column: $IC_{50}$ values for protein synthesis as measured by the OPP assay. See also **Figure S8a**.

**Figure S11: USP28 inhibitor structure-activity relationship (SAR) analysis reveals chemical features associated with off-target translation inhibition**

Basis for the design of refined USP25/28 inhibitors where off-target effect is dialled out. Critical chemical moieties required for the interaction with specific USP28 amino acid residues are indicated in green and with arrows. Structural elements linked to off-target / protein synthesis inhibition are indicated in light purple (compounds listed in the left column) that were not present in compounds without noticeable OPP activity (compounds listed in the right column).

**Figure S12: CRISPR screen reveals USP28 inhibitor off-target in protein synthesis**

**a** Growth curves of HAP1 controls (Ctrl) and cells at $IC_{50}$ and $IC_{99}$ of FT224 inhibitor concentrations (1μM and 3μM, respectively). Examples of control and $IC_{50}$ growth characteristics of different cell clones are shown.

**b** Growth curves of HAP1 cells at Ctrl and $IC_{50}$ of FT224 inhibitor concentrations (3μM).

**c** $Log_{10}$ reads per sgRNA in HAP1 cells under control, $IC_{50}$ and $IC_{99}$ conditions. Two independent replicates (N=2) are shown.

**d** Volcano plots showing transcriptomes of HAP1 ctrl cells compared to $IC_{50}$ (left panel) and $IC_{99}$ (right panel) FT224 inhibitor concentrations (1μM and 3μM, respectively).

**Figure S13: USP28 inhibitor - omics reveals signatures of altered ribosomal quality control (RQC)**

**a** Volcano plot showing the USP28 KO proteome versus wildtype (WT) controls in HAP1cells.

**b** Volcano plot showing the USP28 inhibitor FT224 (3μM) adapted "compound resistant – CR" cell proteome versus DMSO controls (<1% final concentration) in HAP1 cells.

**c** Scatter plot showing the ubiquitome versus the proteome of USP28 inhibitor FT224 exposed for 24 hrs versus DMSO control (<1% final concentration) in HAP1 cells. The red frame indicates the area magnified out of the scatter plot. Ribosome proteins are indicated in blue, histones in green and metabolic enzymes in red, respectively.

**d** Reduced levels of proteins involved in translation after USP28 KD or inhibition. HAP1 wildtype and "compound resistant – CR" cells were treated with the indicated concentrations of FT224 for 6 hrs, followed by cell lysis, separation by SDS-PAGE and immunoblotting.

**e** mRNA levels of genes involved in the cell cycle & translation affected by acute or chronic USP28 inhibitor treatment, respectively. $Log_2$ transformed and Z-score normalised values are shown in a heatmap format (see also Supplementary Information).

**Figure S14: USP28 inhibitor effect on short-lived proteins**

**a** Quantitative proteomics analysis reveals short-lived proteins affected by 4h 3μM FT224 USP25/28 (3 μM), but not FT709 USP9X inhibitor (1μM) treatment in HCT116 cells originally described in [17]. The red frame indicates the area magnified out of the scatter plot. Protein half-lives were derived from published RPE1 data sets [27].

**b** FT224 treatment reduces Cyclin D1, c-Myc p53 and p21 protein levels and the ubiquitinated form of p53 as analysed by SDS-PAGE and immunoblotting.

**c** FT224 treatment reduces c-Myc protein levels and its ubiquitinated form as analysed by SDS-PAGE and immunoblotting.

**Figure S15: Characterisation of FT224 compound resistant cells**

**a** HAP1 cell growth (IncuCyte) characteristics in the presence of 1μM of USP25/8 inhibitor compound FT224. HAP1 CR: compound-resistant HAP1 cells.

**b** Microscopy pictures of HAP1 cells after 48, 72, 96 hrs and one-week inhibitor FT224 exposure to 1μM and 3μM and DMSO control.

**c** Microscopy pictures of HAP1 compound resistant "CR" cells after 48, 72, 96 hrs and one-week inhibitor exposure to FT224 at 1μM, 3μM and DMSO control (<1% final concentration).

**d** USP25/28 protein expression levels in HAP1 wildtype and CR cell exposed to 3mM FT224 for 3 hrs as well as controls. USP25/28 deubiquitinase activity was assessed by labelling cellular extracts with HA-$UbBr_2$ for 45 min at 37°C, followed by SDS-PAGE separation and immunoblotting.

**e** PCA plot of proteomics analysis comparing HAP1 cells exposed to DMSO (<1% final concentration – DMSO_0-3), 3μM FT224 exposure for 6 hrs (X3μM_6H0-3), 10μM FT224 exposure for 6 hrs (X10μM_6H0-3) or chronic exposure to 3μM FT224 (CR HAP1 cells – X3μM0-3). Experiments were performed in triplicates (N=3).

**f** Volcano plot of proteomics analysis comparing the proteomes of HAP1 control versus cells grown in the presence of FT224 for three weeks (compound resistant "CR" cells).

**Figure S16: Compound resistant cells upregulate protein translation**

Western blotting analysis of HAP1 cells exposed to different doses of FT224 or DTT at the indicated concentrations for 24 hrs, followed by cell lysis, separation by SDS-PAGE and immunoblotting. Antibodies used are indicated.

**Figure S17: USP28 inhibitor FT224 does not acylate tRNA**

Microhelix acylation assay showing a representative urea acid-PAGE gel image to visualise tRNA charging by molecular weight shift, performed as described previously [28,29]. The +/- lanes are with/without USP28 inhibitor FT224 at 1 μM final concentration, respectively. See Methods for additional details.

**Figures S18-32: Uncropped Western blots for all figures**

All gel band regions used to create figure composites are labelled with a red or black rectangle.

### 3. Supplementary Results & Extended Discussion

#### Thienopyridine carboxamide - omics reveals signatures of altered ribosomal quality control (RQC)

Uncovering off-targets is challenging. Several approaches were used including chemoproteomics, genetic and proteomics screens and cellular adaptation experiments. First, to complement mapping off-target effects of FT224 by Chemoproteomics (**Figure 3**), a genome-wide CRISPR gain-of-function (GOF) screen was performed to identify transcripts/proteins that could overcome compound-mediated toxicity [30]. We used delta CAS9 (dCAS9) –VP64 and MS2-p65 lentivirus constructs together with a lentiviral sgRNA expression library to infect HAP1 cells. These HAP1 cells were treated either with USP28 inhibitor FT224 at $IC_{50}$ (1 μM), $IC_{99}$ (3μM – not shown) or DMSO control and subjected to multiple rounds of growth cycles for 10 days (**Figure S12a**), where the growth rate of cells in $IC_{50}$ conditions became comparable to control cells after ~100 hours (**Figure S12b**). mRNA was subsequently extracted from two independent biological replicates and analysed by RNA Seq (**Figure S12c**). Interestingly, main sgRNAs enriched in $IC_{50}$ and $IC_{99}$ samples were component of the mitochondrial translation machinery, highlighting mitochondrial protein translation as a potential mechanism of resistance against the cytotoxic effect of FT224 (**Figure S12d**).

Second, we performed an integrative – omics approach. We compared the proteomes of wildtype HAP1 cells to USP28 KO and compound treated cells (HAP1 - **Figure S13**). A prominent effect was noted in HAP1 USP28-/- KO cells (**Figure S13a**). The latter profile correlated with the one observed in HAP1 cells treated for >3 weeks with FT224 at increasing concentrations, leading to growth in the presence of 3 μM of compound, referred to as "compound-resistant CR" cells (**Figure S13b, S15**).

Third, a cross-comparison of the compound-dependent proteomes and ubiquitomes (enriched ubiquitylated cellular proteins) was performed in HAP1 cells. A clear accumulation of ubiquitylated histones, ribosomal subunits and metabolic enzymes was observed, shown here after 24h FT224 treatment of HAP1 cells (**Figure S13c**). Immunoblotting confirmed the ubiquitylation of the ribosomal subunit RPS10, differential phosphorylation of elongation factor 2 (pThr56) and reduced protein levels of PRAME, a melanoma antigen preferentially expressed in tumors (**Figure S13d**). Prolonged compound exposure also affected cellular transcription profiles, in particular for c-Myc, LSD1, USP25 and a number of Cyclins, including D1, D2 and E1, but not E2 (**Figure S13e** This indicated compound-dependent effects on ribosome function/translation, similar to ribosomal stalling [31] that seem to be aggravated by prolonged incubation of HAP1 cells with FT224 (CR). Further evidence for a potential effect on translation was revealed by acute treatment of HCT116 cells with FT224, altering proteins with a short half-life that was not observed with a different inhibitor targeting USP9X (FT709) (**Figure S14a**) [16]. Proteins with short half-lives that were modulated by acute FT224 incubation in HCT116 cells included USP28 candidate substrates, such as c-Myc, p53, p21 and Cyclin-D1, all of which may have been affected at both, their translation and deubiquitylation level (**Figure S14b**). We observed similar trends for ubiquitylated p53 and c-Myc enriched by GST-UBA pulldowns, even in the presence of proteasome inhibitor Bz (**Figure S14b, c**), indicating that FT224 may act upstream of the expected deubiquitylation step. Also, this explained, at least in part, the delayed growth of cells acutely exposed to FT224 and reaching an adapted phenotype upon prolonged compound treatment up to 3μM (**Figure S13e**), a trait not observed upon genetic deletion of USP28 (**Figure 2, S4**). Taken together, USP28 compound FT224 dependent proteome, ubiquitome and RNAseq bulk transcriptome analyses suggested target-independent effects, possibly on the translation machinery.

Fourth, we characterized CML-derived HAP1 cells progressively adapted to grow at otherwise toxic concentrations of FT224 for a period of three months (**Figure S13, S15**). Compound resistant (CR) cells grew significantly slower than the parental cells (**Figure S15a-c**). ABPP assays showed that compound was still able inhibit USP25/28 in a similar extent as in the parental cells after acute treatment (**Figure S15d**). A distinct transcriptome profile was observed between parental (untreated and acutely treated) and CR cells (**Figure S15e**). This included transcriptional repression of cell cycle genes **(S15f**). Also, chronic FT224 compound exposure led to upregulated mRNA levels encoding for p53, p21, HIF1A, the multidrug resistance pump ABCB1, and other genes involved in the unfolded protein response (UPR) and protein transport such as HSPA8, XBP1 and DDIT3 (**Figure S13, S16**). Remarkably, an increase in S6RP phosphorylation was observed, suggesting induced mTORC1 activity (**Figure S16**). RRBP1 acts as a receptor for the ribosome in the ER membrane [32]. RPS27L differs from RPS27 by three amino acids. RPS27 is located in the 40S subunit of the ribosome and interactions with eIF3 occur near the mRNA exit channel [33]. RPS27A extends into the A site of the decoding centre on the small ribosomal subunit and would prevent tRNA binding [34]. EEF2 is required for the translocation of the tRNA2-mRNA module, in the interface between the small and large ribosomal subunits [35]. DHX29 is a RNA helicase and has key functions in translation initiation and, interestingly, like RPS27, also interacts with eIF3 [36]. Together, these experimental evidences support a possible off-target for thienopyridine / thienopyrazine carboxamide based compounds in inhibition of protein translation, correlating with affecting cell viability (**Figure 7d**).

As a major hallmark of altering translation and ribosome function, ubiquitylation of ribosomal subunits RS2, RS3, RPS10, RPS20 and others has been linked to ribosomal stalling under stress conditions [31,37–39]. We noted significantly ubiquitylated peptides for these proteins after compound treatment (**Figure S13c**). Interestingly, ubiquitylation of RPS10 seems to be more prevalent after longer incubation times (4 hrs) by immunoblotting (**Figure 4f, S13d**). Activation of the mTOR pathway upon inhibition of protein synthesis has also been reported [40,41]. Indeed, we could observe a dose-dependent activation by phosphorylation of S6RP, p70S6K and 4E-BP1, in particular upon prolonged inhibitor exposure, leading to a “compound resistant CR” phenotype (**Figure S15, S16**). Moreover, we could detect phosphorylation of S6RP (S235/S236) after just 10 minutes of incubation with FT224 (**Figure S15/S16**). To explore the exact mechanism of protein translation inhibition, we tested whether FT224 may interfere with tRNA amino acid loading in a tRNA acylation assay using microhelix RNA (mihx) and flexizyme (eFx) [25], but with no obvious effect (**Figure S17**).

### 4. References


1. Konermann, S., Brigham, M.D., Trevino, A.E., Joung, J., Abudayyeh, O.O., Barcena, C., Hsu, P.D., Habib, N., Gootenberg, J.S., Nishimasu, H., et al. (2015). Genome-scale transcriptional activation by an engineered CRISPR-Cas9 complex. Nature *517*, 583–588. https://doi.org/10.1038/nature14136.

2. Sundaramoorthy, E., Leonard, M., Mak, R., Liao, J., Fulzele, A., and Bennett, E.J. (2017). ZNF598 and RACK1 Regulate Mammalian Ribosome-Associated Quality Control Function by Mediating Regulatory 40S Ribosomal Ubiquitylation. Mol. Cell *65*, 751-760.e4. https://doi.org/10.1016/j.molcel.2016.12.026.

3. Clancy, A., Heride, C., Pinto-Fernández, A., Elcocks, H., Kallinos, A., Kayser-Bricker, K.J., Wang, W., Smith, V., Davis, S., Fessler, S., et al. (2021). The deubiquitylase USP9X controls ribosomal stalling. Journal of Cell Biology *220*. https://doi.org/10.1083/JCB.202004211.

4. Cui, X.A., Zhang, H., and Palazzo, A.F. (2012). p180 Promotes the Ribosome-Independent Localization of a Subset of mRNA to the Endoplasmic Reticulum. PLoS Biol. *10*, e1001336–e1001336.

5. Valášek, L.S., Zeman, J., Wagner, S., Beznosková, P., Pavlíková, Z., Mohammad, M.P., Hronová, V., Herrmannová, A., Hashem, Y., and Gunišová, S. (2017). Embraced by eIF3: structural and functional insights into the roles of eIF3 across the translation cycle. Nucleic Acids Res. *45*, 10948–10968. https://doi.org/10.1093/nar/gkx805.

6. Klinge, S., Voigts-Hoffmann, F., Leibundgut, M., Arpagaus, S., and Ban, N. (2011). Crystal Structure of the Eukaryotic 60<em>S</em> Ribosomal Subunit in Complex with Initiation Factor 6. Science (1979). *334*, 941 LP – 948. https://doi.org/10.1126/science.1211204.

7. Flis, J., Holm, M., Rundlet, E.J., Loerke, J., Hilal, T., Dabrowski, M., Bürger, J., Mielke, T., Blanchard, S.C., Spahn, C.M.T., et al. (2018). tRNA Translocation by the Eukaryotic 80S Ribosome and the Impact of GTP Hydrolysis. Cell Rep. *25*, 2676-2688.e7. https://doi.org/10.1016/j.celrep.2018.11.040.

8. Pisareva, V.P., and Pisarev, A. V (2016). DHX29 and eIF3 cooperate in ribosomal scanning on structured mRNAs during translation initiation. RNA *22*, 1859–1870. https://doi.org/10.1261/rna.057851.116.

9. Higgins, R., Gendron, J.M., Rising, L., Mak, R., Webb, K., Kaiser, S.E., Zuzow, N., Riviere, P., Yang, B., Fenech, E., et al. (2015). The Unfolded Protein Response Triggers Site-Specific Regulatory Ubiquitylation of 40S Ribosomal Proteins. Mol. Cell *59*, 35–49. https://doi.org/10.1016/j.molcel.2015.04.026.

10. Garzia, A., Jafarnejad, S.M., Meyer, C., Chapat, C., Gogakos, T., Morozov, P., Amiri, M., Shapiro, M., Molina, H., Tuschl, T., et al. (2017). The E3 ubiquitin ligase and RNA-binding protein ZNF598 orchestrates ribosome quality control of premature polyadenylated mRNAs. Nat. Commun. *8*, 16056. https://doi.org/10.1038/ncomms16056.

11. Juszkiewicz, S., Chandrasekaran, V., Lin, Z., Kraatz, S., Ramakrishnan, V., and Hegde, R.S. (2018). ZNF598 Is a Quality Control Sensor of Collided Ribosomes. Mol. Cell *72*, 469-481.e7. https://doi.org/10.1016/j.molcel.2018.08.037.

12. Kimball, S.R., Abbas, A., and Jefferson, L.S. (2008). Melatonin represses oxidative stress-induced activation of the MAP kinase and mTOR signaling pathways in H4IIE hepatoma cells through inhibition of Ras. J. Pineal Res. *44*, 379–386. https://doi.org/https://doi.org/10.1111/j.1600-079X.2007.00539.x.

13. Watanabe-Asano, T., Kuma, A., and Mizushima, N. (2014). Cycloheximide inhibits starvation-induced autophagy through mTORC1 activation. Biochem. Biophys. Res. Commun. *445*, 334–339. https://doi.org/https://doi.org/10.1016/j.bbrc.2014.01.180.

14. Goto, Y., Katoh, T., and Suga, H. (2011). Flexizymes for genetic code reprogramming. Nat. Protoc. *6*, 779–790. https://doi.org/10.1038/nprot.2011.331.

15. Josue Ruiz, E., Pinto-Fernandez, A., Turnbull, A.P., Lan, L., Charlton, T.M., Scott, H.C., Damianou, A., Vere, G., Riising, E.M., Da Costa, C., et al. (2021). Usp28 deletion and small-molecule inhibition destabilizes c-myc and elicits regression of squamous cell lung carcinoma. Elife *10*. https://doi.org/10.7554/eLife.71596.

16. Wrigley, J.D., Gavory, G., Simpson, I., Preston, M., Plant, H., Bradley, J., Goeppert, A.U., Rozycka, E., Davies, G., Walsh, J., et al. (2017). Identification and Characterization of Dual Inhibitors of the USP25/28 Deubiquitinating Enzyme Subfamily. ACS Chem. Biol. *12*, 3113–3125. https://doi.org/10.1021/acschembio.7b00334.

17. Chan, W.C., Liu, X., Magin, R.S., Girardi, N.M., Ficarro, S.B., Hu, W., Tarazona Guzman, M.I., Starnbach, C.A., Felix, A., Adelmant, G., et al. (2023). Accelerating inhibitor discovery for deubiquitinating enzymes. Nat. Commun. *14*, 686. https://doi.org/10.1038/s41467-023-36246-0.

18. Varca, A.C., Casalena, D., Chan, W.C., Hu, B., Magin, R.S., Roberts, R.M., Liu, X., Zhu, H., Seo, H.-S., Dhe-Paganon, S., et al. (2021). Identification and validation of selective deubiquitinase inhibitors. Cell Chem. Biol. *28*, 1758-1771.e13. https://doi.org/10.1016/j.chembiol.2021.05.012.

19. Varca, A.C., Casalena, D., Auld, D., and Buhrlage, S.J. (2021). Identification of deubiquitinase inhibitors via high-throughput screening using a fluorogenic ubiquitin-rhodamine assay. STAR Protoc. *2*, 100896. https://doi.org/10.1016/j.xpro.2021.100896.

20. Jones, H.B.L., Draganov, S.D., Huamán, S.S., Wing, P.A.C., Nguyen, C., Liang, Z., Dörner, J., Lithgow, J., Murphy, E., Beard, A., et al. (2025). *ob* ABPP-HT*: A Precision-Engineered Activity Proteomics Pipeline for the Streamlined Discovery of Deubiquitinase Inhibitors https://doi.org/10.1101/2025.05.27.656269.

21. Pinto-Fernandez, A., Salio, M., Partridge, T., Chen, J., Vere, G., Greenwood, H., Olie, C.S., Damianou, A., Scott, H.C., Pegg, H.J., et al. (2021). Deletion of the deISGylating enzyme USP18 enhances tumour cell antigenicity and radiosensitivity. Br. J. Cancer *124*. https://doi.org/10.1038/s41416-020-01167-y.

22. Turnbull, A.P., Ioannidis, S., Krajewski, W.W., Pinto-Fernandez, A., Heride, C., Martin, A.C.L., Tonkin, L.M., Townsend, E.C., Buker, S.M., Lancia, D.R., et al. (2017). Molecular basis of USP7 inhibition by selective small-molecule inhibitors. Nature *550*. https://doi.org/10.1038/nature24451.

23. Pinto-Fernández, A., Davis, S., Schofield, A.B., Scott, H.C., Zhang, P., Salah, E., Mathea, S., Charles, P.D., Damianou, A., Bond, G., et al. (2019). Comprehensive Landscape of Active

Deubiquitinating Enzymes Profiled by Advanced Chemoproteomics. Front. Chem. *7*. https://doi.org/10.3389/fchem.2019.00592.

24. Jones, H.B.L., Heilig, R., Fischer, R., Kessler, B.M., and Pinto-Fernández, A. (2021). ABPP-HT - High-Throughput Activity-Based Profiling of Deubiquitylating Enzyme Inhibitors in a Cellular Context. Front. Chem. *9*. https://doi.org/10.3389/fchem.2021.640105.

25. Jones, H.B.L., Heilig, R., Davis, S., Fischer, R., Kessler, B.M., and Pinto-Fernández, A. (2022). ABPP-HT*—Deep Meets Fast for Activity-Based Profiling of Deubiquitylating Enzymes Using Advanced DIA Mass Spectrometry Methods. Int. J. Mol. Sci. *23*. https://doi.org/10.3390/ijms23063263.

26. Hsu, J.C.-C., Laurent-Rolle, M., Pawlak, J.B., Wilen, C.B., and Cresswell, P. (2021). Translational shutdown and evasion of the innate immune response by SARS-CoV-2 NSP14 protein. Proceedings of the National Academy of Sciences *118*. https://doi.org/10.1073/pnas.2101161118.

27. Hsu, J.C.-C., Pawlak, J.B., Laurent-Rolle, M., and Cresswell, P. (2022). Protocol for assessing translational regulation in mammalian cell lines by OP-Puro labeling. STAR Protoc. *3*, 101654. https://doi.org/10.1016/j.xpro.2022.101654.

28. Goodman, C.A., and Hornberger, T.A. (2013). Measuring Protein Synthesis With SUnSET. Exerc. Sport Sci. Rev. *41*, 107–115. https://doi.org/10.1097/JES.0b013e3182798a95.

29. Ravi, V., Jain, A., Mishra, S., and Sundaresan, N.R. (2020). Measuring Protein Synthesis in Cultured Cells and Mouse Tissues Using the Non-radioactive SUnSET Assay. Curr. Protoc. Mol. Biol. *133*. https://doi.org/10.1002/cpmb.127.

30. Davis, S., Charles, P.D., He, L., Mowlds, P., Kessler, B.M., and Fischer, R. (2017). Expanding Proteome Coverage with CHarge Ordered Parallel Ion aNalysis (CHOPIN) Combined with Broad Specificity Proteolysis. J. Proteome Res. *16*. https://doi.org/10.1021/acs.jproteome.6b00915.

31. Tyanova, S., Temu, T., and Cox, J. (2016). The MaxQuant computational platform for mass spectrometry-based shotgun proteomics. Nat. Protoc. *11*, 2301–2319. https://doi.org/10.1038/nprot.2016.136.

32. Perez-Riverol, Y., Bandla, C., Kundu, D.J., Kamatchinathan, S., Bai, J., Hewapathirana, S., John, N.S., Prakash, A., Walzer, M., Wang, S., et al. (2025). The PRIDE database at 20 years: 2025 update. Nucleic Acids Res. *53*, D543–D553. https://doi.org/10.1093/nar/gkae1011.

33. Li, W., Köster, J., Xu, H., Chen, C.-H., Xiao, T., Liu, J.S., Brown, M., and Liu, X.S. (2015). Quality control, modeling, and visualization of CRISPR screens with MAGeCK-VISPR. Genome Biol. *16*, 281. https://doi.org/10.1186/s13059-015-0843-6.

34. Li, W., Xu, H., Xiao, T., Cong, L., Love, M.I., Zhang, F., Irizarry, R.A., Liu, J.S., Brown, M., and Liu, X.S. (2014). MAGeCK enables robust identification of essential genes from genome-scale CRISPR/Cas9 knockout screens. Genome Biol. *15*, 554. https://doi.org/10.1186/s13059-014-0554-4.

35. McCoy, A.J., Grosse-Kunstleve, R.W., Adams, P.D., Winn, M.D., Storoni, L.C., and Read, R.J. (2007). Phaser crystallographic software. J. Appl. Crystallogr. https://doi.org/10.1107/S0021889807021206.

36. Emsley, P., and Cowtan, K. (2004). Coot: Model-building tools for molecular graphics. Acta Crystallogr. D Biol. Crystallogr. https://doi.org/10.1107/S0907444904019158.

37. Murshudov, G.N., Skubák, P., Lebedev, A.A., Pannu, N.S., Steiner, R.A., Nicholls, R.A., Winn, M.D., Long, F., and Vagin, A.A. (2011). REFMAC5 for the refinement of macromolecular crystal structures. Acta Crystallogr. D Biol. Crystallogr. https://doi.org/10.1107/S0907444911001314.

38. McShane, E., Sin, C., Zauber, H., Wells, J.N., Donnelly, N., Wang, X., Hou, J., Chen, W., Storchova, Z., Marsh, J.A., et al. (2016). Kinetic Analysis of Protein Stability Reveals Age-Dependent Degradation. Cell *167*, 803-815.e21. https://doi.org/10.1016/j.cell.2016.09.015.

39. Goto, Y., Ohta, A., Sako, Y., Yamagishi, Y., Murakami, H., and Suga, H. (2008). Reprogramming the Translation Initiation for the Synthesis of Physiologically Stable Cyclic Peptides. ACS Chem. Biol. *3*, 120–129. https://doi.org/10.1021/cb700233t.

40. Murakami, H., Ohta, A., Ashigai, H., and Suga, H. (2006). A highly flexible tRNA acylation method for non-natural polypeptide synthesis. Nat. Methods *3*, 357–359. https://doi.org/10.1038/nmeth877.

41. Niwa, N., Yamagishi, Y., Murakami, H., and Suga, H. (2009). A flexizyme that selectively charges amino acids activated by a water-friendly leaving group. Bioorg. Med. Chem. Lett. *19*, 3892–3894. https://doi.org/https://doi.org/10.1016/j.bmcl.2009.03.114.
